# A biGWAS strategy reveals the genetic architecture of the interaction between wheat and *Blumeria graminis* f. sp. *tritici*

**DOI:** 10.1101/2025.04.09.647224

**Authors:** Jingzhong Xie, Qiaoling Luo, Limin Wang, Dan Qiu, Caihong Zhao, Jinghuang Hu, Jing Zhang, Xinyu Zhao, Zhaogen Chen, Yibo Wang, Yang Yu, Mengzhen Luo, Haoyuan Song, Yuexuan Hou, Zhimeng Zhang, Mou Yin, Haojie Wang, Xuanzhao Li, Xiaomeng Fu, Bei Xiao, Yahui Li, Jiajie Wu, Wenxuan Liu, Yanpeng Wang, Mo Zhu, Yanming Zhang, Alisdair R. Fernie, Wei Wang, Hongjie Li, Fei He

## Abstract

**Background:** Wheat powdery mildew, caused by *Blumeria graminis* f. sp. *tritici* (*Bgt*) is one of the most significant diseases affecting global wheat production. Breeding for disease resistance (*R*) genes against *Bgt* often follows a ‘boom-bust’ cycles, where cultivars with effective resistance are widely deployed on an expanding area until virulence emerges in the avirulence (*Avr*) pathogen population. While extensive effort has been devoted to identifying and cloning *R* genes, Avr genes remain relatively understudied, limiting our understanding of R-Avr interactions. That said, understanding *R-Avr* interactions is crucial for developing durable resistance strategies.

**Results:** We conducted whole genome sequencing on 245 *Bgt* isolates collected from major wheat-growing regions in China, identifying 120 genetically unique isolates. Each of these unique isolates was inoculated and phenotyped on a diverse panel of wheat germplasm. Through Genome-Wide Association Analysis (GWAS) in both *Bgt* and wheat, we identified 65 *Avr* loci and 251 *R* loci overlapping with nine and eight cloned *Avr* or *R* genes, respectively. On average, each isolate carries eight *Avr* alleles, ranging from one to 17. Little geographical preference for *Avr* alleles or their combinations was observed, suggesting that disease resistance breeding against this pathogen should be coordinated at the national level. The level of resistance level is positively correlated with the number of *R* alleles carried by a wheat line, and the average frequency of an *R* allele is 2% among the wheat panel, indicating the potential for accumulating *R* alleles in breeding programs. We mapped 212 *R-Avr* interactions based on joint GWAS using both plant and pathogen genomes and cross-species epistasis, where a network of interactions was formed between wheat and *Bgt*. These interactions indicate that pyramiding five major *R* loci has the potential to confer resistance to half of the *Avr* loci. As a proof of concept, we successfully verified two previously described *R*-*Avr* pairs using tobacco experiments. Furthermore, we provided molecular validation evidence for three new *Avr* genes, including *Bgt-50651*, *BgtE-5826* and *BgtE-20009*, and two new *R-Avr* interactions. Among them, *Bgt-50651* interacts with *Pm1a*, *Pm2a*, and another unidentified *R* gene located near the *Pm6*/*Pm52* interval.

**Conclusion:** Our study provides a framework for understanding the genetic interaction between plants and pathogens. The discovery of novel *R*/*Avr* loci and their complex interaction network underscores the need to integrate crop and pathogen genetic background, particularly the *R*/*Avr* allele composition, into breeding program design. These findings have significant implications for developing durable resistance strategies in wheat alongside offering valuable insights into the broader dynamics of plant-pathogen interactions.

## Introduction

Plant diseases pose a significant threat to global food security, causing substantial yield losses and food quality deterioration. Up to 20% of the annual wheat yield is lost due to fungal pathogens ^1^. Effectively managing plant pathogens in a sustainable way depends on disease resistance breeding, which follows the so-called ‘boom-bust’ cycles. These cycles involve periods of expanding deployment of resistance genes (the ‘boom’) until virulence arises in the pathogen population (the ‘bust’). One of the key theories explaining this interaction is the ‘Gene-for-Gene hypothesis’ proposed by Harold H. Flor ^2^. This hypothesis suggests that for every resistance (*R*) gene in the host plant, there is a corresponding avirulence (*Avr*) gene in the pathogen. When the *R* gene and the *Avr* gene are compatible, the plant recognizes the pathogen and initiates a defense response, leading to programmed cell death and resistance. However, if the pathogen mutates its *Avr* gene, it can evade detection and cause disease ^3^. The ongoing “arms race” means that dissecting the interaction between *R* and *Avr* is essential for durable disease control in agriculture ^4^.

Wheat powdery mildew, caused by the fungus *Blumeria graminis* f. sp. *tritici* (*Bgt*), is one of the most significant airborne foliar diseases affecting wheat globally. It is characterized by white, powdery fungal growth on leaves, stems, and spikes, leading to reduced grain yield and quality. More than 100 resistance (*R*) genes or alleles against *Bgt*, known as *Pm* genes, have been identified in wheat and its wild relatives ^5^. In parallel, nine *Avr* genes from *Bgt* have been cloned ^3^, namely *AvrPm1a*, *AvrPm1a.2*, *AvrPm2a*, *AvrPm3^a^*^2^*^/f^*^2^, *AvrPm3^b^*^2^*^/c^*^2^, *AvrPm3^d3^*, *AvrPm8*, *AvrPm17* and *AvrPm60* ^6^. Despite this progress, the identification of *Avr* genes and the elucidation of their functional mechanisms considerably trails that of *Pm* genes. Moreover, many *Pm* genes were overcome by virulent isolates shortly after their application in agriculture. For example, the *Pm8*, introgressed from rye (*Secale cereale* L.), lost its immunity against *Bgt* after its widespread deployment due to the standing variation of virulence alleles in pathogen population. Cloning *R* genes and deploying them in breeding programs without considering the corresponding *Avr* gene it recognizes can sometimes be catastrophic, as virulence may exist as standing variation within pathogen populations and their frequencies can be quickly increased under the selection pressure exerted by widespread presence of the *R* genes ^3^.

Increasing evidence suggest that the *R* and *Avr* interactions is often more complex than the simple one-for-one gene interaction model. In the *Leptosphaeria maculans/Brassica napus* pathosystem, a single *Avr* gene can be recognized by multiple *R* genes. For instance, *AvrLm1* is recognized by *Rlm1* and *LepR3*, two *R* genes located on different chromosomes ^7^. Similarly, two *R* genes can also collaborate to recognize a single *Avr* gene. For example, Cesari et al. showed that *AVR-Pia* is recognized by two head-to-head *R* genes, *RGA4* and *RGA5*, both of which are required for resistance against the rice blast fungus ^8^. Similarly, a pair of NLR genes, *RXL*/*Pm5e*, are required to confer resistance to wheat powdery mildew ^9^. Additionally, a single *R* gene can recognize multiple *Avr* genes. For example, *Pm1a* recognizes two different *Avr* genes (*AvrPm1a* and *AvrPm1a.2*) on different *Bgt* chromosomes ^10^. Moreover, epistasis has been observed between *R* genes or between *Avr* genes. When breeding an *R* gene into a wheat cultivar, resistance is not always effectively expressed. This dependency on the genetic background has long been known. For example, a subunit of the mediator complex was found to suppress the stem rust resistance in wheat ^11^. The recognition of *AvrLm3* by *Rlm3* is lost in the presence of *AvrLm4-7*, which itself is recognized by two *R* genes, *Rlm4* and *Rlm7* ^12^. Likewise, in powdery mildew, *SvrPm3* can suppress the recognition of *AvrPm3* by *Pm3* ^13^. Furthermore, homologous *R* genes such as *Pm3* and *Pm8* can form dimer, and thereby loss the ability to recognize pathogen effectors ^14^. These recent advances indicate a complex interaction pattern between *R* and *Avr* that extends beyond the simple ‘one *R* for one *Avr*’ scenario.

Systematic dissection of interaction between plants and pathogens can be achieved through joint association mapping using both plant and pathogen genomes. Such a design successfully mapped gene interactions between *Arabidopsis thaliana* and *Xanthomonas arboricola* ^15^, where a panel of 130 *Arabidopsis* lines were phenotyped against to 20 pathogen strains. A cross-species 2D GWAS was similarly used to study the interaction between 701 rice lines and 23 *X. oryzae* pv. *oryzae* strains, revealing complex genetic interactions between plants and pathogens ^16^. However, such a strategy has never been applied to study either wheat or any critical biotrophic fungal pathogens. Inspired by these studies in *Arabidopsis* and rice, we phenotyped 581 diverse wheat lines using a panel of 120 genetically unique *Bgt* isolates. These wheat lines represent well-known landraces and widely-grown cultivars that have played critical roles in the history of wheat breeding, while the pathogen isolates were isolated from wheat powdery mildew samples collected from major wheat growing regions in China in recent years. We here aimed to address the following questions: (1) Can GWAS be used to identify *R* alleles in the wheat germplasms or *Avr* alleles in the pathogen population? (2) How many *R* or *Avr* genes can be cloned from GWAS results, and are there any new genes? (3) Can the genetic interaction between *R* and *Avr* loci be mapped using two-genome-based approaches? (4) How many *R-Avr* gene interactions can be cloned and molecularly verified, and are there any new interactions? By performing GWAS on the wheat population and the *Bgt* population separately, we detected 251 *R* loci and 65 *Avr* loci, including several previously described genes. Using a cross-species GWAS method developed in this study, we mapped a network of interactions between *R* and *Avr* loci, including many known *R-Avr* interactions. We validated four *Pm* genes (*Pm1a*, *Pm2a*, *Pm4b*, *Pm5b*) and five *Avr* genes (*AvrPm1a*, *AvrPm2a*, *AvrN1*, *AvrN2*, *AvrN16*) from those loci, where three *Avr* genes were reported for the first time. Four *R-Avr* interactions were molecularly verified, including two that are described here for the first time. Our results thus demonstrate the power of cross-species joint association analysis for studying plant-pathogen interactions and revealed the complex genetic architecture of wheat disease resistance against wheat powdery mildew.

## Results

### Assembling a genetically diverse panel of *Bgt* isolates

While wheat germplasm resources are widely exchanged and accessible to researchers, assembling strictly obligate biotrophic pathogen such as powdery mildew is not straightforward. In addition to 123 *Bgt* isolates collected and maintained in our laboratory since 2007, 122 isolates were collected in 2022 with the help of the wheat community. This panel of 245 isolates represents powdery mildew field samples of 88 geographical sites across 19 provinces, covering the major wheat-growing regions of China. (Fig. 1a; Supplementary Table 1). To obtain purified isolate, a single tiny spore colony was randomly selected and inoculated onto fresh wheat leaves in a petri dish (Supplementary Fig. 1a). This process was repeated six to ten times to ensure purification. In total, 245 purified isolates were obtained, and these were subjected to whole-genome sequencing (WGS).

**Figure 1.**
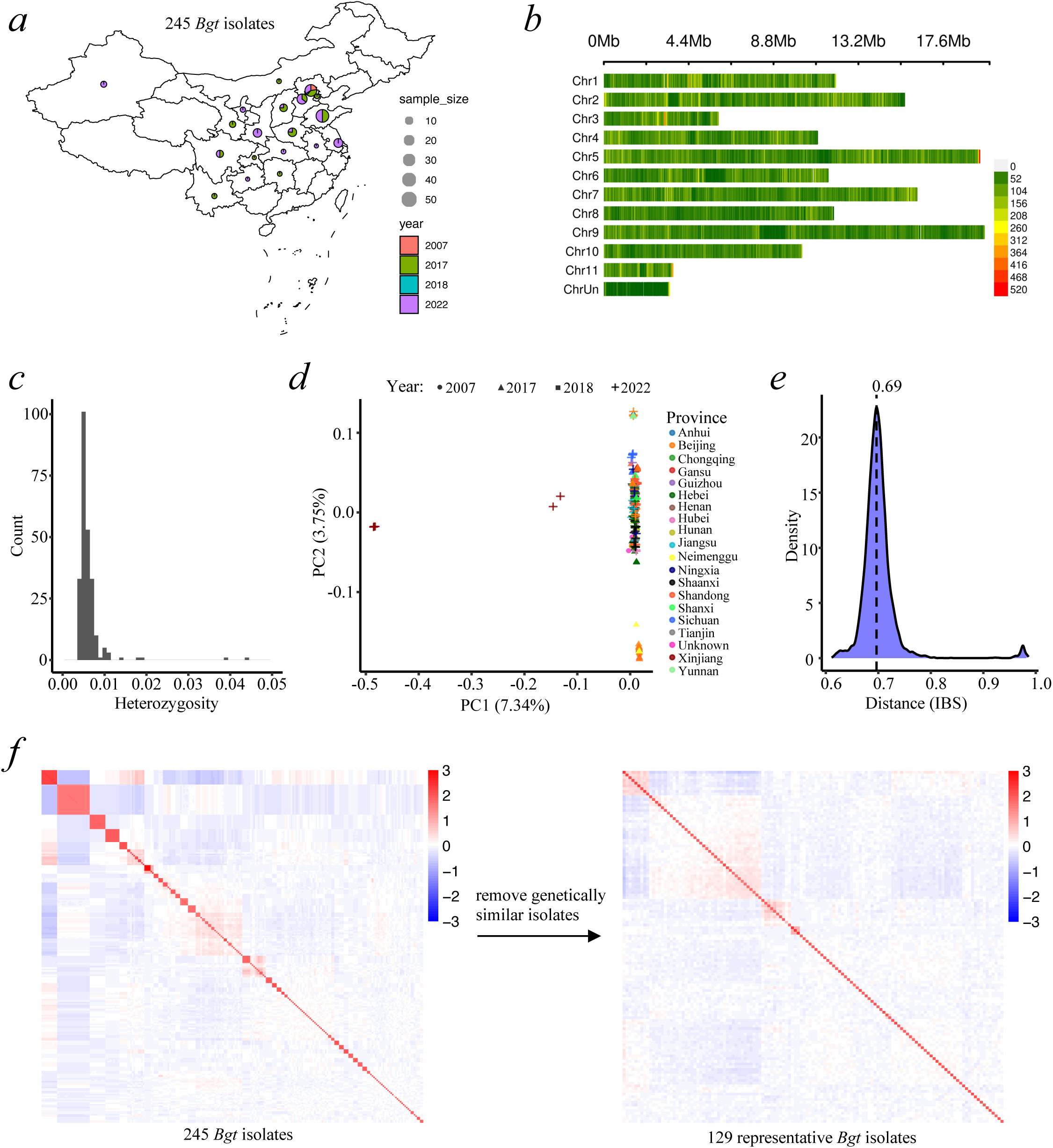
Creating a genetically diverse panel of *B. graminis* f. sp. *tritici* isolates. (a) Geographic origins of the 245 *Bgt* isolates. (b) Genome-wide SNP density distribution for the called 677,500 SNPs. Bars are colored by the number of SNPs within 10 kb window size. (c) Histogram of heterozygosity across all 245 *Bgt* isolates. Heterozygosity is measured as the proportion of heterozygous genotypes in germline variants. (d) PCA of all 245 *Bgt* isolates based on genome-wide SNP variation. The first two principal components (PC1 and PC2) are plotted, representing major axes of genetic variation within the diversity panel. The percentage of explained variation is indicated in parentheses. Each point corresponds to an isolate, with color indicating geographic origin and shape denoting the year of collection. (e) Distribution of pairwise identity-by-state (IBS) values among isolates. A vertical dotted line indicates the most common degree of genetic similarity observed. (f) Heatmap of pairwise genetic relatedness of all 245 isolates (left) and 129 representative isolates (right).

For each isolate, approximately 12 Gb of short-read WGS data were generated, yielding an average genome coverage of 73.96× (Supplementary Fig. 1b; Supplementary Table 1). A total of 93.43% of reads were properly mapped to the reference genome, identifying 677,500 SNPs with a minor allele frequency (MAF) greater than 0.05. The SNP density across the population was approximately one per 208 bp, with most SNPs evenly distributed throughout the genome, except for a hotspot at the end of chromosome 5 (Fig. 1b). On average, each isolate carried 231,767 mutations relative to the reference genome (Supplementary Fig. 1c), and 38.15% of SNPs were located in genic regions (Supplementary Fig. 1d). Additionally, as *Bgt* is airborne and prone to cross-contamination, we assessed the level of purity by leveraging the nature of haploid conidiospores. If heterozygote genotypes were observed for a SNP site, one of the possible reasons is that the sequenced sample contained multiple isolates and that the isolates harbored different alleles at that site. If a sequenced sample contains more than one isolates, it is likely to observe genome-wide heterozygotes. The proportion of heterozygous genotypes in all germline variants was approximately 0.49% per isolate (Fig. 1c). This indicated low contamination levels, with 235 out of the 245 isolates displaying below 1% heterozygosity.

Principal component analysis (PCA) of the population structure showed that many isolates from different provinces are clustered together, except for a few isolates sampled from Xinjiang and some collected in 2017 (Fig. 1d; Supplementary Fig. 1e). The average identity-by-state (IBS) similarity between isolates was 0.69, ranging from 0.62 to nearly 1 (Fig. 1e). The average nucleotide diversity (π) across the genome is 1.48 × 10^-3^ within nonoverlapping 10 kb windows (Supplementary Fig. 1f). Linkage disequilibrium (LD) decayed rapidly in the *Bgt* population, with r^2^ deceasing from 0.74 to 0.29 within 10 kb (Supplementary Fig. 1g). Half of its maximum value occurred at 1.2 kb. This suggests that extensive historical recombination has occurred between divergent *Bgt* lineages. A total of 116 genetically highly related isolates were excluded from the downstream phenotyping experiment based on any of three criteria: (1) genetic relatedness (kinship coefficient > 0.5), (2) identity-by-descent (IBD) > 0.25, and (3) somatic SNP differences > 5,000 in pairwise comparisons among all 245 isolates (Fig. 1f; Supplementary Fig. 1h). This resulted in a diversity panel of 129 genetically distinct isolates representing the recent *Bgt* population in China.

### Construction of a wheat diversity panel with public sequencing data available

To construct a diverse wheat panel with publicly available exome-capture sequencing data ^17, 18^, we combined 262 lines from the Chinese-mini-core wheat germplasm with 319 lines from the USDA wheat germplasm (Supplementary Table 2). These 581 wheat lines were selected due to their genetic diversity and representativeness in breeding (Fig. 2a). While genotype data (i.e., genomic variants) are available for the 319 USDA lines, the 262 Chinese-mini-core lines only had publicly released raw sequencing data but lack genotype data. To obtain genotype data of these lines, we downloaded the raw sequencing data and performed variant calling using the Chinese Spring reference genome ^19^, ensuring consistency with the genotype data for the USDA lines. After merging the two genotype datasets by retaining only common variant sites, we obtained an integrated genotype dataset containing 675,235 variants with a MAF > 0.01 and a missing rate < 0.25 (Supplementary Fig. 2a). The merging process reduced the number of rare alleles, as shown in the MAF distribution plot (Fig. 2b). Consistent with the exome sequencing data, more variants were located in the terminal regions of chromosomes (Fig. 2c).

**Figure 2.**
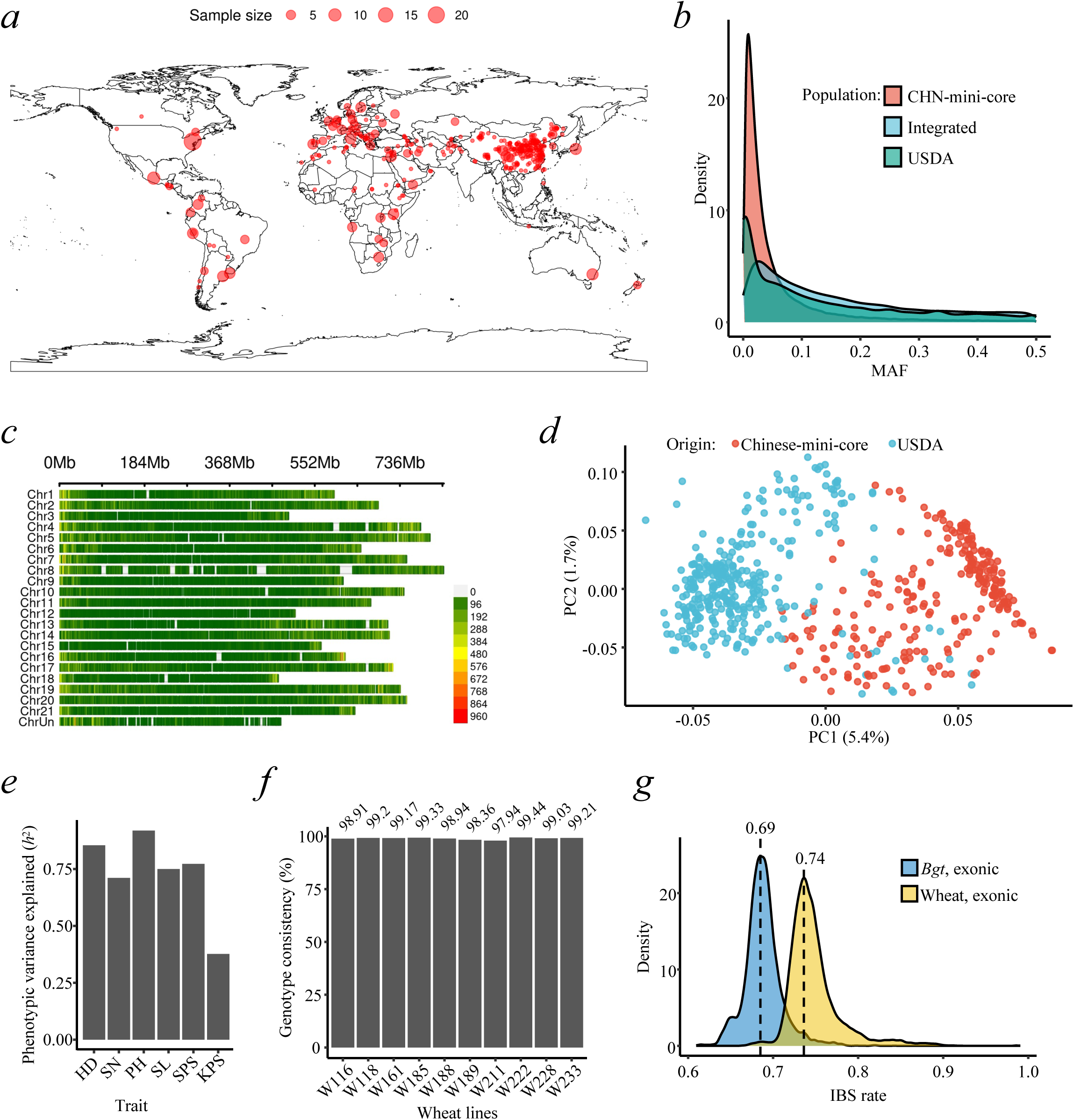
Creating a wheat diversity panel with public exome-capture sequencing data available. (a) Geographic origins of the 581 wheat lines. (b) Minor allele frequency (MAF) distribution in the CHN-mini-core germplasms, USDA germplasms, and the integrated panel. (c) Genome-wide SNP density distribution across chromosomes for the called 675,235 SNPs. Bars are colored by the number of SNPs within a 1 Mb window size. (d) PCA of wheat lines based on integrated genome-wide SNP variation. The first two principal components (PC1 and PC2) are plotted, representing the major axes of genetic variation within the diversity panel. The percentage of explained variation is indicated in parentheses. Each point represents a wheat line, color-coded by its germplasm collection of origin. (e) Narrow-sense heritability (*h*^2^) of 8 traits: HD: heading date, SN: spike number, PH: plant height, SL: spike length, SPS: spikelet number per spike, KPS: kernel number per spike. (f) Genotype consistency between WES-derived SNPs and RNA-seq-derived SNPs in randomly selected 10 wheat lines. (g) Distribution of pairwise identity-by-state (IBS) values at exonic regions within wheat lines or representative *Bgt* isolates.

The USDA and Chinese-mini-core wheat germplasm are largely separated in the population structure PCA plot, with Chinese varieties clustered together (Fig. 2d). This suggests that Chinese wheat germplasm possesses unique genetic characteristics compared to those from other global collections. To validate the reliability of our genotype data, we conducted field trials and calculated the narrow-sense heritability (*h*^2^), which ranged from 0.4 to 0.9 (Fig. 2e). This is comparable to published metrics ^18^, confirming the high reliability of our integrated genotype data. We additionally performed whole-transcriptome sequencing and SNP calling on a subset of randomly selected wheat lines. The genotypic concordance between transcriptome-derived SNPs and SNPs from published data in our integrated dataset are about 99% (Fig. 2f), further confirming the consistency between the published genotype data and the germplasms we obtained. The IBS similarity calculated using exonic SNPs between different wheat lines was generally around 0.74, which is slightly higher than that (0.69) of *Bgt* (Fig. 2g). In summary, we have established a diverse panel of global wheat germplasm for assaying powdery mildew resistance.

### High-throughput phenotyping experiment for wheat powdery mildew

To dissect the genetic mechanism of interaction between wheat and *Bgt*, the seedling-stage disease resistance level were evaluated at the cross-species population scale. A total of 581 wheat lines and 120 of the 129 representative *Bgt* isolates were used, yielding 50,064 infection-type data points (Fig. 3a). To assess the reproducibility of the phenotypic data, we performed repeated phenotyping assays on 23 randomly selected wheat lines across all tested *Bgt* isolates. The correlation between two independent measurements was 0.82 (Fig. 3b). On average, 10% of host-pathogen interactions showed a shift from resistant to susceptible or *vice versa* (Fig. 3b). The average narrow-sense heritability across the 23 traits (i.e., wheat lines) was 90% (Fig. 3b), indicating a strong genetic effect and high reproducibility for the phenotypes. These results demonstrate high quality and reliability of our phenotypic data for studying this fungal disease. Overall, 8.51% of all interactions were classified as resistant (infection types of 0-2). Among the tested wheat lines, 110 exhibited resistances to at least 10% of the 120 *Bgt* isolates, while 193 were highly susceptible to 99% of the isolates (Fig. 3a). These results suggest that resistance carried by wheat lines are isolate-specific and effective resistance gene against powdery mildew has not been widely deployed among the wheat population used here. The proportion of resistant infection types was 11.88% for the USDA germplasm, compared to 4.57% for the Chinese-mini-core lines (Fig. 3c), indicating that the USDA lines likely contain more effective resistance genes against the Chinese *Bgt* population. Additionally, we did not observe 100% virulence or 100% avirulence for any *Bgt* isolates. The variability in virulence levels for all *Bgt* isolates ranged from 74% to 99%, suggesting a highly likelihood for mapping resistance locus against each *Bgt* isolate.

**Figure 3.**
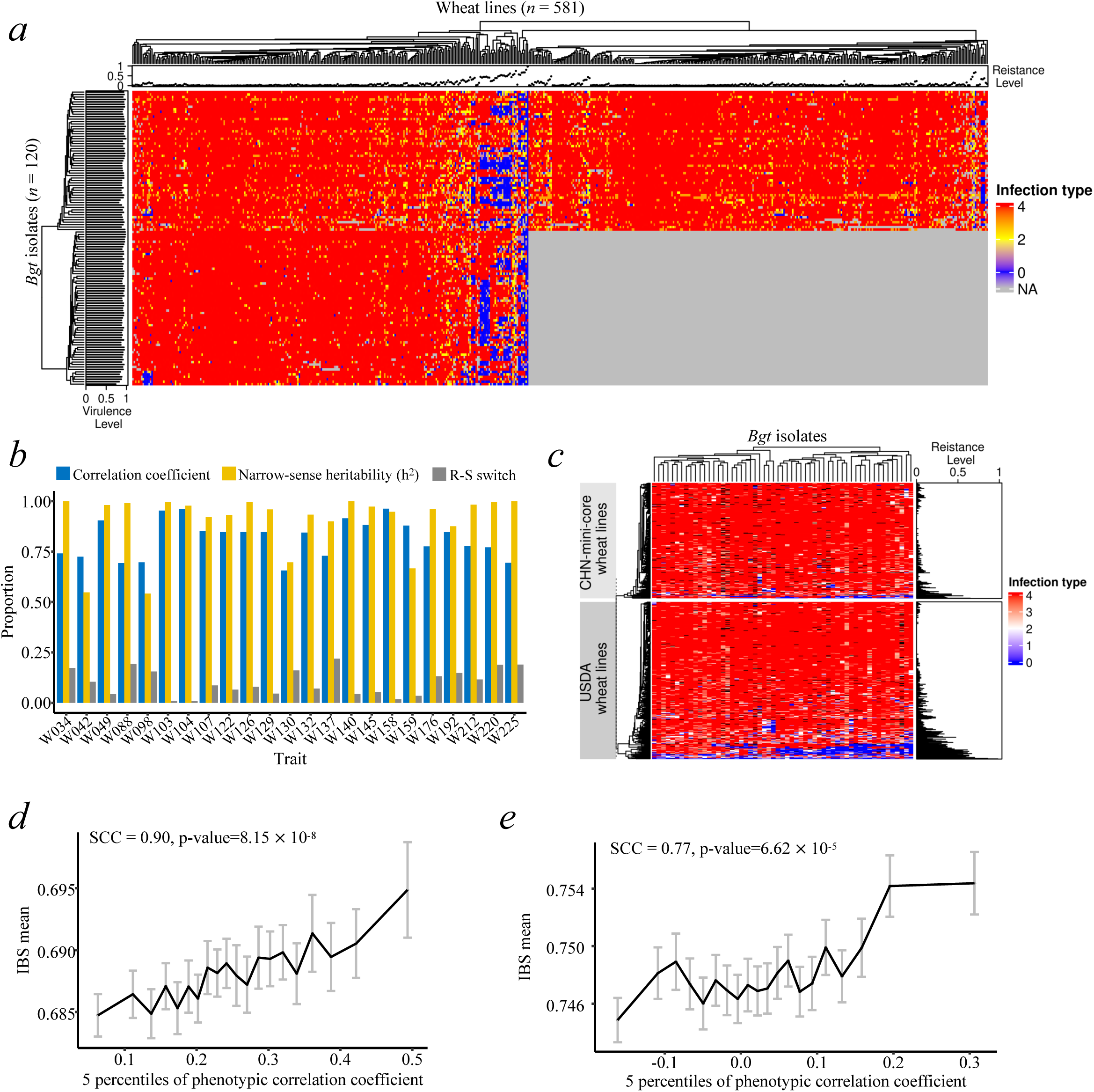
High-throughput phenotyping overview and evaluation of wheat powdery mildew. (a) Graphic infection types for 120 *Bgt* isolates across 581 wheat lines. Grey cells indicate missing phenotypes. The top panel shows the resistance level of each wheat line, calculated as the proportion of avirulent *Bgt* isolates. The left panel shows the virulence level of each *Bgt* isolate, calculated as the proportion of susceptible wheat lines. (b) Reproducibility of infection type scoring across two replicates for 23 wheat lines infected by 116 *Bgt* isolates. The correlation coefficient, Resistant-susceptible switch proportion, and average narrow-sense heritability (*h*²) between replicates are shown. (c) Graphic representation of infection types for 56 *Bgt* isolates across two wheat collections (Chinese-mini-core wheat lines and USDA wheat lines). The other *Bgt* isolates were excluded due to a high rate of missing values. The right panel shows the resistance level of each wheat line, calculated as the proportion of avirulent *Bgt* isolates. (d) Correlation between pairwise IBS value and binned phenotypic correlation coefficient in *Bgt* isolates. The error bar indicates 95% confidence interval. (e) Correlation between pairwise IBS value and binned phenotypic correlation coefficient in wheat lines. The error bar indicates 95% confidence interval.

We compared the genetic similarity (IBS) with the phenotypic similarity (Spearman’s correlation coefficient, SCC) to determine their correlation. Isolates with similar phenotypic spectra tended to share a higher genetic background when analyzed in bins of 5 percentiles (SCC.bin = 0.90, *P* = 8.15 × 10^-8^; Fig. 3d). Similarly, wheat lines with comparable resistance spectra also tended to be genetically similar (SCC.bin = 0.77, *P* = 6.62 × 10^-5^; Fig. 3e), suggesting that genetic variance may contribute to phenotypic variation. The average phenotypic correlation between any two *Bgt* isolates was 0.25, which was significantly higher than the correlation between any two wheat lines (*P* < 2.26 × 10^−16^) (Supplementary Fig. 3a; Supplementary Fig. 3b). This aligns with our population structure analysis, where the wheat population exhibited greater genetic divergence compared to the *Bgt* population.

### The genetic architecture of avirulence in *Bgt* population

We performed GWAS for each of the 581 wheat lines’ phenotypes to map the avirulence (*Avr*) loci on the *Bgt* reference genome. For instance, wheat line W204 was used to assay phenotypic responses against our *Bgt* population. A significant GWAS peak (MAF > 0.05 and Bonferroni-corrected *P* < 1.47 × 10^-8^) was identified at Chr4:237,461-487,461 kb across four statistical models (Supplementary Fig. 4a). This overlapping peak was defined as a high-confidence *Avr* locus, with the most significant variant representing the locus. In total, 576 traits (wheat lines) were successfully analyzed in at least three of the four statistical models, with only five traits failing in a single model (in each instance MLMM). Overall, we identified 65 unique high-confidence *Avr* loci (Fig. 4a; Supplementary Table 3). Among the traits analyzed, 489 did not yield overlapping significant peaks (Supplementary Fig. 4b), primarily because these wheat lines were susceptible to all tested *Bgt* isolates. Meanwhile, 83 traits exhibited a single peak, and nine traits showed more than one peak. These results suggest that resistant wheat lines tend to recognize only one *Avr* locus in the pathogen population.

**Figure 4.**
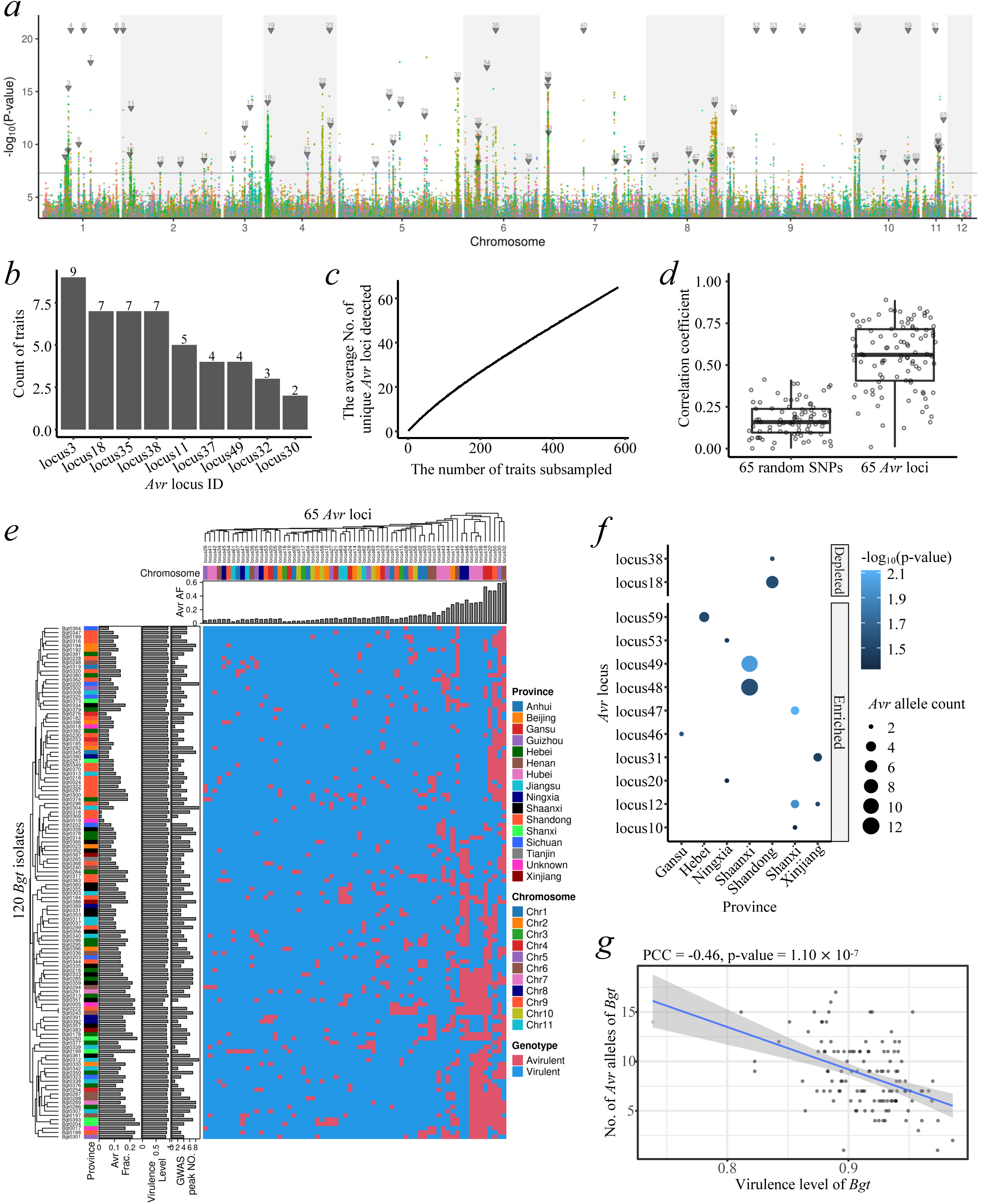
Genomic characterization and distribution of *Avr* loci in the *Bgt* panel. (a) Merged GWAS Manhattan plots for all 581 traits (wheat lines). Different traits are color-coded. Inverted triangles indicate *Avr* loci, with locus IDs labeled above. The solid and dashed horizontal lines represent the Bonferroni - corrected p-value and FDR threshold, respectively. (b) Frequency of traits (wheat lines) in which each *Avr* locus was identified across multiple GWAS analyses. Only loci detected in more than one GWAS run are included. (c) Relationship between the number of sampled traits (wheat lines) and the average number of identified non-redundant *Avr* loci. For each sampling size (1 to 581, step = 1), traits were randomly selected 1000 times, and the mean number of non-redundant *Avr* loci was calculated. (d) gBLUP genomic prediction accuracy comparing the 65 non-redundant *Avr* loci and 65 randomly sampled SNPs (10-fold cross validation). Each test repeated 100 times, and randomly sampled 100 times. (e) Heatmap summarizing the genotypes (virulent and avirulent alleles) of 65 non-redundant *Avr* loci identified through GWAS in the *Bgt* panel. Rows represent *Bgt* isolates, while columns represent *Avr* loci. The upper panel indicates the chromosomal location of each *Avr* locus (color-coded) and the overall frequency of the avirulent allele in the *Bgt* panel. The left panel provides metadata for each *Bgt* isolate, including its geographic origin (province, color-coded), the frequency of avirulent genotypes across the 65 *Avr* loci, the virulence level, and the number of *R* loci identified when the isolate was treated as a trait in GWAS for the wheat panel. ‘AF’: allele frequency; ‘Frac.’: fraction. (f) Over-representation test for *Avr* loci across provinces. Columns represent provinces, and rows represent *Avr* loci. The p-value of the over-representation test for each *Avr* locus–province combination is color-coded, while point size indicates the number of *Bgt* isolates carrying the corresponding *Avr* locus within each province. Significant associations (p-value < 0.05) are shown. (g) Relationship between the virulence level and the number of avirulent alleles across the 65 *Avr* loci for each *Bgt* isolate. A regression curve with a 95% confidence interval, fitted using a generalized linear model, is shown.

Out of the 65 *Avr* loci, nine encompass previously cloned *Avr* genes and 56 are novel (Supplementary Table 3; Supplementary Note2). These cloned *Avr* genes are *AvrPm17* in *Avr.locus04*, *SvrPm3^a1/f1^* in *Avr.locus23*, *AvrPm3^b2/c2^* in *Avr.locus30*, *AvrPm3^a2/f2^* in *Avr.locus33*, *AvrPm1a* in *Avr.locus35*, *AvrPm2a* in *Avr.locus38*, *AvrPm3^d3^* in *Avr.locus50*, *AvrPm1a.2* in *Avr.locus49*, and *AvrPm8* in *Avr.locus64* (Supplementary Note2). Several *Avr* loci were repeatedly detected by at least two wheat lines, including loci overlapping with the cloned *Avr* genes and newly identified loci such as *Avr.locus03*, *11*, *18*, *37*, and *32* (Fig. 4b). Among the identified *Avr* loci, 56 were detected by only a single wheat line. To assess the power of *Avr* locus detection, we simulated the sampling process using varying numbers of wheat lines. As the number of wheat lines used for phenotyping increased, the number of identified *Avr* loci continued to accumulate (Fig. 4c), suggesting that additional *Avr* loci are likely to be discovered as more diverse wheat lines are tested against the *Bgt* population. This indicates that the diversity of resistance genes in wheat remains incompletely explored using our current germplasm collection. Nevertheless, the identification of these 65 *Avr* loci enables us to predict the virulence level of *Bgt* isolates with an accuracy of 0.54, significantly higher than the accuracy of 0.12 achieved using 65 random SNPs (Fig. 4d). This demonstrates the reliability of these *Avr* loci and their value in monitoring the emergence of highly virulent *Bgt* isolates. As expected, candidate effector secreted proteins (CESP) and the biological process of RNA nuclease activity were significantly enriched among the genes surrounding the 65 *Avr* loci, despite the limited functional annotation of *Bgt* proteins (Supplementary Fig. 4c; Supplementary Table 4). This aligns with the previously proposed hypothesis that *Bgt* avirulence genes regulate pathogenicity through RNA metabolism^20^, further supporting the reliability of these identified *Avr* loci. Additionally, protein domains of ribonuclease/ribotoxin-related structures were significantly enriched in the 65 *Avr* loci (Supplementary Table 5), reinforcing the reliability of these identified *Avr* loci.

To understand the population frequency of avirulence alleles in the *Bgt* panel, we generated a virulent-avirulent allelic variation map for the 65 *Avr* loci based on allele-based genetic effects in the GWAS results (Fig. 4e; Supplementary Table 6). The most frequent *Avr* loci carrying an avirulent allele (*Avr* alleles) were detected in more than half of the *Bgt* isolates, including *Avr.locus32*, *30* and *18* (Supplementary Fig. 4d). The *R* genes recognizing these specific loci are likely the most desirable for breeding deployment, as they provide resistance to near half of the pathogen isolates within the population. Seventy seven percent of the *Avr* alleles were found at low frequencies (< 15%) within the *Bgt* population. The average frequency of the 65 *Avr* alleles was 13.28%, ranging from 3% to 59%. Population structure analysis of the presence-absence variation of the *Avr* alleles showed no clear divergence among isolates from different geographical regions (Supplementary Fig. 4e), suggesting that *Avr* allele combinations are likely random. On average, each isolate carried 8.63 *Avr* alleles, with frequencies ranging from 2% to 26%. Notably, each *Bgt* isolate possessed at least one *Avr* allele, implying that *Avr* alleles might play a critical role in the life cycle of *Bgt*. Both frequency and geographical distribution of avirulence genes are critical factors for deploying disease resistance genes in local wheat breeding programs. The presence of approximately eight avirulence alleles per *Bgt* isolate highlights the effectiveness of breeding for disease resistance as a strategy to combat this fungal pathogen. We examined the presence of the *Avr* alleles overlapping with previously cloned *Avr* genes in the 120 *Bgt* isolates (Supplementary Note2, Supplementary Table 3). The *Avr* allele of *Avr.locus04* (*AvrPm17*) was found in only 4% of the *Bgt* isolates, while the remaining *Avr* alleles had frequencies ranging from 7% to 58% in the pathogen population (Supplementary Fig. 4f). Geographical biases were observed for certain *Avr* alleles: *Avr.locus38* (*AvrPm2a*) and *Avr.locus18* were rare in Shandong, while *Avr.locus59* was overrepresented in Hebei. Additionally, *Avr.locus12, 21, 31, 46, 47, 48, 49,* and *53* were enriched in northwestern provinces, including Xinjiang, Ningxia, Shaanxi, Gansu, and Shanxi (Fig. 4f), suggesting that some *Avr* loci are under selective pressure, probably due to the expanded deployment of the corresponding *R* genes. In fact, *Pm2a* was present in cultivar Jimai22, a leading wheat cultivar used in breeding and production in Shandong and other provinces in northern China ^21^. Collectively, the high prevalence of virulence alleles in the *Bgt* population, along with the regional enrichment of certain virulence alleles, underscores the need for careful consideration when deploying *R* genes at both global and local scales for controlling wheat powdery mildew.

The scale of our phenotyping assay allowed us to define the virulence level of a *Bgt* isolate as a quantitative trait, which was measured as the proportion of tested wheat lines to which a *Bgt* isolate is virulent. Using the 65 identified *Avr* loci, we examined the relationship between virulence level and the number of *Avr* alleles present in *Bgt* isolates and found a strong negative correlation (Pearson’s correlation coefficient, PCC=−0.46) (Fig. 4g). This suggests that virulence level in *Bgt* is a quantitative trait primarily determined by the number of *Avr* alleles rather than being controlled by a few large-effect loci.

### The genetic architecture of resistance in a global collection of wheat varieties

Unlike previous GWAS studies that investigated disease resistance to powdery mildew using a few isolates^22^, our experimental design allows us to understand seedling-stage resistance for each of the 120 diverse isolates. This approach provides insight into how different genes are utilized by the plant to combat the pathogen. For each isolate used for phenotyping, GWAS was conducted to map resistance (*R*) loci on the wheat genome. For instance, screening isolate Bgt0211 against our wheat panel identified an overlapping significant GWAS peak at chr5D:43,407,990-43,412,600 in four statistical models, indicating the presence of an *R* locus for this isolate (Supplementary Fig. 5a). Across 120 isolates (i.e., traits), we conducted 480 GWAS analyses using four models, considering only significant peaks detected in at least three statistical models as high-confidence *R* loci. In total, 545 *R* loci were identified from the 4,708 significant GWAS peaks (Supplementary Fig. 5b), with an average of 4.54 *R* loci per isolate (range: 0-9). Only two isolates lacked any *R* loci, while 95% of the 120 isolates had at least two *R* loci, suggesting that wheat employs a diverse arsenal of *R* genes to combat *Bgt* isolates with varying virulence profiles. Isolates Bgt0200 (originated from Sichuan), Bgt0304 (Jiangsu), and Bgt0312 (Jiangsu) had nine *R* loci identified in the wheat germplasm that could be used for resistance breeding (Supplementary Table 1). By contrast, isolates Bgt0298, Bgt0318, and Bgt0322, sampled from Shandong, had very few responding *R* loci, indicating potential challenges in breeding for resistance against these isolates (Supplementary Table 1).

We consolidated multiple nearby *R* loci across traits into a single *R* locus, resulting in 251 non-redundant *R* loci on the wheat genome (Fig. 5a; Supplementary Table 7; See Methods). Simulations revealed that increasing the number of traits leads to the discovery of more *R* loci (Fig. 5b). This suggests that additional *R* loci could be identified in a more diversified wheat germplasm population. Among the 251 *R* loci, 35 were repeatedly identified in at least three traits (Fig. 5c). The most frequently detected *R* loci correspond to *R.locus222* overlapped with *Pm1a* and *R.locus168* overlapped with *Pm2a*, identified in 48 and 35 traits, respectively. These are the first two named *Pm* genes and their frequent appearance in our GWAS peaks is likely due to their frequent breeding deployment rather than coincidence. Additionally, *R* loci at the distal end of chromosome 2B, including *R.locus58* and *R.locus60*, were detected in 43 and 23 traits, respectively, overlapping with the mapping regions of *Pm6* ^23^and *Pm52* ^24^. Other cloned *R* genes, including *Pm4* (*R.locus41*) and *Pm5* (*R.locus232*) were also detected in multiple traits (eight each). Moreover, 86% of *R* loci were identified in less than three traits, suggesting that wheat employs diverse *R* genes or alleles to recognize different *Bgt* isolates. Only two of the 251 *R* loci exhibited positive (susceptible) genetic effects (Supplementary Table 7), implying that most alternative alleles confer resistance. This aligns with the observation that the Chinese Spring reference genome carries susceptible alleles and is universally susceptible to all tested *Bgt* isolates. Using genotypes of the 251 *R* loci, we achieved a prediction accuracy of 0.65 for wheat resistance level, significantly higher than the 0.22 accuracy obtained using 251 random SNPs (Fig. 5d). This highlights the reliability of the identified *R* loci and their potential for practical applications in resistance breeding.

**Figure 5.**
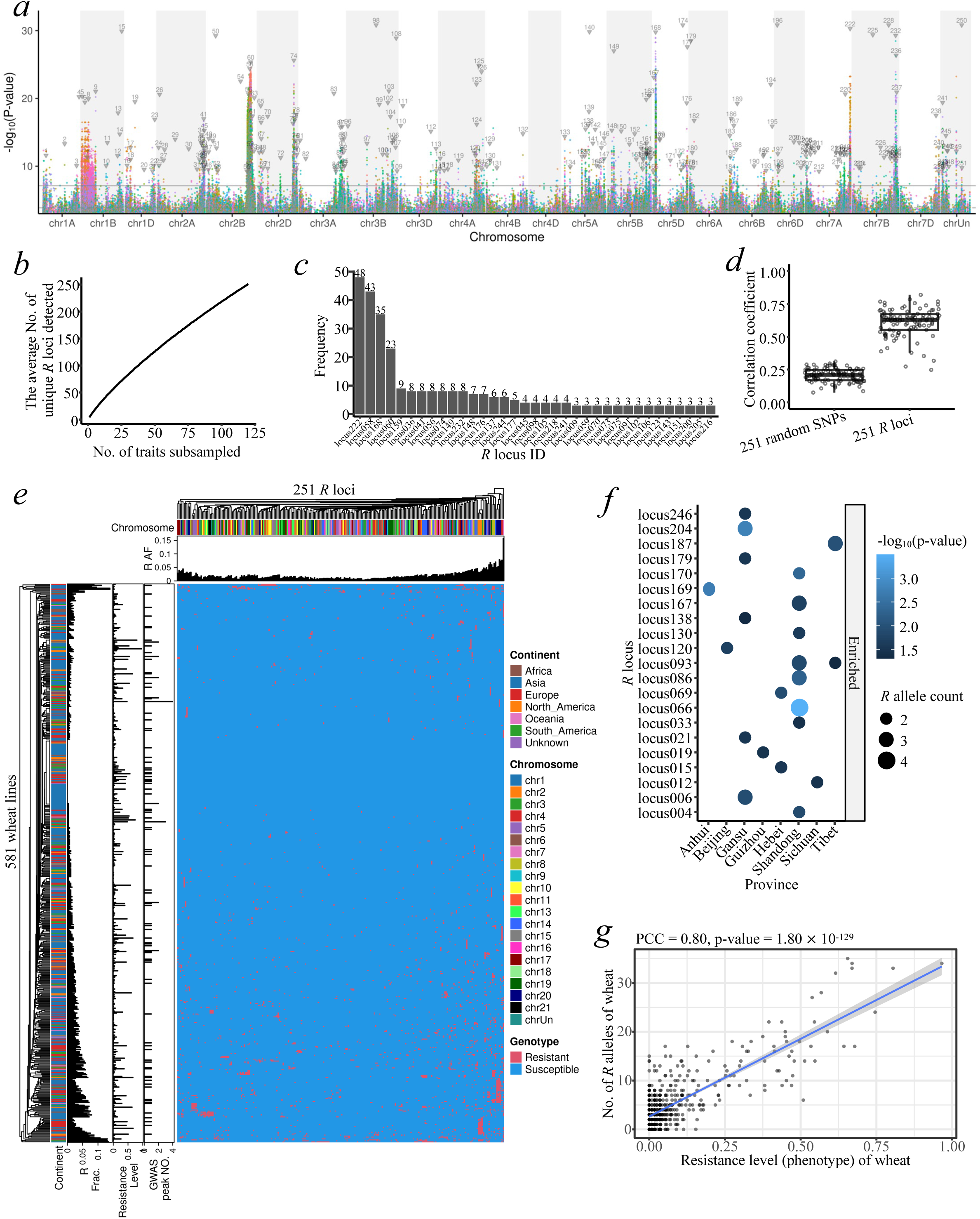
Genomic characterization and distribution of *R* loci in the wheat panel. (a) Merged GWAS Manhattan plots for all 120 traits (*Bgt* isolates). Different traits are color-coded. Inverted triangles indicate *R* loci, with locus IDs labeled above. The solid and dashed horizontal lines represent the Bonferroni-corrected p-value and FDR threshold, respectively. (b) Relationship between the number of sampled traits (*Bgt* isolates) and the average number of identified non-redundant *R* loci. For each sampling size (1 to 120, step = 1), traits were randomly selected 1000 times, and the average number of non-redundant *R* loci was calculated. (c) Frequency of traits (*Bgt* isolates) in which each *R* locus was identified across multiple GWAS analyses. Loci with frequency >=3 are shown. (d) gBLUP genomic prediction accuracy comparing the 251 non-redundant *R* loci and 251 randomly sampled SNPs (10-fold cross validation). Each test repeated 100 times, and randomly sampled 100 times. (e) Heatmap summarizing the genotypes (resistant and susceptible alleles) of the 251 non-redundant *R* loci identified through GWAS in the wheat panel. Rows represent wheat lines, while columns represent *R* loci. The upper panel indicates the chromosomal origin of each *R* locus (color-coded) and the overall frequency of the resistant allele in the wheat panel. The left panel provides metadata for each wheat line, including its geographic origin (continent, color-coded), the frequency of resistant genotypes across the 251 non-redundant *R* loci, the resistance level, and the number of *Avr* loci identified when the wheat line was treated as a trait in GWAS for the *Bgt* panel. ‘AF’: allele frequency; ‘Frac.’: fraction. (f) Over-representation test for *R* loci across provinces. Columns represent provinces of China, and rows represent *R* loci. The p-value of the over-representation test of each *R* locus-province combination is color-coded. (g) Relationship between the resistance level and the number of resistant alleles across the 251 *R* loci for each wheat line. A regression curve with a 95% confidence interval, fitted using a generalized linear model, is shown.

Genes flanking the identified *R* loci are significantly enriched for biological processes related to defense response and oxidative stress, as revealed by GO enrichment analysis (Supplementary Fig. 5c). Additionally, domain enrichment analysis shows an overrepresentation of protein domains commonly found in disease resistance proteins, including NB-ARC, LRR, and Zinc fingers (Supplementary Fig. 5d). These results support the validity of the *R* loci identified through GWAS.

We constructed a resistant-susceptible allelic variation map for the 251 *R* loci (Fig. 5e; Supplementary Table 8). The frequency of resistant alleles (*R* alleles) is on average 2% in the wheat population, ranging from 1% to 15% for different *R* loci. Among them, *R.locus86* and *R.locus232* (overlapped with *Pm5*) are the most frequently detected *R* alleles in the wheat population, yet they are only present in 15% and 9% of wheat lines, respectively. On average, a wheat line carries five *R* alleles, with the number ranging from 0 to 35. Notably, 68 wheat lines carry none of the 251 *R* alleles, and 24% of the wheat lines possess at most one *R* allele. This suggests that most *R* alleles tend to be rare variants and have not been widely deployed in breeding programs. This trend is even more evident when examining the frequency spectrum of the eight *R* loci with cloned *Pm* alleles (Supplementary Fig. 5e, Supplementary Note2), where 99% of the wheat lines carry no more than one of these alleles. Furthermore, the combination of *R* alleles in the wheat panel appears to follow no clear pattern, as indicated by a single cluster in the results of PCA analysis (Supplementary Fig. 5f). This highlights the potential of pyramiding different *R* loci to enhance disease resistance in wheat breeding programs. Interestingly, *R* allele frequencies are significantly higher in the USDA germplasm than in the Chinese-mini-core germplasm (Supplementary Fig. 6a), with the lowest frequencies observed in wheat lines originating from China (Supplementary Fig. 6b-d). This emphasizes the importance of germplasm exchange in acquiring novel disease resistance genes, which could be particularly beneficial for improving resistance in Chinese wheat varieties. Another alternative explanation is that the Chinese pathogen population has not been subjected to selection pressures by the USDA wheat germplasms.

Enrichment analysis of *R* alleles across provinces reveals that certain *R* alleles are not randomly distributed geographically (Fig. 5f). For instance, *R* alleles at *R.locus66* are enriched in Shandong, *R.locus204* in Gansu, and *R.locus169* in Anhui. This pattern suggests that these *R* alleles may have been subjected to artificial selection in breeding programs or nature selection to enhance adaptation to local ecosystems.

The resistance level of a wheat line was measured as the proportion of *Bgt* isolates to which the wheat line is resistant. With the identification of 251 *R* loci, we found a strong positive correlation (PCC = 0.80) between resistance level and the number of *R* alleles present in wheat lines (Fig. 5g). Additionally, the correlation between resistance level and the accumulated genetic effect of *R* alleles was significant (PCC=−0.78, Supplementary Fig. 6e), whereas the average genetic effect of *R* alleles is not well correlated with the resistance level (Supplementary Fig. 6f). These results indicate that resistance to *Bgt* is primarily determined by the number of *R* alleles present in individual wheat lines in this study.

In summary, we performed GWAS in the host population to identify *R* loci associated with disease resistance phenotypes against each *Bgt* isolate, and GWAS in the pathogen population to map *Avr* loci linked to avirulence phenotypes on individual wheat lines. These analyses identified 251 *R* loci in wheat and 65 *Avr* loci in *Bgt*, providing a comprehensive genetic framework for understanding host-pathogen interactions.

### Mapping *R-Avr* interactions based on a cross-species strategy

To investigate the interactions between the *R* loci and *Avr* loci identified above, we developed a bi-directional GWAS (biGWAS) approach, leveraging differentially enriched locus (DEL) analysis between phenotypically contrasting haplotype groups (Supplementary Fig. 6a-c). This method aimed to identify potential *R-Avr* interacting pairs through two complementary analyses: forward-DEL and reverse-DEL. In the forward-DEL analysis, wheat lines were categorized based on the hapotypes of a given R locus (e.g. R1) into groups R1-R carrying the resistant haplotype (R1^hap-R^) and group R1-S carrying the susceptible haplotype (R1^hap-S^) (Supplementary Fig. 6b). If R1 recognizes a specific *Avr* locus (*Avr^R1^*), the corresponding *Avr* locus should be repeatedly identified via GWAS in the *Bgt* population for wheat lines in the R1-R group, but not in the R1-S group. *Avr* loci differentially enriched between these two groups were identified as potential cognate *Avr* loci (*Avr^R1^*) for *R1* (Supplementary Fig. 6c). Conversely, in the reverse-DEL analysis, *Bgt* isolates were classified based on the haplotypes of a given *Avr* locus (e.g., *A1*) into A1-V (virulent) group and A1-A (avirulent) group. *R* loci differentially enriched between these two groups were identified as potential cognate *R* loci (*R^A1^*) for *A1*. By integrating the results from both forward-DEL and reverse-DEL analyses, we identified bi-directional *R-Avr* interacting pairs. Specifically, if the *Avr* locus *A1* matched *Avr^R1^* from the forward-DEL analysis and the corresponding *R* locus *R1* matched *R^A1^* from the reverse-DEL analysis, *R1* and *A1* were considered a bi-directional *R-Avr* interacting pair. Examples of biGWAS are illustrated in Supplementary Figure 8-11.

The forward-DEL analysis identified 216 *R* loci corresponding to 47 *Avr* loci (Supplementary Fig. 12a; Supplementary Table 9). This analysis successfully recovered many known *R-Avr* interactions, including R1 (i.e., *R.locus001*, covering *Pm3*)-Avr33 (i.e. *Avr.locus33*, *AvrPm3^a2^*^/*f2*^), R168 (*Pm2a*)-Avr38 (*AvrPm2a*), and R222 (*Pm1a*)-Avr35 (*AvrPm1a*). Similarly, the reverse-DEL analysis identified 41 *Avr* loci corresponding to 89 *R* loci (Supplementary Fig. 12b; Supplementary Table 9), recovering interactions like Avr35 (*AvrPm1a*)-R222 (*Pm1a*), Avr38 (*AvrPm2a*)-R168 (*Pm2a*), and Avr64 (*AvrPm8*)-R5 (*Pm8*). Integrating both analyses, we identified 66 bi-directional *R-Avr* interacting pairs (bi-pairs), involving 39 *R* loci and 26 *Avr* loci (Fig. 6a; Supplementary Table 9). Fourteen out of 39 *R* loci interacted with at least two *Avr* loci, and 15 out of 26 *Avr* loci were interacted with multiple *R* loci. Only a minority (7 of the 66 bi-pairs) followed a one-to-one interaction pattern, suggesting a multiple-to-multiple recognition pattern, where individual *R* locus recognize multiple *Avr* loci, and *vice versa*. The genomic distribution of these interacting loci revealed an uneven arrangement of *R* loci along wheat chromosomes, whereas *Avr* loci were more evenly positioned. Several well-characterized *R-Avr* pairs, such as R222 (*Pm1a*)-Avr35 (*AvrPm1a*), R168 (*Pm2a*)-Avr38 (*AvrPm2a*), and R5 (*Pm8*)-Avr64 (*AvrPm8*), were identified as bi-pairs. Beyond these known interactions, our analysis mapped novel *R-Avr* interacting pairs, where known *R* or *Avr* genes interacted with previously uncharacterized loci. These included R168 (*Pm2a*)-Avr18, R232 (*Pm5b*)-Avr3, R58/60-Avr38 (*AvrPm2a*), R36-Avr35 (*AvrPm1a*), R36-Avr33 (*AvrPm3^a2^*^/*f2*^), R222 (*Pm1a*)-Avr48, R176-Avr30 (*AvrPm3^b2^*^/*c2*^), and R41 (*Pm4*)-Avr49 (*AvrPm1a*.*2*). Additionally, several novel *R-Avr* bi-pairs were identified, such as R58/60-Avr18, R73/74-Avr18, R148-Avr18, R58/60-Avr53, R169-Avr3, R137-Avr48, and R5-Avr24. Among them, R58/60 and Avr3 emerged as super loci, interacting with multiple *Avr* and *R* loci, respectively. This suggests their potential significance in wheat disease resistance and breeding programs. These results reveal a complex, likely multiple-to-multiple genetic interaction network underlying the immune response of wheat against *Bgt*.

**Figure 6.**
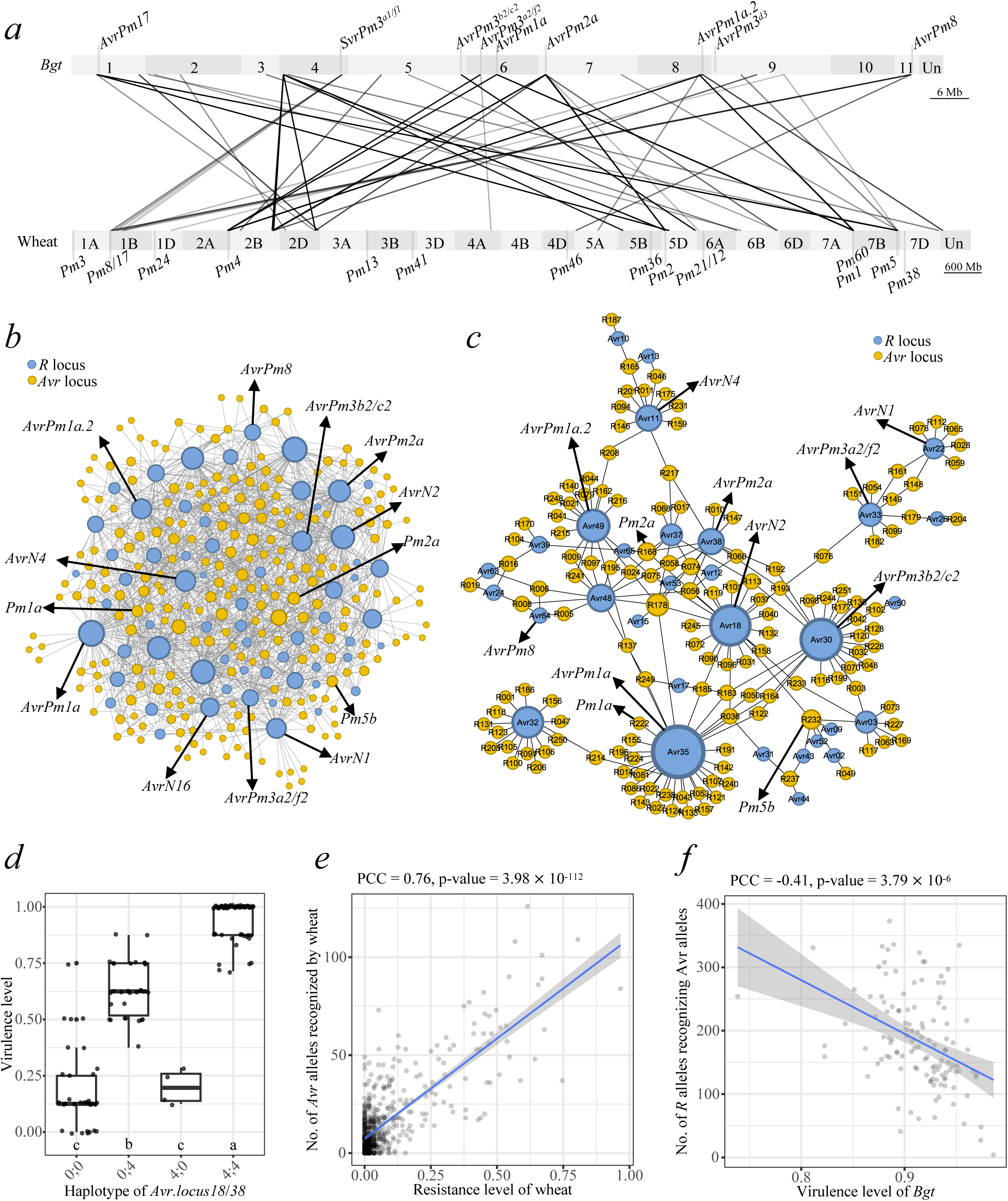
Mining *R-Avr* interacting pairs based on cross-species bi-directional GWAS and epistasis. (a) *R-Avr* recognition pairs identified by biGWAS. Each grey rectangle represents a chromosome, with ID labeled in the center. The *Bgt* chromosomes are scaled 100× for comparison with wheat. Lines connect *R* loci on wheat chromosomes to corresponding *Avr* loci on *Bgt* chromosomes. Previously cloned *Pm* and *AvrPm* genes are labeled at their respective physical positions. (b) *R-Avr* recognition pairs identified in biEpi map. Each node represents an *R* locus (blue) in the wheat genome or an *Avr* locus (orange) in the *Bgt* genome. Node size corresponds to the number of recognition connections (edges) associated with each locus. (c) The 212 *R-Avr* pairs commonly identified by biGWAS and biEpi, with *Avr* loci shown in orange nodes and *R* loci in light blue nodes. Locus ID is abbreviated within the node. For instance, *R.locus123* is abbreviated to R123, and *Avr.locus50* is abbreviated to Avr50. (d) Virulence level comparison of haplotype combinations for the two *Avr* loci (*Avr.locus18*/*38*) recognized by *Pm2a* across all 120 tested *Bgt* isolates. The box edges represent the first and third quartiles, with whiskers extending 1.5 times then interquartile range. Different letters indicate statistically significant differences (*P* < 0.05) based on one-way ANOVA test. ‘0’: avirulent haplotype; ‘4’: virulent haplotype. (e) Relationship between wheat resistance level and the number of *Avr* alleles recognized by the wheat. A regression curve based on a generalized linear model is displayed, with the 95% confidence interval shaded. The correlation coefficient (PCC) and p-value are shown at the top of the plot. (f) Relationship between *Bgt* virulence level and the number of *R* alleles recognizing *Bgt*. A regression curve based on a generalized linear model is displayed, with the 95% confidence interval shaded. The correlation coefficient (PCC) and p-value are shown at the top of the plot.

We constructed a cross-species epistasis interaction map (biEpi map) to complement the *R-Avr* interacting pairs detected in biGWAS. This analysis identified 1,409 significantly associated *R-Avr* pairs within the biEpi map (Fig. 6b; Supplementary Table 10). Notably, 42% (212) of the *R-Avr* pairs identified in biGWAS were also present in the biEpi map, representing a significant overlap (*P* = 1.96**×**10^-60^, Supplementary Fig. 13a). These overlapping pairs retained the multiple-to-multiple pattern (Fig. 6c), further reinforcing the complexity of *R-Avr* interactions in wheat immunity. As a representative example, the predicted R222 (*Pm1a*)-Avr35 (*AvrPm1a*) interaction from biGWAS and biEpi were examined. The infection type was significantly lower only when the resistant haplotype (*Pm1a*^hapR^) was paired with the avirulent haplotype (*AvrPm1a*^hapAvr^), demonstrating a defense response likely triggered by the recognition of *AvrPm1a*^hapAvr^ by *Pm1a*^hapR^ (Supplementary Fig. 13b).

Among the common *R-Avr* pairs identified in both the biGWAS and biEpi, certain *R* and *Avr* loci emerged as interaction hubs, consistently implying the presence of super *R* and *Avr* loci that play a central role in plant-pathogen interactions (Fig. 6c). To confirm this, we investigated the virulence level for different allelic combinations for two *Avr* loci, Avr18 and Avr38 (*AvrPm2a*), which were the two *Avr* loci predicted to be recognized by *Pm2a* (*R.locus168*). Using wheat lines that exclusively carried the resistance allele of *Pm2a* (*R.locus168*), we observed that the virulence level is the lowest when both alleles were present and the highest when both *Avr* were absent (Fig. 6d). The function of Avr18 likely depends on the presence of Avr38 (*AvrPm2a*), indicating epistasis between the two alleles. Similar patterns were observed for other putative super *R*/*Avr* loci (Supplementary Fig. 13c-d). Given these results, we hypothesized that the number of *Avr* loci a wheat line can recognize is associated with its resistance level. Consistent with this, we observed a strong positive correlation (PCC=0.76) between resistance level and the number of detectable *Avr* loci by *R* allele in wheat at the population level (Fig. 6e). Conversely, in *Bgt*, the virulence level was negatively correlated (PCC=−0.41) with the number of corresponding recognizable *R* loci (Fig. 6f). These results suggest that broad-spectrum resistance is likely associated with the number of *Avr* genes a wheat line can recognize. Considering that most *R* loci identified in biGWAS were loci with similar scale of genetic effect (Supplementary Table 7), these results further support the strategy of pyramiding multiple *R* loci to enhance wheat resistance against *Bgt* ^25^.

These *R-Avr* interactions enable us to infer the minimal number of *R* loci needed for recognizing most of the *Avr* loci. We performed the analysis via random-ranking summary found that the top five *R* loci with the broadest *Avr* recognition capacity could recognize half of the *Avr* loci while gathering the top 15 *R* loci could recognize all identified *Avr* loci recognizable (Supplementary Fig. 13e). This study thus paves the way for deploying resistant cultivars more efficiently and precisely.

### Molecular validation of candidate genes in *R*/*Avr* loci and *R-Avr* interacting pairs

To validate the identified *R*/*Avr* loci and their interactions, we selected four newly mapped *Avr* loci for molecular validation, including *Avr.locus18* (*AvrN2*), a locus detected in seven traits, and three rare loci identified in only one trait each: *Avr.locus56* (*AvrN16*), *Avr.locus22* (*AvrN1*) and *Avr.locus11* (*AvrN4*). Although the above works identified the *R* loci recognizing some of the *Avr* loci, determining *R* gene candidate within a locus is more challenging due to complex LD and frequent introgression from distant relatives^18^. Therefore, we utilized the wheat protoplast system and BSMV to experimentally validate the *Avr* candidates. These systems allowed us to validate the molecular function of *Avr* gene without knowing the *R* gene carried by the wheat variety.

The *Avr.locus18* was mapped via GWAS to a ∼250 kb region on *Bgt* chromosome 4 (Fig. 7a; Supplementary Fig. 7a). Based on gene annotation within this interval, we selected *Bgt-50651*, a typical CESP gene designated as *AvrN2*, as the best candidate gene of the *Avr.locus18* (Supplementary Fig. 14a-b). To test its molecular function, an avirulent haplotype of *AvrN2* was cloned and transferred into the cognate resistant wheat line W130 (Supplementary Table 12). Overexpression of this avirulent haplotype in W130 protoplasts resulted in a significant reduction of fluorescence, indicating cell death activation^26^ (Fig. 7b; Supplementary Fig. 14c; Supplementary Table 14). This provides molecular evidence supporting that *AvrN2* likely interacts with a corresponding *R* gene in W130. To further confirm the function of *AvrN2*, BSMV-induced gene silence experiments were performed on W130 leaves. Plants inoculated with BSMV-γ displayed typical stripe mosaic virus symptoms, whereas those infected with BSMV-γ-TaPDS exhibited pronounced leaf whitening (Fig. 7c), demonstrating the successful establishment of the experiment. Plants infected with BSMV-γ-AVRN2 showed suppressed viral symptoms (Fig. 7c), and the expression of *AvrN2* was confirmed in these plants through qRT-PCR (Supplementary Fig. 14d). Expression of *AvrN2* in susceptible control wheat cultivar Zhongzuo 9504 followed by a similar BSMV experiment induced trailing cell death both locally and systemically (Supplementary Fig. 14e). These results confirmed that the *R* protein in wheat line W130 recognizes *AvrN2*, thereby triggering an immune response. Collectively, these results confirmed that *AvrN2* (*Bgt-50651*) functions as an avirulence gene of *Avr.locus18*.

**Figure 7.**
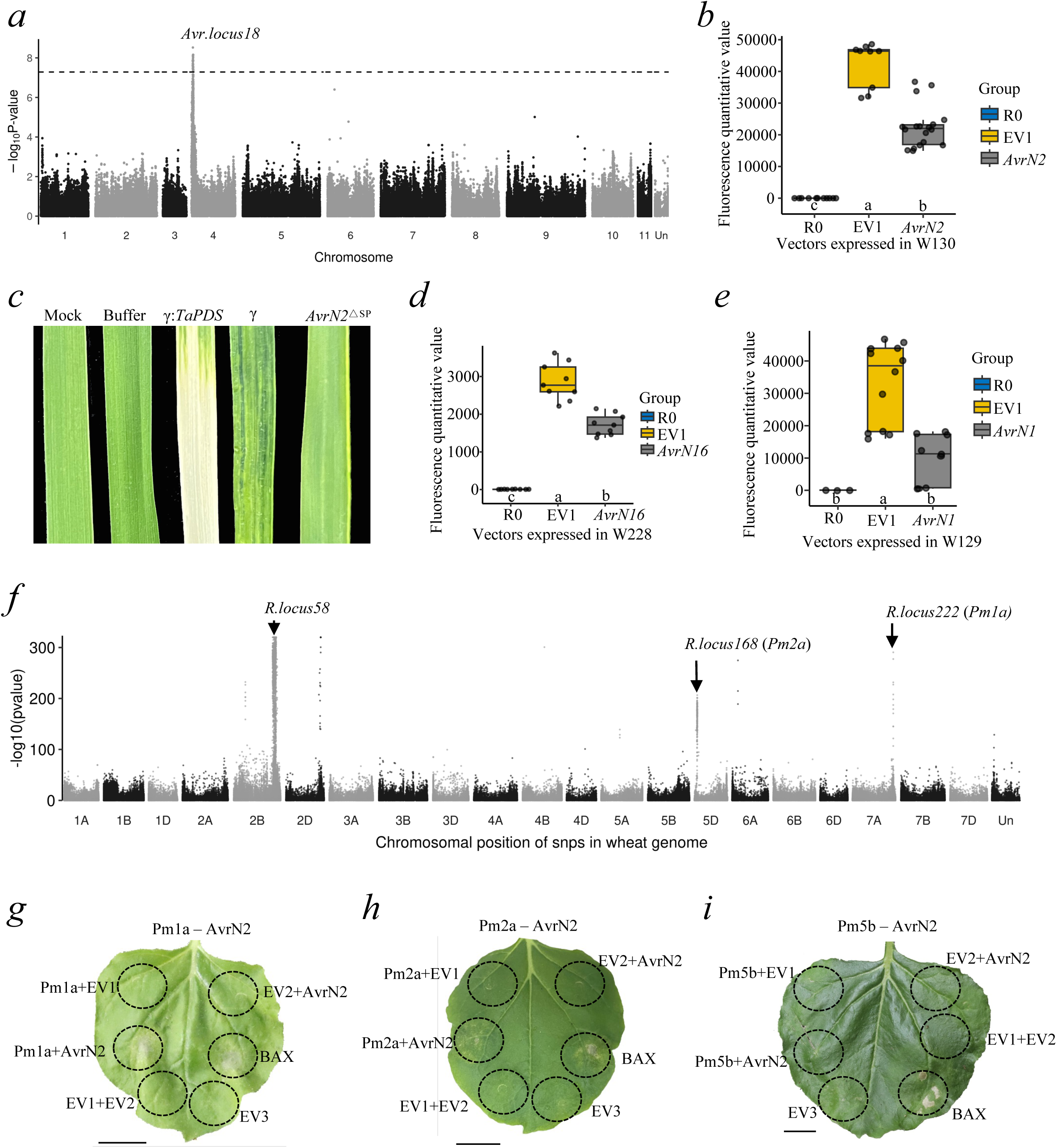
Molecular validation of candidate genes in *R*/*Avr* loci and *R-Avr* pairs. (a) Manhattan plot displaying GWAS results for the wheat line W130 across the *Bgt* panel containing 120 *Bgt* isolates. Association *P* values were obtained from a mixed linear model. The horizontal black dashed line represents the significant threshold determined by Bonferroni correction. *Avr.locus18* (*AvrN2*) is labelled as indicated. (b) Comparison of fluorescence levels between *AvrN2* constructs in protoplasts of wheat accession W130. ‘*AvrN2*’: over-expression vector of the *Avr* allele *AvrN2*; ‘EV1’: empty vector; ‘R0’ water control. Different letters above x-axis indicate statistically significant differences (*P* < 0.05) based on one-way ANOVA test. (c) BSMV-mediated expression of controls and the avirulence protein *AVRN2* in leaves of wheat line W130. Mock: wheat plants without any treatment. Buffer: wheat plants treated with the buffer solution used for application. γ-TaPDS: wheat plants overexpressing the etiolation gene TaPDS. γ: Wheat plants overexpressing the empty vector. *AvrN2*^Λ1sp^: Wheat plants overexpressing *AvrN2* with the signal peptide removed. Photos were taken at 10 dpi. (d) Comparison of fluorescence levels between *AvrN16* constructs in protoplasts of wheat accession W228. ‘*AvrN16*’: over-expression vector of the *Avr* allele *AvrN16*; ‘EV1’: empty vector; ‘R0’: water control. Different letters above x-axis indicate statistically significant differences (*P* < 0.05) based on one-way ANOVA test. (e) Comparison of fluorescence levels between *AvrN1* constructs in protoplasts of wheat accession W129. ‘*AvrN1*’: over-expression vector of the *Avr* allele *AvrN1*; ‘EV1’: empty vector; ‘R0’: water control. Different letters above x-axis indicate statistically significant differences (*P* < 0.05) based on one-way ANOVA test. (f) Association levels of *AvrN2* with all SNPs in the wheat genome based on a biEpi map. Association p-values were obtained using a general linear model. Each dot represents a single SNP. The *R* loci with known *Pm* genes are annotated using corresponding gene symbols. (g) Transient expression assay of the *Pm1a*-*AvrN2* pair in *N. benthamiana*. The *Avr* allele *AvrN2* was expressed alone or co-expressed with *Pm1a*. Empty vectors (EV1/2/3) and BAX were used as controls. OD_600_ = 0.5. Four-week-old *N. benthamiana* plants were used, and photos were taken at 4 dpi. Three biological replicates were performed with identical results. Scale bar: 1 cm. (h) Transient expression assay of the *Pm2a*-*AvrN2* pair in *N. benthamiana*. The *Avr* allele *AvrN2* was expressed alone or co-expressed with *Pm2a*. Empty vector (EV1/2/3) and BAX were used as controls. OD_600_ = 0.5. Four-week-old *N. benthamiana* plants were used, and photos were taken at 4 dpi. Three biological replicates were performed with the same results. Scale bar: 1 cm. (i) Transient expression assay of the *Pm5b*-*AvrN2* pair in *N. benthamiana*. The *Avr* allele *AvrN2* was expressed alone or co-expressed with *Pm5b*. Empty vector (EV1/2/3) and BAX were used as controls. OD_600_ = 0.5. Four-week-old *N. benthamiana* plants were used, and photos were taken at 4 dpi. Three biological replicates were performed with the same results. Scale bar: 1 cm.

The *Avr.locus56* locus was identified through GWAS within a ∼35 kb region on *Bgt* chromosome 10 (Supplementary Fig. 15a). Haplotype analysis and gene annotation revealed a single CESP gene *BgtE-20009,* designated as *AvrN16*, as the most likely candidate gene of *Avr.locus56* (Supplementary Fig. 15b). To validate its function, an avirulent haplotype of *AvrN16*, amplified from an avirulent isolate, was overexpressed in protoplasts of the cognate resistant wheat line W228 (Supplementary Table 12). This resulted in a significant reduction in fluorescence (Fig. 7d), indicating activation of cell death and validating that *AvrN16* (*BgtE-20009*) is a functional avirulence gene of *Avr.locus56*.

The *Avr.locus22* was mapped using GWAS into a ∼220 kb region on *Bgt* chromosome 4 (Supplementary Fig. 16a). Haplotype analysis and gene annotation identified *BgtE-5826*, a CESP gene designated as *AvrN1*, as the most likely candidate for this locus (Supplementary Fig. 16b). To functionally validate this gene, an avirulent haplotype of *AvrN1*, amplified from an avirulent isolate (Supplementary Table 12), was overexpressed in protoplasts of the cognate resistant wheat line W129. This resulted in a significant reduction in fluorescence (Fig. 7e), indicating activation of cell death and confirming that *AvrN1* (*BgtE-5826*) is a functional avirulence gene of *Avr.locus22*.

The *Avr.locus11* was mapped using GWAS into a ∼50 kb region on *Bgt* chromosome 2 (Supplementary Fig. 17a). Haplotype analysis and gene annotation within this region identified *BgtE-50180*, a CESP gene designated as *AvrN4*, as the most likely candidate for the locus (Supplementary Fig. 17b). To validate this gene’s function, an avirulent haplotypes of *AvrN4*, amplified from an avirulent isolate (Supplementary Table 12), was overexpressed in protoplasts of the cognate resistant wheat line W189. We did not, however, observe significant difference of the fluorescence signal compared to controls, indicating *AvrN4* (*BgtE-50180*) may not be a functional avirulence gene of *Avr.locus11* (Supplementary Fig. 17c). In summary, three of the four selected *Avr* genes were successfully cloned and their roles in triggering cell death in host cells in which the cognate *R* genes are expressed were verified.

In order to validate the mapped *R-Avr* interactions, two frequently identified *R-Avr* pairs R222-Avr35 and R168-Avr38 were selected, presumably *Pm1a-AvrPm1a* and *Pm2a*-*AvrPm2a*, for validation via transient co-expression in *Nicotiana benthamiana*. Our analysis predicted that *AvrPm1a* is recognized by multiple *R* loci, including its expected cognate *R* locus *R.locus222* with *Pm1a* (Supplementary Fig. 18a). The avirulent allele of *AvrPm1a* cloned from an avirulent isolate was expressed in *N. benthamiana* either alone or co-expressed with *Pm1a* cloned from a resistant wheat line. We observed that only the co-expression of *AvrPm1a* and *Pm1a* induced a hypersensitive response comparable to the positive control BAX, confirming that *AvrPm1a* is recognized by *Pm1a* and triggers cell death (Supplementary Fig. 18b). Similarly, we predicted that *AvrPm2a* is recognized by multiple *R* loci, including its expected *R* locus *R.locus168* with *Pm2a* (Supplementary Fig. 18c). The avirulent *AvrPm2a* allele was cloned and expressed in *N. benthamiana* either alone or co-expressed with *Pm2a* cloned from a resistant wheat line. Co-expression of *AvrPm2a* and *Pm2a* induced a hypersensitive response comparable to positive control BAX, demonstrating that *AvrPm2a* is specifically recognized by *Pm2a* and subsequently triggers cell death (Supplementary Fig. 18d). Therefore, we demonstrated our cross-species strategy successfully mapped previously cloned *R-Avr* interactions.

Our analysis predicted that *AvrN2* is likely recognized by multiple *R* loci, including *R.locus222* (*Pm1a*), *R.locus168* (*Pm2a*), and *R.locus58* (Fig. 7f; Supplementary Table 14). To validate this prediction, an avirulent haplotype of *AvrN2* was transiently expressed in *N. benthamiana* either alone or co-expressed with *Pm1a* or *Pm2a*. Co-expression of *AvrN2* with either *Pm1a* or *Pm2a* induced HR comparable to the positive control BAX, demonstrating that *AvrN2* is recognized by both *Pm1a* and *Pm2a*, thereby triggering cell death (Fig. 7g-h; Supplementary Table 13). Furthermore, as previously mentioned, *AvrN2* was also recognized in wheat line W130, which lacks both *Pm1a* and *Pm2a* (Supplementary Fig. 19). This finding suggests that *AvrN2* is recognized by at least one additional *R* gene distinct from *Pm1a* and *Pm2a*. In our study, this third *R* gene was mapped at *R.locus58*, which is overlapped with the *Pm6*/*Pm52* locus (Supplementary Table 7). A candidate gene from *R.locus58*, NLR gene *TraesCS2B02G496800* designated as *PmN2* (Supplementary Table 13), was inferred based on the Chinese Spring reference genome, but we failed to detect its interaction with *AvrN2* using tobacco system (Supplementary Fig. 20; Supplementary Table 14). Our analysis also predicted that *Avr.locus49* (*AvrPm1a.2*) is likely recognized by multiple *R* loci including *R.locus041* covering *Pm4b* (Supplementary Fig. 18e), but we failed to validate this recognition in *N. benthamiana* (Supplementary Fig. 18f). We also tested an unpredicted random *R-Avr* pair *Pm5b*-*AvrN2* as control and, as expected, in this instance found no HR in *N. benthamiana* (Fig. 7i).

In summary, our molecular validation provides evidence for three novel *Avr* genes and two new *R-Avr* interactions identified in our analysis. Combined with previous work, we showed that a novel *Avr* gene could be recognized by three *R* genes while a single *R* gene could recognize multiple *Avr* genes (Supplementary Fig. 18g). The collective data thus supports a predominantly multiple-to-multiple interaction model between *R* genes and *Avr* genes.

## Discussion

Our newly developed biGWAS method is capable of mapping *R*-*Avr* interactions. Among the 251 *R* loci and 65 *Avr* loci identified in our GWAS analysis, we successfully mapped four previously reported *R* genes (*Pm1a*, *Pm2a*, *Pm4b* and *Pm5b*) and two *Avr* genes (*AvrPm1a* and *AvrPm2a*). This demonstrates the validity of our experimental approach. Our cross-species GWAS identified 674 *R-Avr* interactions, while epistasis analysis between plant and pathogen SNPs revealed 1,409 *R-Avr* pairs. Notably, two previously reported interactions, *Pm1a-AvrPm1a* and *Pm2a*-*AvrPm2a*, were among the predicted pairs. We validated these interactions by cloning and co-expressing the corresponding *R* and *Avr* genes in *N. benthamiana* leaves. The successful identification of these known genes and interactions serves as a proof-of-concept for our methodology. Beyond recovering previously characterized genes and interactions, we cloned three novel *Avr* genes (*Bgt-50651*, *BgtE-5826* and *BgtE-20009*) and verified two novel interactions, demonstrating that both *Pm1a* and *Pm2a* recognize *Bgt-50651*. Our results highlight a complex network of *R-Avr* interactions, with some *R* and *Avr* loci acting as hubs. However, further investigation is needed to determine whether this multiple-to-multiple interaction model is exception or the norm in *R*-*Avr* interactions.

Our method identified candidate loci rather than candidate genes. Although we successfully verified three novel *Avr* genes and two new *R-Avr* interactions, we were unable to validate one candidate *Avr* locus (*Avr.locus11*) and two putative *R-Avr* interactions (*R.locus041*-*Avr.locus49* and *R.locus58*-*Avr.locus18*). As with any association mapping study, narrowing down from mapped interval to candidate genes is not always trivial. The *R* loci often encompass large genomic regions containing numerous candidate genes, yet only a single gene was selected for validation, and this may, or may not, be the correct one. Additionally, *R* genes may require ‘helper’ genes ^9^ or alternative splicing isoforms ^27^ for recognition of *Avr* genes, and the mapped *R* loci may not directly interact with the *Avr* genes but instead encode components of downstream immune signaling pathways. Given the limited scope of the molecular validation experiments performed in this study, we cannot exclude these possibilities. Moreover, some *R* genes may be absent from the reference genome due to introgressions from wild relatives or translocation events, such as those frequently reported in wheat ^28^. The integration of pan-genome resources with kmer-based approaches could be a promising avenue for overcoming these challenges and facilitating the cloning of *R* genes ^22^. Similar challenges exist in resolving *Avr* loci. Hafeez et al. ^25^ proposed a framework for leveraging an *R* gene atlas and pathogen diversity to guide breeding strategies. Our results contribute to this framework by providing a wealthy of functional *R* and *Avr* loci for future research and application in resistance breeding.

We identified a novel *Avr* gene which is recognized by three different *R* genes. In addition to *Pm1a* and *Pm2a*, the effector gene *Bgt-50651* is inferred to interact with a third *R* gene located near *Pm52* region. This inference is supported by multiple lines of evidence: (1) Our biGWAS analysis identified that *R.locus058* recognizes *Avr.locus18* (Supplementary Table 9), and this *R* locus overlapped with *Pm52*; (2) *Bgt-50651* from *Avr.locus18* triggered a cell death response in wheat line W130, which lacks neither *Pm1a* nor *Pm2a*; (3) *Avr.locus18* exhibited the strongest association signal in the *Avr* GWAS for wheat cultivar Liangxing 99 carrying the *Pm52* gene (Supplementary Fig. 21), suggesting that *Pm52* recognizes effectors located within this interval; and (4) Diagnostic PCR markers confirmed that W130 carries the *Pm52* gene. While recognition of a single *Avr* gene by multiple *R* genes has been reported in other plant-pathogen systems ^7^, this is the first such observation in wheat powdery mildew. Kloppe et al. ^10^ proposed a two-gene model to explain virulence toward *Pm1a* in *Bgt* following the identification of a second *Avr* gene for *Pm1a*. According to this model, virulence against *Pm1a* occurs only when both *AvrPm1a* and *AvrPm1a.2* evade recognition by *Pm1a*, a hypothesis that accounted for 95% of the observed *Pm1a* phenotypes in a panel of 216 isolates. Although both *AvrPm1a* and *AvrPm1a.2* were mapped via GWAS, these studies were conducted using different *Bgt* populations. We utilized a *Bgt* panel collected in China and identified *Bgt-50651* as a third *Avr* gene recognized by *Pm1a*. As shown in Supplementary Fig. 13c, the relationship between virulence level and the combinations of these three *Avr* genes is complex, with only ∼28% of virulence variation explained, likely due to epistatic interactions between different *Avr* genes. It can however be anticipated that as additional *R-Avr* interactions are cloned and characterized, a more comprehensive understanding of the molecular mechanisms underlying *R* gene recognition of *Avr* effectors will emerge.

Our study identified 251 *R* loci deployed in representative wheat germplasms based on a diverse panel of pathogen isolates. We could not detect rare, but large-effect *R* loci in this study. We provided the number of avirulence isolates for each wheat line for people who wants to clone *R* genes from a particular wheat line (Supplementary Table 1). Known *Pm* genes can be examined for those wheat lines to determine whether a new gene exists. Unlike *R* genes of wild relatives, these *R* genes have already been deployed in breeding. Additionally, 120 *Bgt* isolates were tested against the wheat panel, with 95% of the isolates associated with at least two *R* loci detected through GWAS. Compared to previous GWAS studies on plant-pathogen interactions ^29^, our study is conducted on a much larger scale, providing extensive insights for further research. Beyond the identified *Avr* loci corresponding to tested wheat lines, additional *Avr* loci may also be recognized, as the statistical power to detect such loci depends on allelic frequency. Among 92 *Avr* GWAS analyses yielding significant peaks, 83 were detected only at a single *Avr* locus, suggesting that resistance in most wheat lines is conferred by recognition of a single *Avr* gene. Given that each wheat line harbors approximately five *R* alleles, it is reasonable to hypothesize that multiple *R* genes may target a common *Avr* gene, or alternatively, that only one *R* gene is actively involved in resistance during infection. Our *R* GWAS results support the former hypothesis, as 95% of *R* GWAS analyses identified at least two peaks. The low frequency of *R* alleles in the population (averaging 2%) suggests that major *Pm* genes have not been widely deployed within this wheat panel. Accumulating *R* alleles appears to expand the range of avirulent isolates (Fig. 5g); for instance, a wheat line of our panel carries up to 35 *R* alleles and is resistance to roughly 70% of *Bgt* isolates. Whether hybrid breeding to accumulate additional *R* alleles can further enhance resistance remains to be tested. To facilitate breeding applications, we provide a presence-absence map of the 251 *R* alleles in our wheat germplasms (Supplementary Table 8). Since these germplasms are widely used by breeders, our dataset may aid in selecting hybrid parents carrying distinct sets of *R* alleles.

Mapping *Avr* genes by GWAS for *Bgt* typically involves handling a large number of isolates within a single laboratory. *Bgt* is airborne, and different isolates are indistinguishable to human eye. Moreover, long-term storage of this obligate biotrophic fungus is challenging ^30^. Fortunately, high density SNP data enabled us to perform quality control for cross-contamination events by examining the spectrum of heterozygotes. Although population-scale sequencing efforts have been reported for *Bgt* in several reports ^10, 31^, reusing those sequenced isolates for phenotyping in another laboratory is nearly impossible. Firstly, fungi samples may not withstand transportation as readily as plant seeds. Secondly, and arguably more importantly, live pathogen isolate samples must be carefully controlled during international exchange. Recent research indicates that the worldwide spread of powdery mildew is most likely due to human activity ^31^, further emphasizing the need for caution when transporting pathogen samples. Therefore, *Avr* GWAS typically requires sample collection from local environments and genome sequencing of the purified isolates. Kunz et al. ^6^ developed a ‘Avirulence depletion assay’ to combine *R* gene-mediated selection with bulk sequencing for rapid avirulence gene identification in wheat powdery mildew. They created a hybrid between two isolates that were virulent or avirulent to *Pm60*. Bulk sequencing for the progeny isolates grown on plants with or without *Pm60* lead to the cloning of *AvrPm60*. By contrast to that study, our biGWAS method identifies genetic variation present in both wheat germplasm and the recent pathogen population, rather than focusing on a single specific *R* gene. Interestingly, our study detected several *Avr* loci near the *AvrPm60* region and mapped the candidate *R* loci for them. Since both *R* and *Avr* genes tend to cluster on chromosomes ^6^, further work is required to resolve the candidate genes within these loci. In summary, we believe our work provide a unique resource for the study of wheat powdery mildew resistance.

The decreasing cost of sequencing have facilitated the application of genomics in plant-pathogen interaction studies. Lin et al. constructed a pan-NLRome of *Solanum americanum* immune receptor genes and screened 315 pathogen effectors across 52 *S. americanum* accessions, resulting in the identification and cloning of three *R*-*Avr* pairs ^32^. Compared to their work, our study assayed population variation in functional *R*/*Avr* alleles existed in the plant/pathogen population. As demonstrated by haplotype analysis of *R*/*Avr* candidates, resistance/avirulence may be caused by a different haplotype rather than that of the reference genome. Therefore, population-scale dissection of functional *R*/*Avr* alleles and their interactions is important for the understanding of plant-pathogen interactions. On the other hand, false-positive correlation between SNP and traits can arise due to population structure and other confounding factors ^33^. This issue is exacerbated when considering two co-evolving populations. Wang et al. developed a two-way mixed-effect model to detect cross-species genetic interactions, accounting for population structure including interactions between the genetic backgrounds of both the plant and the pathogen ^15^. Unfortunately, we were unable to implement this model in our study due to computational limitations.

We developed a cross-species joint GWAS approach, utilizing differentially enriched locus (DEL) analysis between phenotypically contrasting haplotype groups. This method was applied to dissect genetic interaction between wheat and one of its most damaging fungal pathogens. Our study design, encompassing 581 wheat lines and 245 pathogen isolates, surpasses the scale of previous studies ^15, 16, 33^. We identified 212 pairs of *R-Avr* interacting loci, including many previously known genes and interactions. Notably, our findings suggest a more complex *R*-*Avr* interaction mode than the traditional one-to-one model ^2^. Integrating our molecularly verified interactions with recent reports on non-canonical *R-Avr* interactions ^7–14^, we advocate for further research into the mechanisms underlying multiple-to-multiple interactions between plant hosts and pathogens. Unraveling this *R-Avr* landscape will undoubtedly be crucial for breeding crops with durable resistance.

## Methods

### Preparation of wheat germplasm resources and field trials

The wheat panel used in this study includes 262 lines from the Chinese-mini-core wheat germplasm ^17^ and 319 lines from the USDA wheat germplasm ^18^ (Supplementary Table 2). They were exome sequenced ^17, 18^, and their genotypes or genotyping data were downloaded from the those two previous studies. Additional wheat germplasm used includes cv. Zhongzuo 9504 for *Bgt* propagation and as a susceptible control, and cv. ND399 as a positive control of *Pm2a* testing. Wheat plants were grown in field for propagation. Seeds from 10 plants per accession were visually assessed for phenotypic uniformity every year, with plants exhibiting high deviations were discarded. Field trials were conducted from 2020 to 2023 at Changping Farm, IGDB/CAS, Beijing and at Zhao County Experimental Station, Shijiazhuang, Hebei Province. Two replicates per accession were planted in fields, with each replicate consisting of three rows. Seeds were sown in a row length of 1 m, with plant spacing of 10 cm and row spacing of 25 cm. Five uniform plants in the middle row were harvested. Standard agronomic practices were followed throughout the trials. Agronomic traits including heading date (HD), plant height (PH), spike length (SL) and number (SN), spikelet number (SPS) and kernel number (KPS) per spike were measured. The average values of all measurements represented the performances.

### Sampling, phenotyping, and DNA extraction of *Bgt* isolates

We collected *Bgt* field samples from 19 provinces covering the major wheat-growing regions of China from 2007 to 2022 (Supplementary Table 1). To purify isolates from each pathogenic specimen, a single tiny colony of spores were randomly picked and inoculated onto clean 9-day old leaves of the wheat cultivar Zhongzuo 9504, which were attached to petri dishes. This process was repeated six to ten times to obtain a purified isolate. In total, 245 purified isolates were obtained. These isolates were inoculated onto seedlings of the wheat population in a controlled greenhouse environment and phenotyped as described in Supplementary Note 1. The virulence level was indicated by linear infection types (IT) with IT values of 0-2 representing avirulence and 3-4 representing virulence. For DNA extraction, the *Bgt* spore powder was completely lysed before DNA extraction using NucleoBond^®^ HMW DNA kit (REF: 740160.20, MACHEREY-NAGEL GmbH & Co. KG, Dueren, Germany) following the standard protocol. DNA extraction was performed using the E.Z.N.A^®^ HP Fungal DNA Kit (REF:D3195-01, omega Bio-tek, Inc., Gerorgia, USA) following the protocol for samples with lower DNA contents.

### Whole-genome sequencing and variant discovery

All *Bgt* isolates used in this study were subjected to whole-genome sequencing (WGS) with an average depth of 74×, based on the reported reference genome size of 166 Mb ^20^. Library preparation and sequencing were performed using 150 bp paired-end model on the BGI DNBSEQ T7 platform at BGI (BGI-Shenzhen Co., Ltd., Shenzhen, China). The raw sequencing data, either generated in this study or downloaded from previous studies, were quality controlled and cleaned using software fastp v0.21.0 ^34^. Cleaned reads were aligned to the reference genome assembly v3.16 of *Bgt* isolate 96224 (GenBank accession number GCA_900519115.1) or the reference genome assembly RefSeq v1.0 of wheat variety Chinese Spring ^19^ using BWA MEM v0.7.17-r1188 ^35^. The resulting alignments files in BAM format of wheat were further processed for genomic germline variant calling using GATK v4.2.2.0 ^36^ following the best-practice guideline ^37^. Genomic germline variants of *Bgt* were called using DeepVariant v1.3.0 ^38^, while GLnexus v1.4.1 ^39^ was used to jointly genotype the GVCF files generated by DeepVariant. Genomic somatic variants between *Bgt* isolates were called using Strelka v2.9.10 ^40^. Multi-allelic or filter non-passed genomic variants were removed using VCFtools v0.1.16 ^41^. Only biallelic variants that passed quality control were retained for downstream analysis. All software tools used were run with their default parameter settings.

### GWAS and LD analysis

GWAS was conducted using the R package GAPIT v3.1.0 ^42^ with genomic variants filtered to include only those with a minor allele frequency (MAF) > 0.05 or 0.01 and a missing data rate < 10%. In GAPIT, four statistical models (MLM, MLMM, FarmCPU, and Blink) were applied to perform GWAS. Population structure was controlled using the first three principal components (PCs) as fixed effects and a kinship matrix as a random effect. To account for multiple testing, the Bonferroni correction was applied, setting the significance threshold at 0.05 divided by the number of variants tested. Variants with *P*-value below this Bonferroni-corrected threshold were considered as the significantly associated variants. Linkage disequilibrium was assessed using LDBlockShow v1.40 ^43^ with default settings.

### Identification and definition of *R* loci and *Avr* loci

To define an *Avr* locus, nearby GWAS peaks were merged based on the following criteria: if the most significant SNP of a peak (defined by Bonferroni cutoff: p-value ≤ 0.05/SNP-number) and the most significant SNP of another peak have a linkage disequilibrium (LD, R^2^) less than 0.8, these two peaks are merged into one locus, with the most significant SNP regarded as the representative SNP of the locus. This process is iterated multiple times to obtain the final set of loci. For defining an *R* locus, due to the lower SNP density, the method differs from that used for *Avr* loci. Both LD and physical distance are considered. The lead SNPs of two peaks are merged into one locus if they meet any of the following conditions: a. They are less than 0.25 Mb apart and have an LD of ≥ 0.2. b. Or they are less than 0.5 Mb apart and have an LD of ≥ 0.35. c. Or they are less than 1.0 Mb apart and have an LD of ≥ 0.5. d. Or they are less than 2.5 Mb apart and have an LD of ≥ 0.7 (Supplementary Note 3). For each *R* or *Avr* locus, the resistant or avirulent genotype of the representative SNP in GWAS results is used to represent the resistant or avirulent allele of the *R* or *Avr* locus, respectively.

### Annotation and enrichment analysis of GO terms and protein domains

Gene Ontology (GO) and protein domain annotations (InterProscan) for the wheat genes in the *Triticum aestivum* cv. Chinese Spring genome were obtained from database Ensembl Plants release 60 ^44^. Functional annotations of *Bgt* genes were transferred from *B. graminis* f. sp. *hordei* (*Bgh*) DH14 v4 based on sequence homology analysis. *Bgt* protein sequences were aligned to the *Bgh* protein database using BLASTp (Protein-Protein BLAST 2.16.0+), and the top match was used to transfer the corresponding GO terms and protein domains (InterProscan) from *Bgh* to *Bgt*. GO enrichment analysis was performed using the *enrichGO* function from the *R* package *clusterProfiler* ^45^ with default settings, considering a set number of nearby genes for each of the *R* or *Avr* loci in wheat or *Bgt*. Significance was determined using an adjusted *P*-value threshold of (pAdjustMethod=“BH”) < 0.05. Similarly, domain enrichment analysis (InterProscan) was conducted using the *enricher* function in *clusterProfiler* in the same way. Signal peptides in *Bgt* genes were annotated using SignalP v6.0 ^46^ with default settings. Effectors were predicted using EffectorP v3.0 ^47^ with default settings.

### Candidate gene identification and prioritization

To identify candidate genes within a given *R* or *Avr* locus, we selected the 60 genes closest to the representative SNP of the locus for consideration. Effector protein of *Bgt* genes were predicted using EffectorP v3.0 ^47^ with default settings. Classical NLR genes, defined as genes with NB-ARC and LRR domain predicted, or CESP genes, defined as genes with signal peptide and effector protein predicted, that exhibited a LD value greater than 0.8 with the representative SNP were deemed candidate genes. Only the closest candidate gene flanking or overlapping the representative SNP was selected for downstream molecular cloning and functional validation. For each *R* or *Avr* candidate gene, an avirulent *Bgt* isolate or a resistant wheat line was randomly chosen from the individuals exhibiting both the avirulent or resistant phenotype and genotype. The coding sequence (CDS) of the candidate gene was cloned from these individuals, as detailed in Supplementary Table 12 and S13. Eight individual colonies from the cDNA library of the chosen *Bgt* isolate or wheat line were screened by single-colony PCR. Three colonies showing positive amplification bands were randomly chosen for further validation through Sanger sequencing. If sequence inconsistencies were observed among these three monoclonal clones, three additional colonies will be amplified and sequenced for further confirmation. Finally, one of the most abundant sequences was selected randomly as the avirulent or resistant allele of the candidate gene for molecular validation.

### Genomic prediction

To evaluate the prediction accuracy of SNPs associated with wheat resistance to individual *Bgt* isolates, we implemented a 10-fold cross-validation framework using the GAPIT version *R* package v3.4.0. The resistance level of each wheat line was defined as the proportion of resistant responses (infection type < 2.3) across all tested *Bgt* isolates. Associated SNPs were analyzed in conjunction with the resistance levels. The first three principal components were included as fixed effects in the model to account for population structure. Genomic prediction was conducted using the genomic best linear unbiased prediction (gBLUP) model. To ensure robustness, the prediction process was repeated 100 times. For comparison, an equal number of randomly selected SNPs were subjected to the prediction procedure.

### Cross-species bi-directional GWAS (biGWAS)

To identify recognition pairs between *R* loci and *Avr* loci, bi-directional GWAS were conducted using forward and reverse differentially enriched locus (DEL) analysis. In the forward-DEL analysis, interactive *Avr* loci corresponding to a specific *R* locus (denoted as R_i_) were investigated. Haplotypes of R_i_ were determined from wheat GWAS results, distinguishing resistant haplotype (R_i_^hapR^) and susceptible haplotype (R_i_^hapS^). Wheat varieties were grouped based on whether they harbored R_i_^hapR^ or R_i_^hapS^. For wheat varieties in the R_i_^hapR^ group, the frequencies of *Avr* loci identified from *Bgt* GWAS results were calculated. The same analysis was performed for varieties in the R_i_^hapS^ group. For each *Avr* locus *j*, a Fisher exact test was used to assess significant enrichment of the *Avr* locus *j* in the R_i_^hapR^ group compared to the R_i_^hapS^ group (test conducted based on the following counts: n_1_=number of wheats having *Bgt* GWAS result including Avr_j_ in R_i_^hapR^ group, n_2_=number of wheats having *Bgt* GWAS result without Avr_j_ in R_i_^hapR^ group, n_3_=number of wheats having *Bgt* GWAS result including Avr_j_ in R_i_^hapS^ group, n_4_=number of wheats having *Bgt* GWAS result without Avr_j_ in R_i_^hapS^ group). A significance threshold of α=0.05 was applied to control the false positive rate. The same analysis was performed for reverse-DEL analysis to investigate the interactive *R* loci corresponding to a specific *Avr* locus. The weights of the identified pairs were calculated as n_1_/n_2_ – n_3_/n_4_. See Supplementary Note 3 for more details.

### Cross-species interaction analysis (biEpi)

The cross-species interaction at the genotypic level were evaluated by fitting a linear model: Y ∼ β_1_X_1_ + β_2_X_2_ + β_12_X_1_X_2_, where Y denotes the infection types (resistance/virulence level) between wheat and *Bgt*; X_1_ denotes the genotype of wheat (SNP_i_ or Gene_i_ in wheat); X_2_ demotes the genotype of *Bgt* (SNP_j_ or Gene_j_ in *Bgt*); X_1_X_2_ denotes the interaction between X_1_ and X_2_. A significant interaction term suggests dependency between the two genotypes. The genotype of wheat and *Bgt* were concatenated into a single matrix for the calculation. To evaluate the significance of the interaction effect between X_1_ and X_2_, the model was implemented in R language using the *lm* function. The model’s coefficients were extracted using the *summary* function, and the *P*-value corresponding to the interaction term was specifically retrieved to test its statistical significance. A threshold of α= 0.05/(number of test) was used to determine significance.

### Plasmid construction design and molecular cloning

The plasmids and oligonucleotide primers employed in the current study are catalogued in Supplementary Table 15. We utilized the previously reported plasmids ^48^ encoding the α, β, and γ chains of the ND18 strain of barley stripe mosaic virus (BSMV), designated as pBSMVα, pBSMVβ, and pBSMVγ. The CDSs of *TaPDS* and *AvrN2* were amplified and subcloned into the pBSMVγ vector. The CDSs of the wheat and *Bgt* genes with resistant or avirulent genotype were cloned within the binary vector of pCAMBIA2300 and unary vector of pJIT163-Ubi-EGFP using the pEASY^®-^Basic Seamless Cloning and Assembly Kit (TransGen, China). Sanger sequencing was performed on all vectors to confirm the accuracy of the insert sequences.

### Protoplast isolation from wheat

Approximately 50 seeds of wheat lines were sown in 13 cm pots filled with Martins Seed Raising and Cutting Mix, enriched with 3 g/L of Osmocote fertilizer. Seedlings were nurtured in a growth cabinet at 22 ±2 °C under a photoperiod of 14-h light (approximately 100 μmol m^-2^ s^-1^) and 10 hours darkness for a duration of 7 to 15 days. For protoplasts generation, the middle section of wheat leaves was used. Protoplast isolation and transformation followed previously published protocols ^26^. The liberated protoplasts were then resuspended to achieve final concentrations of either 2×10^5^ to 1×10^6^ cells/mL utilizing the MMG solution for individual transformation procedures. Transformation was conducted in the dark for 30 minutes, followed by incubation of the transformed protoplasts at 23°C for 48 h. After incubation, the protoplasts were collected by centrifugation at 100 ×*g* for 3 minutes. The precipitated products were lysed with 1× cell lysis buffer and briefly centrifuged at 13523 ×*g* for 15 seconds. For luciferase (LUC) activity assays, 20 ul of the supernatant was mixed with 100 ul of LUC substrate (luciferase) in a 96-well plate. LUC level was quantified using a Spectramax^®^ ID3 instrument (Molecular Devices, CA, USA). See Supplementary Note 4 for more details.

### BSMV-VOX

The BSMV-VOX experiment was conducted as previously described ^49^. BSMV, BSMV-PDS, and BSMV-AVRN2 were injected into tobacco leaves for transient expression in *Nicotiana benthamiana*. Wheat leaves at the jointing stage were inoculated with the transformational buffer, BSMV, BSMV-PDS, or BSMV-AVRN2. The procedure was repeated 3 times for each plant. In each experiment, three groups of controls, including FES buffer, wild-type BSMV, and BSMV-PDS. The virus infection symptoms were typically appeared 10 days post inoculation. See Supplementary Note 5 for more details.

### Transient expression in *Nicotiana benthamiana*

*Nicotiana benthamiana* plants were grown in a controlled chamber at a temperature of 23±2 °C with a 16 h photoperiod and were used for agroinfiltration at 3 to 4-week-old. *Agrobacterium tumefaciens* strains were cultured at 28°C with continuous shaking at 200 revolutions per minute (RPM) in Luria-Bertani (LB) broth, supplemented with the 50 mg/L Kanamycin. Bacterial cells were collected by centrifugation and subsequently resuspended in an infiltration solution containing 10 mM MES buffer at (pH 5.6), 10 mM MgCl_2_, and 150 μM acetosyringone. The cell suspension for each construct was adjusted to an optical density (OD_600_) of 0.5 (unless otherwise specified) and incubated at ambient temperature for 2 h prior to infiltration. Leaf samples were photographed 5 to 10 d post-infiltration to assess cell mortality. See Supplementary Note 6 for more details.

### PCR and qRT-PCR experiments

PCR amplifications were performed using the Tks Gflex™ DNA Polymerase Low DNA Kit (R091, TaKaRa, Beijing, China) according to the manufacturer’s recommendations. To study the expression of specific genes, qRT-PCR experiments were conducted using Taq Pro Universal SYBR qPCR Master Mix (Q712, VAZYME, Nanjing, China) according to the QuantStudio 5 guidelines. Each sample was assayed in triplicate to ensure reproducibility. Total RNA was extracted and purified using the FastPure Universal Plant Total RNA Isolation Kit (RC411, Vazyme, Beijing, China). cDNA synthesis was performed using the PrimeScript^TM^ RT Reagent Kit with gDNA Eraser (RR047, TAKARA, Beijing, China). The transcript levels of target genes were standardized using Triticum aestivum elongation factor 1 (*TraesCS4D01G197200*) as an internal reference. Gene expression variations were assessed by calculating the average 2^−ΔΔCt^ values, as described by Schmittgen and Livak ^50^. The thermal cycling protocol consisted of an initial denaturation at 95°C for 30 s, followed by 40 cycles of denaturation at 95°C for 10 s and annealing/extension at 60°C for 10 s. A melt curve analysis was performed to confirm the specificity of the amplification products. Primers for PCR and qRT-PCR targeting *Bgt* and wheat sequences are listed in Supplementary Table 15.

### Genetic relatedness analysis, correlation test, and statistical test

The genetic relationship matrix was calculated using GCTA v1.94.1 ^51^. Correlation coefficient and *P*-value were obtained in correlation test via Pearson (PCC) or Spearman (SCC) method of the function *cor.test* in the *R* package *stats* v4.1.2. To generate the identity-by-state (IBS) and identity-by-descent (IBD) matrices, genotype data were first pruned based on LD using PLINK v1.9 ^52^. The LD-pruned genotype data were then used to calculate IBS distance matrix and IBD matrix using PLINK. To assess statistical differences between groups, an analysis of variance (ANOVA) was performed, followed by a Tukey’s HSD (honestly significant difference) test. These analyses were conducted in R language using the *aov* and *TukeyHSD* functions of *R* package *stats* v4.1.2, respectively. Letters corresponding to significance groups have been assigned using *multcompLetters* function of the R package *multcompView* v0.1-10. All software or functions used were run at their default parameter settings.

## Data availability

All data generated in this study are included in this article and supplementary Information files as well as the public databases. The raw short-read whole-genome sequencing data reported in this paper have been deposited in the Genome Sequence Archive ^53^ of the National Genomics Data Center (https://ngdc.cncb.ac.cn/), Beijing Institute of Genomics (BIG), Chinese Academy of Sciences, under accession number CRA020312 for the 245 *Bgt* isolates. The genotype data in VCF format were deposited on Genomic Variation Map of National Genomics Data Center of BIG (https://ngdc.cncb.ac.cn/gvm/) under accession GVM000961 for the 245, GVM000962 for the 581 wheat lines and GVM000986 for the 10 RNAseq wheat lines.

## Acknowledgements

We are grateful to the people listed in Supplementary Table 1 for generously providing the initial *Bgt* specimens used in this study. We also extend our gratitude to Dr. Yiwen Li of IGDB, CAS, and Dr. Xueyong Zhang of ICS, CAAS, for their generous provision of the germplasm used in this study. This work was financially supported by the National Key Research and Development Program of China (2023YFF1000100), the Biological Breeding-National Science and Technology Major Project (2023ZD04073 and 2023ZD04076), the National Natural Science Foundations of China (32302369 and 32172001). We thank GF. Ren and YY. Wang for their assistance in maintaining the powdery mildew isolates, and M. Xu, Z. Liu, R. Mcintosh, B. Wulff and E. Akhunov for helpful discussions.

## Author contributions

J.X. lead the data analyses and manuscript writing. J.X. and Q.L. developed the *Bgt* in vitro culture and high-throughput phenotyping platform, performed the isolate purification, propagation and maintenance, and harvested sufficient conidiophore of *Bgt* isolates for sequencing. Q.L. prepared wheat seeds through field propagation and rogueing, lead the field trials and investigated the agronomic traits of wheat population with help from Z.Z., M.Y., C. Z., H.W., X.L., Y.H. and Z.C.. Q.L. lead the phenotyping with contribution from J.Z., C.Z., J.X., L.W., Y.H., Z.C., Y.W. and X.L.. L.W. lead the molecular experiments with contribution from F.Z., X.Z., Y.Y., H.S., M.L. and B.X.. D.Q., J.H., Y.L. and H.L. conducted isolates collection and maintenance before 2022. Y.W. built the web server for demonstration of biGWAS. J.W., Y.W., W.L., M.Z., Y.Z., and W.W. contributed to experimental design and interpretation of the results. A.R.F. contributed to manuscript writing. H.L. provided guidance for handling pathogen isolates and disease phenotyping, coordinated isolates collection and edited the manuscript. F.H. conceived the idea, supervised the project and edited the manuscript. All authors discussed the results and commented on the manuscript. All authors read and approved the final manuscript.

## Supplementary Figures

**Supplementary Figure 1. Genetic diversity and population structure of the *Bgt* panel.** (a) Schematic illustration of the *Bgt* isolation and purification process. The left panel represents the initial field-collected bulk population, consisting of a mixture of genetically diverse isolates. The right panel shows the purified *Bgt* isolate after 6-10 rounds of single-colony propagation on *ex vivo* wheat leaves. (b) Distribution of sequencing quality metrics across samples (all 245 *Bgt* isolates), including sequencing volume, properly mapped reads, and effective sequencing. (c) Distribution of the germline SNPs counts across all 245 *Bgt* isolates compared to reference genome. (d) The proportion of SNPs located in different genomic features, including intergenic regions, gene upstream and downstream regions, untranslated regions (UTRs), exons, introns, and splice sites. (e) PCA of all 245 *Bgt* isolates based on genome-wide SNP variation. The scatter plot shows the second (PC2) and third (PC3) principal components, with the percentage of explained variation indicated in parentheses. (f) Genome-wide distribution of nucleotide diversity (π) in the *Bgt* panel. π was calculated using a 10 kb non-overlapping window after filtering SNPs with a MAF 0.05. A vertical dotted line indicates the most common degree of π observed. (g) Linkage disequilibrium (LD) decay in the *Bgt* panel. The plot shows the decay of pairwise linkage disequilibrium (r²) as a function of physical distance (kb). LD was calculated using genome-wide SNPs, and the LOESS-smoothed curve represents the overall decay pattern. (h) Distribution of pairwise somatic SNPs among all *Bgt* isolates.

**Supplementary Figure 2. Diagram illustrating the integration of genotypes from Chinese-mini-core wheat lines and USDA wheat lines.** Each heatmap represents a genotype matrix, where homozygous reference alleles are shown as ‘2’ (red), homozygous alternative alleles are shown as ‘0’ (blue), heterozygous genotypes as ‘1’ (yellow), and missing data as black. Filtering criteria of each step are displayed along the arrows, with the number of retained variants indicated above the corresponding genotype heatmaps. GT: genotype.

**Supplementary Figure 3. Pairwise phenotypic correlation coefficients in wheat and *Bgt*.** (a) Density plot of phenotypic correlation coefficients for pairwise comparisons between *Bgt* isolates. (b) Density plot of phenotypic correlation coefficients for pairwise comparisons between wheat lines.

**Supplementary Figure 4. The genetic architecture of avirulence for the *Bgt* panel**. (a) Manhattan plots displaying GWAS results for wheat line W204 across the *Bgt* panel using four statistical models: MLM, MLMM, Blink, and FarmCPU. Statistical model name is labelled on the right. The horizontal dotted line represents the Bonferroni-corrected p-value threshold, while the vertical dotted lines highlight the common significant peak identified across all four models. (b) Frequency histogram of the number of *Avr* loci detected per trait (wheat line). (c) Enrichment analysis of putative effectors surrounding the 65 non-redundant *Avr* loci. The x-axis represents the number of surrounding genes considered per *Avr* locus. The y-axis shows the percentage of putative effectors among the selected genes. (d) The percentage of avirulent alleles for each identified non-redundant *Avr* loci in the *Bgt* panel. (e) PCA of genotypes of the 65 non-redundant *Avr* loci. The first two principal components (PC1 and PC2) are plotted. Gap statistics shows that one cluster is the optimal number of clusters. (f) Heatmap showing the virulent-avirulent allele distribution of nine *Avr* loci with cloned *AvrPm* genes in the *Bgt* panel. ‘AF’: allele frequency; ‘Frac.’: fraction.

**Supplementary Figure 5. The genetic architecture of resistance for the wheat panel**. (a) Manhattan plots displaying GWAS results for *Bgt* isolate Bgt0211 across the wheat panel using four statistical models: MLM, MLMM, Blink, and FarmCPU. Statistical model name is labelled on the right. The horizontal dotted line represents the Bonferroni-corrected p-value, while the vertical dotted line highlights the common significant peak identified across all four models. (b) Frequency histogram of the number of *R* loci detected per trait (*Bgt* isolate). (c) GO terms significantly enriched in the 40 nearest genes flanking the 251 *R* loci. (d) Protein domains significantly enriched in the 40 nearest genes flanking the 251 *R* loci. (e) Heatmap showing the resistant-susceptible allele distribution of eight *R* loci with cloned *Pm* genes in the wheat panel. ‘AF’: allele frequency; ‘Frac.’: fraction. (f) PCA of genotypes of all 251 *R* loci. The first two principal components (PC1 and PC2) are plotted. Gap statistics shows that one cluster is the optimal number of clusters.

**Supplementary Figure 6. Proportion comparisons and genetic effects of *R* alleles in the wheat panel.** (a) The percentage of *R* alleles for each identified non-redundant *R* loci in USDA wheat lines and Chinese-mini-core (CHN-mini-core) wheat lines. Each dot represents a wheat line. Different letters on top indicate statistically significant differences (*P* < 0.05) based on one-way ANOVA test. (b) The percentage of *R* alleles for each identified non-redundant *R* loci across different continents. Most wheat lines from Asia originate from China. Different letters on top indicate statistically significant differences (*P* < 0.05) based on one-way ANOVA test. (c) The percentage of *R* alleles for each identified non-redundant *R* loci in USDA germplasms across different continents. Most wheat lines from Asia are not from China. Different letters on top indicate statistically significant differences (*P* < 0.05) based on one-way ANOVA test. (d) The percentage of *R* alleles for each identified non-redundant *R* loci across different countries with more than seven wheat lines. Most wheat lines from Asia originate from China. Significant differences between means are indicated by asterisks. Different letters on top indicate statistically significant differences (*P* < 0.05) based on one-way ANOVA test. (e) Dot plot showing the resistance level and the accumulated genetic effects of *R* alleles for each wheat line. Regression curve with 95% confidence interval using general line model in shadow was draw as indicated. (f) Relationship between the resistance level and average genetic effect of *R* alleles per wheat line. A regression curve with a 95% confidence interval, fitted using a generalized linear model, is shown.

**Supplementary Figure 7. Diagram of biGWAS for mining *R*-*Avr* interacting pairs**. (a) Schematic representation of *R* locus identification (R1) in the wheat panel against *Bgt* isolate1 by detecting common significant GWAS in at least three statistical models. (b) Haplotype analysis of R1 classifies wheat lines into resistant (Hap-R) and susceptible (Hap-S) groups. (c) DEL analysis between peaks of Hap-R and Hap-S wheat lines identifies Avr1 as the candidate *Avr* gene recognized by R1.

**Supplementary Figure 8. Example of biGWAS: the *R.locus222-Avr.locus35* (*Pm1a-AvrPm1a*) interacting pair identified in forward-DEL analysis.** (a) Manhattan plots of 4 models in association mapping of *R.locus222*. Trait (Infection type) of *Bgt* isolate Bgt0194 was used as an example. The common peak at *R.locus222* was labelled with a black arrow. (b) The haplotype analysis of the representative SNP of *R.locus222*. The cumulative probability of phenotypes of the two haplotype groups was shown. HapR: resistant haplotype group, HapS: susceptible haplotype group. (c) Manhattan plots of HapR wheat lines and HapS wheat lines in *Bgt* genome. Results of representative HapR wheat lines and representative HapS wheat lines were shown. *Avr.locus35* was identified as a DEL between the two groups, as shown by the black arrow.

**Supplementary Figure 9. Example of biGWAS: the *Avr.locus35-R.locus222* (*AvrPm1a-Pm1a*) interacting pair identified in reverse-DEL analysis.** (a) Manhattan plots of 4 models in association mapping of *Avr.locus35*. Phenotypes of wheat line W104 was used as an example. The common peak at *Avr.locus35* was labelled with a black arrow. (b) The haplotype analysis of the representative SNP of *Avr.locus35*. The cumulative probability of phenotypes of the two haplotype groups was shown. HapA: avirulent haplotype group, HapV: virulent haplotype group. (c) Manhattan plots of HapA *Bgt* isolates and HapV *Bgt* isolate in wheat genome. Results of representative HapA *Bgt* isolates and representative HapV *Bgt* isolates were shown. *R.locus222* was identified as a DEL between the two groups, as shown by the black arrow.

**Supplementary Figure 10. Example of biGWAS: the *R.locus168-Avr.locus38* (*Pm2a-AvrPm2a*) interacting pair identified in forward-DEL analysis.** (a) Manhattan plots of 4 models in association mapping of *R.locus168*. Trait (Infection type) of *Bgt* isolate Bgt0393 was used as an example. The common peak at *R.locus168* was labelled with a black arrow. (b) The haplotype analysis of the representative SNP of *R.locus168*. The cumulative probability of phenotypes of the two haplotype groups was shown. HapR: resistant haplotype group, HapS: susceptible haplotype group. (c) Manhattan plots of HapR wheat lines and HapS wheat lines in *Bgt* genome. Results of representative HapR wheat lines and representative HapS wheat lines were shown. *Avr.locus38* was identified as a DEL between the two groups, as shown by the black arrow.

**Supplementary Figure 11. Example of biGWAS: the *Avr.locus38-R.locus168* (*AvrPm2a-Pm2a*) interacting pair identified in forward-DEL analysis.** (a) Manhattan plots of 4 models in association mapping of *Avr.locus38*. Phenotypes of wheat line W220 was used as an example. The common peak at *Avr.locus38* was labelled with a black arrow. (b) The haplotype analysis of the representative SNP of *Avr.locus38*. The cumulative probability of phenotypes of the two haplotype groups was shown. HapA: avirulent haplotype group, HapV: virulent haplotype group. (c) Manhattan plots of HapA *Bgt* isolates and HapV *Bgt* isolate in wheat genome. Results of representative HapA *Bgt* isolates and representative HapV *Bgt* isolates were shown. *R locus168* was identified as a DEL between the two groups, as shown by the black arrow.

**Supplementary Figure 12. Illustration of *R*-*Avr* interacting pairs identified in forward-DEL and reverse-DEL**. (a) Forward-DEL analysis reveals 47 *Avr* loci recognized by *251 R* loci, with each line linking an *R* locus to its corresponding *Avr* locus. Each grey box denotes a chromosome, with chromosome ID labelled in the middle. (b) Reverse-DEL analysis identifies 86 *R* loci recognizing 41 *Avr* loci, with each line connecting an *Avr* locus to its corresponding *R* locus. Each grey box denotes a chromosome, with chromosome ID labelled in the middle.

**Supplementary Figure 13. Overlaps and combinations of *R*/*Avr* loci in the biGWAS and biEpi maps.** (a) Venn diagram displaying the overlap of *R-Avr* interaction pairs identified through both biGWAS and biEpi analyses. (b) Box plot showing that the predicted *Pm1a-AvrPm1a* interacting pair results in significantly lower phenotyping scores (infection types) when *R* allele encounters the cognate *Avr* allele. ‘R’ represents the resistant allele of *Pm1a*; ‘r’ denotes the susceptible allele; ‘A’ denotes the avirulent allele of *AvrPm1a*, while ‘a’ represents the virulent allele. The red line indicates the median value. (c) Box plot depicting phenotypic differences among haplotype combinations of three *Avr* loci (*Avr.locus18*/*35*/*49*) recognized by *Pm1a*. Only wheat lines with *Pm1a* (*R* allele) were included in virulence calculation. ‘0’ represents the avirulent allele, and ‘4’ represents the virulent allele. The edges of the box represent the first and third quartiles, and whiskers extend 1.5 times the interquartile range. Different letters indicate statistically significant differences (*P* < 0.05, one-way ANOVA). (d) Box plot showing phenotypic differences among haplotype combinations of three *Avr* loci (*Avr.locus23*/*30*/*33*) recognized by *Pm3*. ‘0’ represents the avirulent allele, and ‘4’ represents the virulent allele. The edges of the box represent the first and third quartiles, with whiskers spanning 1.5 times the interquartile range. Different letters indicate statistically significant differences (*P* < 0.05, one-way ANOVA). (e) The accumulation of *Avr* loci recognized along with the number of *R* loci included. The dotted line refers to the minimal number of *R* loci needed to reach the maximal number of *Avr* loci recognized.

**Supplementary Figure 14. Association mapping and molecular validation of the *Avr.locus18***. (a) Manhattan plot of GWAS for mapping *Avr.locus18*, showing the 250 kb genomic region surrounding the associated peak on chromosome 4. SNPs are color-coded based on their linkage disequilibrium (LD) strength with the top SNP (measured as the squared correlation coefficient, R^2^). The accompanying LD heatmap illustrates pairwise LD (R²) among SNPs in the 250 kb interval, with stronger LD represented by higher R² values (red) and weaker LD by lower R² values (white). (b) The annotation information of the genes in the mapping interval of *Avr.locus18*. ‘1’: yes; ‘0’: no. (c) Transient expression of *AvrN2* in W130 protoplasts was observed under fluorescence microscope after 48 h with W5 medium inoculation. Scale bar, 200 μm. (d) The relative expression level of *AvrN2* in BSMV-vox plants obtained via qRT-PCR. *AvrN2*-specific primers were used. C: control group, ‘#’: plant ID. Three technical replicates were done for each group. The edges of the box represent the first and third quartiles, with whiskers spanning 1.5 times the interquartile range. Different letters on top of boxes indicate statistically significant differences (*P* < 0.05, one-way ANOVA). (e) BSMV-mediated expression of controls and the avirulence protein AVRN2 in leaves of susceptible wheat cv. Zhongzuo 9504. Mock: wheat plants without any treatment. Buffer: wheat plants treated with the buffer solution used for application. γ-TaPDS: wheat plants overexpressing the etiolation gene TaPDS. γ: Wheat plants overexpressing the empty vector. AvrN2^Λ1sp^: Wheat plants overexpressing *AvrN2* with the signal peptide removed. Photos were taken at 10 dpi.

**Supplementary Figure 15. Association mapping and molecular validation of the *Avr.locus56***. (a) Manhattan plot of GWAS for mapping *Avr.locus56*, showing the 35 kb genomic region surrounding the associated peak on chromosome 10. SNPs are color-coded by their LD strength with the top SNP (R^2^). The corresponding LD heatmap illustrates pairwise LD (R²) among SNPs in the 250 kb genomic interval with stronger LD represented by higher R² values (red) and weaker LD by lower R² values (white). (b) The annotation information of the genes in the mapping interval of *Avr.locus56*. ‘1’: yes; ‘0’: no.

**Supplementary Figure 16. Association mapping and molecular validation of the *Avr.locus22***. (a) Manhattan plot of GWAS for mapping *Avr.locus22*, showing the ∼220 kb genomic region surrounding the associated peak on chromosome 4. SNPs are color-coded by their LD strength with the top SNP (R^2^). The corresponding LD heatmap illustrates pairwise LD (R²) among SNPs in the 220 kb genomic interval with stronger LD represented by higher R² values (red) and weaker LD by lower R² values (white). (b) The annotation information of the genes in the mapping interval of *Avr.locus22*. ‘1’: yes; ‘0’: no.

**Supplementary Figure 17. Association mapping and molecular validation of the *Avr.locus11***. (a) Manhattan plot of GWAS for *Avr.locus11* mapping, showing the ∼50 kb genomic region surrounding the associated peak on chromosome 2. SNPs are color-coded by their LD strength with the top SNP (R^2^). The corresponding LD heatmap illustrates pairwise LD (R²) among SNPs in the 50 kb genomic interval with stronger LD represented by higher R² values (red) and weaker LD by lower R² values (white). (b) The annotation information of the genes in the mapping interval of *Avr.locus11*. ‘1’: yes; ‘0’: no. (c) Comparison of fluorescence levels between *AvrN4* constructs in protoplasts of wheat accession W189. ‘*AvrN4*’: over-expression vector of the *Avr* allele *AvrN4*; ‘EV1’: empty vector; ‘R0’ water control. Different letters above x-axis indicate statistically significant differences (*P* < 0.05) based on one-way ANOVA test.

**Supplementary Figure 18. Molecular validation of the predicted *R-Avr* interacting pairs**. (a) Association levels of *AvrPm1a* with all SNPs in the wheat genome based on the epistasis map. Association p values were obtained using a general linear model. Each dot represents a single SNP. The know *Pm* genes are annotated with arrows and corresponding gene symbols. (b) Transient expression assay of the *Pm1a-AvrPm1a* interacting pair in *N. benthamiana*. The *Avr* allele *AvrPm1a* was expressed alone or co-expressed with *Pm1a* in *N. benthamiana leaves*. Empty vector (EV1/2/3) and BAX were used as controls. OD_600_ = 0.5. Four-week-old *N. benthamiana* plants were used, and photos were taken at 4 dpi. Three biological replicates were performed and showed consistent results. Scale bar: 1 cm. (c) Association levels of *AvrPm2a* with all SNPs in the wheat genome based on the epistasis map. Association p values were obtained using a general linear model. Each dot represents a single SNP. The know *Pm* genes are annotated with arrows and corresponding gene symbols. (d) Transient expression assay of the *Pm2a-AvrPm2a* interacting pair in *N. benthamiana*. The *Avr* allele *AvrPm2a* was expressed alone or co-expressed with *Pm2a* in *N. benthamiana* leaves. Empty vector (EV1/2/3) and BAX were used as controls. OD_600_ = 0.5. Four-week-old *N. benthamiana* plants were used, and photos were taken at 4 dpi. Three biological replicates were performed and showed consistent results. Scale bar: 1 cm. (e) Association levels of *AvrPm1a.2* with all SNPs in the wheat genome based on the epistasis map. Association p-values were obtained using a general linear model. Each dot represents a single SNP. The know *Pm* genes are annotated with arrows and corresponding gene symbols. (f) Transient expression assay of the *Pm4b-AvrPm1a.2* interacting pair in *N. benthamiana*. The *Avr* allele *AvrPm1a.2* was expressed alone or co-expressed with *Pm4b* in *N. benthamiana* leaves. Empty vector (EV1/2) and BAX were used as controls. OD_600_ = 0.5. Four-week-old *N. benthamiana* plants were used, and photos were taken at 4 dpi. Three biological replicates were performed and showed consistent results. Scale bar: 1 cm. (g) Experimentally verified *R-Avr* interactions displayed as a network graph.

**Supplementary Figure 19. Gel electrophoresis results for PCR products of *Pm1a* and *Pm2a*.** (a) Agarose gel electrophoresis results for PCR products of *Pm1a* diagnostic marker *Pm1aSTS1*. Wheat line W140 was used as a positive control, in which the full-length cDNA sequence of *Pm1a* was amplified and confirmed by Sanger sequencing. Wheat line Chinese Spring known for its absence of *Pm1a* was used as a negative control. The target amplicon was labelled with a red arrow. (b) PAGE gel electrophoresis results for PCR products of *Pm2a* diagnostic marker *Pm2a-map-3*. Wheat line ND399 known for its presence of *Pm2a* was used as a positive control. Wheat line Chinese Spring known for its absence of *Pm2a* was used as a negative control. The target amplicon was labelled with a red arrow.

**Supplementary Figure 20. Association mapping and molecular validation of the *R.locus058***. (a) Manhattan plot of GWAS for mapping *PmN2* in trait Bgt0199, showing the ∼1.83 Mb genomic region surrounding the associated peak on chromosome 2B. SNPs are color-coded by their LD strength with the top SNP (R^2^). The corresponding LD heatmap illustrates pairwise LD (R²) among SNPs in the 1.83 kb genomic interval with stronger LD represented by higher R² values (red) and weaker LD by lower R² values (white). (b) The annotation information of the 11 candidate genes in the mapping interval of *R.locus058*. One candidate, *TraesCS2B02G496800* designated as *PmN2*, were randomly chosen for downstream functional validation. (c) Transient expression assay of the *PmN2-AvrN2* interacting pair in *N. benthamiana*. The *Avr* allele of *AvrN2* was expressed alone or co-expressed with *PmN2* in *N. benthamiana* leaves. Empty vector (EV1/2/3) and BAX were used as controls. OD 600 = 0.5. Four-week-old *N. benthamiana* plants were used, and photos were taken at 4 dpi. Three biological replicates were performed and showed consistent results. Scale bar: 1 cm.

**Supplementary Figure 21. *Avr* GWAS results of the wheat cultivar Liangxing 99 carrying *Pm52*.** (a) Manhattan plot of GWAS for mapping *Avr* loci with Liangxing 99 in the *Bgt* panel. *Avr.locus18* on chromosome 4 was identified as the most significant peak.

## Supplementary Tables

**Supplementary Table 1. The information of the tested *Bgt* isolates used in current study.**

**Supplementary Table 2. The information of the tested wheat lines used in current study.**

**Supplementary Table 3. The information of the identified 65 *Avr* loci.**

**Supplementary Table 4. The significantly enriched GO terms for genes flanking the 65 *Avr* loci.** For each *Avr* loci, the flanking 40 genes were included in GO enrichment analysis.

**Supplementary Table 5. The significantly enriched protein domains for genes flanking the 65 *Avr* loci.** For each *Avr* loci, the flanking 40 genes were included in protein domains enrichment analysis.

**Supplementary Table 6. Virulent-avirulent allelic map of the 65 *Avr* loci in 120 *Bgt* isolates.**

**Supplementary Table 7. The information of the identified 251 *R* loci.**

**Supplementary Table 8. Resistant-susceptible allelic map of the 251 *R* loci in 581 wheat lines.**

**Supplementary Table 9. *R-Avr* interacting pairs predicted in biGWAS analysis.**

**Supplementary Table 10. *R-Avr* interacting pairs predicted in biEpi analysis.**

**Supplementary Table 11. *R-Avr* interacting pairs predicted in both biGWAS and biEpi analyses.**

**Supplementary Table 12. The information of *Avr* alleles used in validation experiments.**

**Supplementary Table 13. The information of *R* alleles used in validation experiments.**

**Supplementary Table 14. The summary of molecular validation experiments.**

**Supplementary Table 15. Primers and vectors used in this study.**

