## Supplemental Figures for "A biGWAS strategy reveals the genetic architecture of the interaction between wheat and *Blumeria graminis* f. sp. *tritici*"

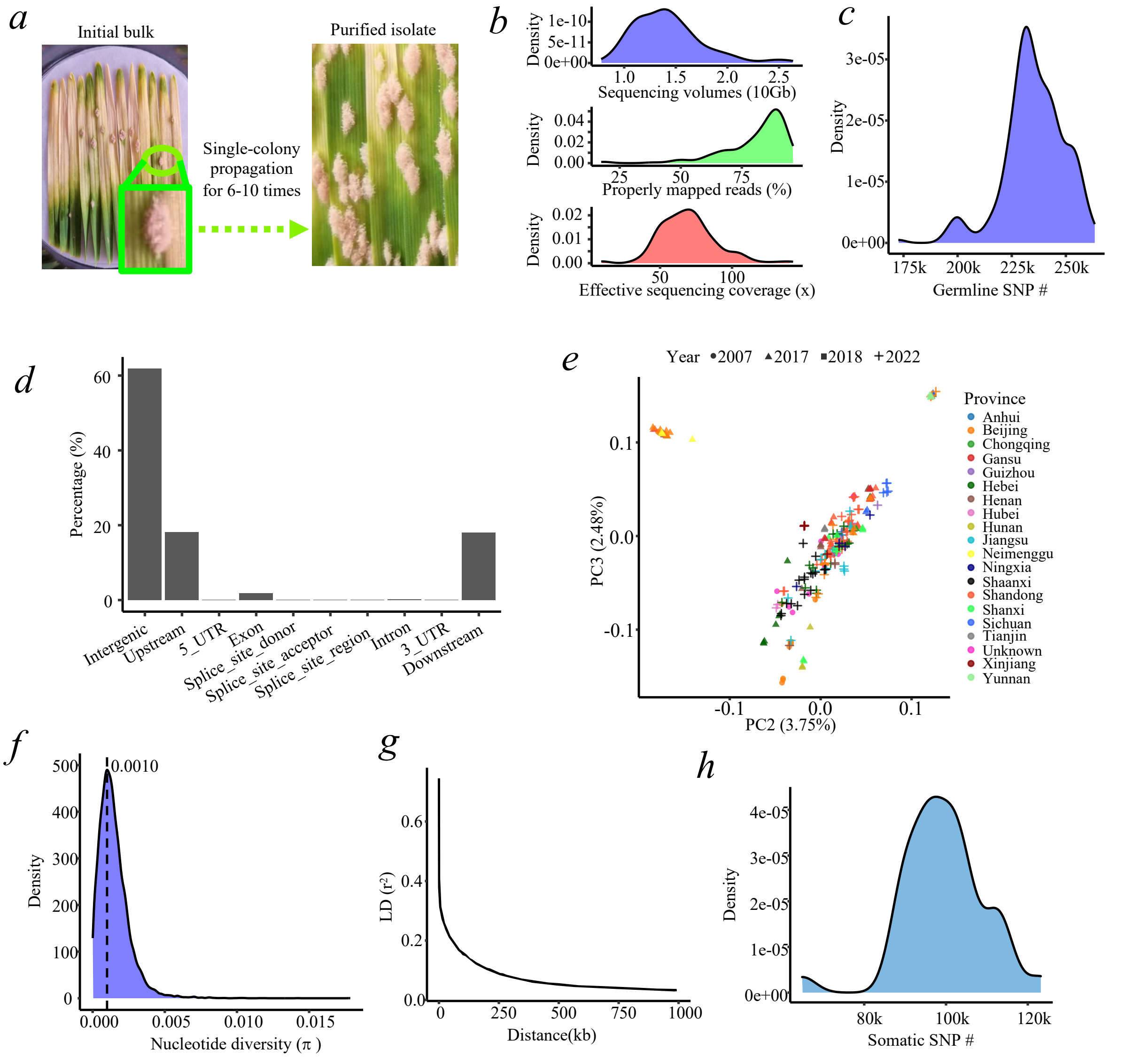


**Supplementary Figure 1. Genetic diversity and population structure of the *Bgt* panel.** (a) Schematic illustration of the *Bgt* isolation and purification process. The left panel represents the initial field-collected bulk population, consisting of a mixture of genetically diverse isolates. The right panel shows the purified *Bgt* isolate after 6-10 rounds of single-colony propagation on *ex vivo* wheat leaves. (b) Distribution of sequencing quality metrics across samples (all 245 *Bgt* isolates), including sequencing volume, properly mapped reads, and effective sequencing. (c) Distribution of the germline SNPs counts across all 245 *Bgt* isolates compared to reference genome. (d) The proportion of SNPs located in different genomic features, including intergenic regions, gene upstream and downstream regions, untranslated regions (UTRs), exons, introns, and splice sites. (e) PCA of all 245 *Bgt* isolates based on genome-wide SNP variation. The scatter plot shows the second (PC2) and third (PC3) principal components, with the percentage of explained variation indicated in parentheses. (f) Genome-wide distribution of nucleotide diversity ($\pi$) in the *Bgt* panel. $\pi$ was calculated using a 10 kb non-overlapping window after filtering SNPs with a MAF 0.05. A vertical dotted line indicates the most common degree of $\pi$ observed. (g) Linkage disequilibrium (LD) decay in the *Bgt* panel. The plot shows the decay of pairwise linkage disequilibrium ($r^2$) as a function of physical distance (kb). LD was calculated using genome-wide SNPs, and the LOESS-smoothed curve represents the overall decay pattern. (h) Distribution of pairwise somatic SNPs among all *Bgt* isolates.

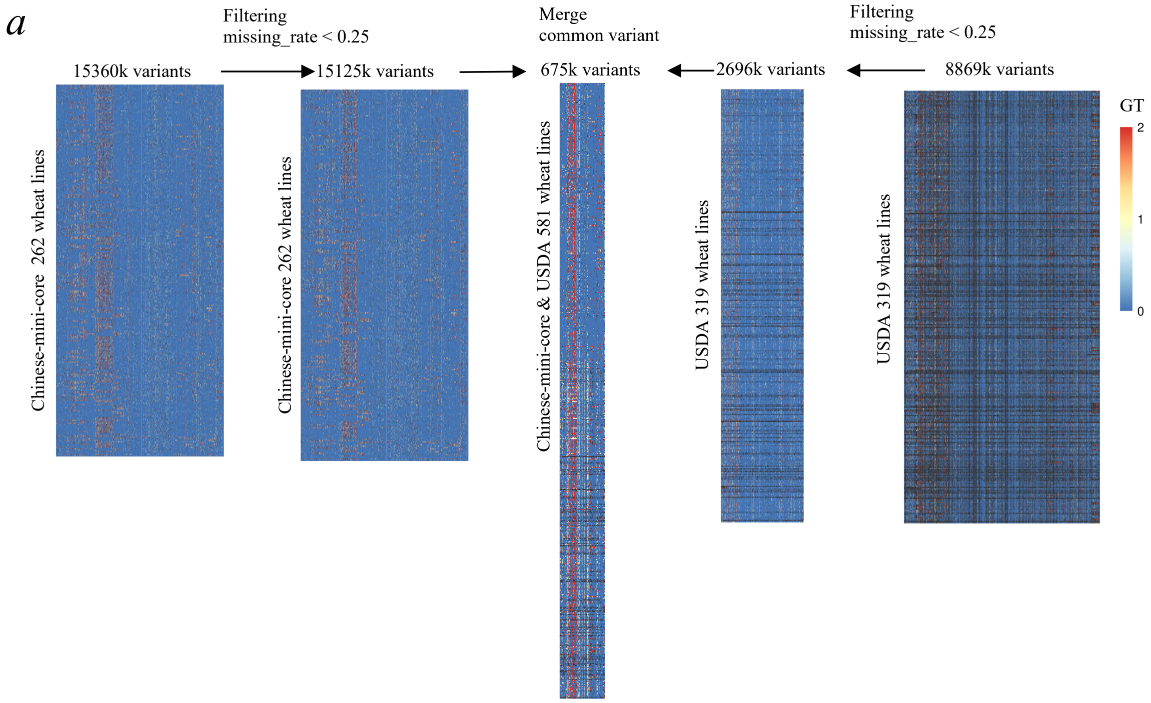


**Supplementary Figure 2. Diagram illustrating the integration of genotypes from Chinese-mini-core wheat lines and USDA wheat lines.** Each heatmap represents a genotype matrix, where homozygous reference alleles are shown as '2' (red), homozygous alternative alleles are shown as '0' (blue), heterozygous genotypes as '1' (yellow), and missing data as black. Filtering criteria of each step are displayed along the arrows, with the number of retained variants indicated above the corresponding genotype heatmaps. GT: genotype.

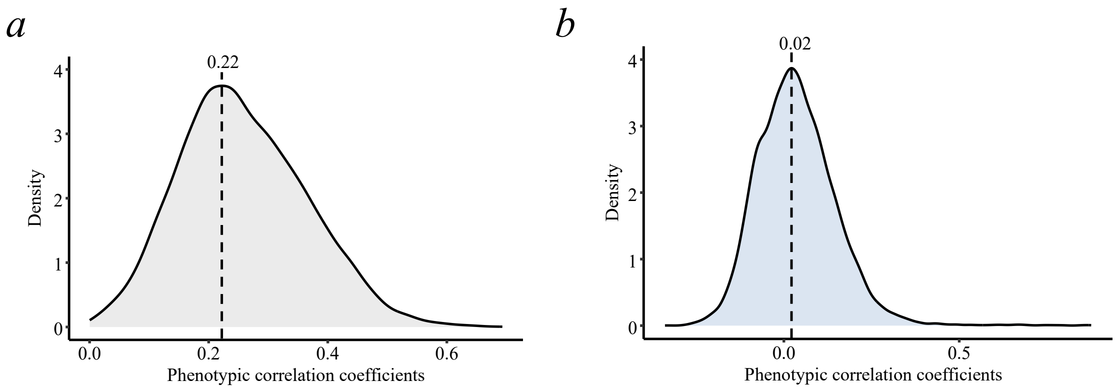


**Supplementary Figure 3. Pairwise phenotypic correlation coefficients in wheat and *Bgt*.** (a) Density plot of phenotypic correlation coefficients for pairwise comparisons between *Bgt* isolates. (b) Density plot of phenotypic correlation coefficients for pairwise comparisons between wheat lines.

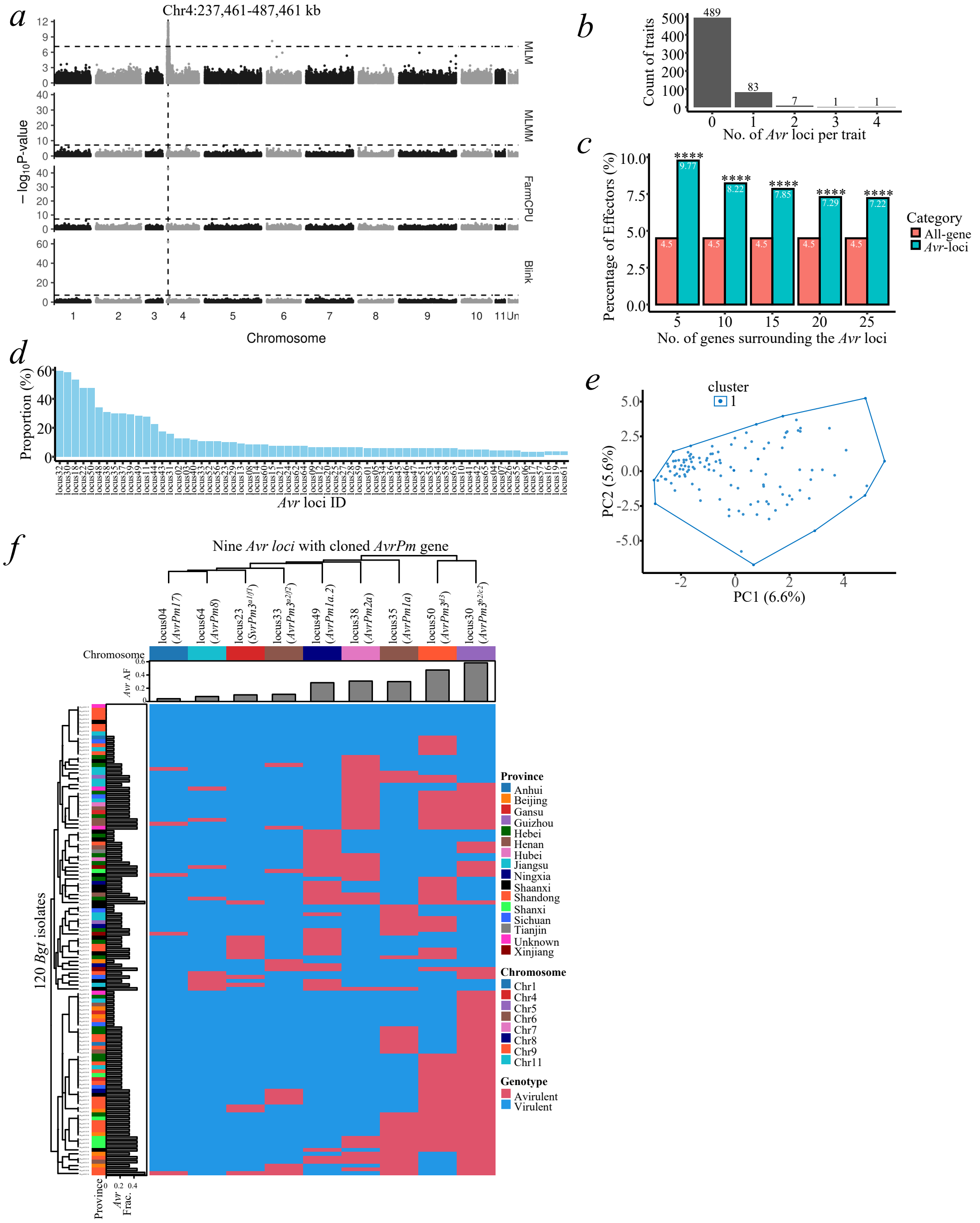


**Supplementary Figure 4. The genetic architecture of avirulence for the *Bgt* panel**. (a) Manhattan plots displaying GWAS results for wheat line W204 across the *Bgt* panel using four statistical models: MLM, MLMM, Blink, and FarmCPU. Statistical model name is labelled on the right. The horizontal dotted line represents the Bonferroni-corrected p-value threshold, while the vertical dotted lines highlight the common significant peak identified across all four models. (b) Frequency histogram of the number of *Avr* loci detected per trait (wheat line). (c) Enrichment analysis of putative effectors surrounding the 65 non-redundant *Avr* loci. The x-axis represents the number of surrounding genes considered per *Avr* locus. The y-axis shows the percentage of putative effectors among the selected genes. (d) The percentage of avirulent alleles for each identified non-redundant *Avr* loci in the *Bgt* panel. (e) PCA of genotypes of the 65 non-redundant *Avr* loci. The first two principal components (PC1 and PC2) are plotted. Gap statistics shows that one cluster is the optimal number of clusters. (f) Heatmap showing the virulent-avirulent allele distribution of nine *Avr* loci with cloned *AvrPm* genes in the *Bgt* panel. 'AF': allele frequency; 'Frac.': fraction.

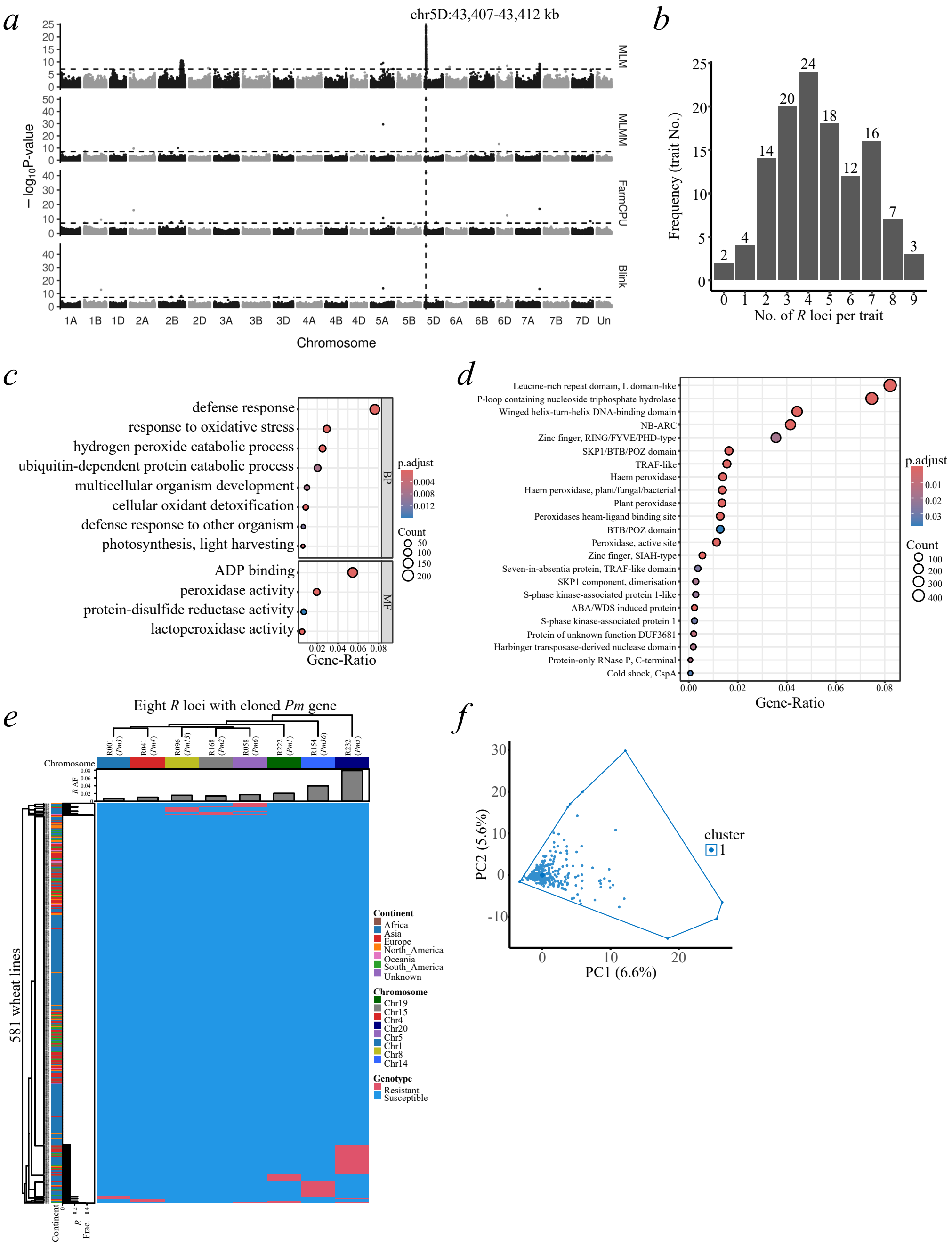


**Supplementary Figure 5. The genetic architecture of resistance for the wheat panel**. (a) Manhattan plots displaying GWAS results for *Bgt* isolate Bgt0211 across the wheat panel using four statistical models: MLM, MLMM, Blink, and FarmCPU. Statistical model name is labelled on the right. The horizontal dotted line represents the Bonferroni-corrected p-value, while the vertical dotted line highlights the common significant peak identified across all four models. (b) Frequency histogram of the number of *R* loci detected per trait (*Bgt* isolate). (c) GO terms significantly enriched in the 40 nearest genes flanking the 251 *R* loci. (d) Protein domains significantly enriched in the 40 nearest genes flanking the 251 *R* loci. (e) Heatmap showing the resistant-susceptible allele distribution of eight *R* loci with cloned *Pm* genes in the wheat panel. ‘AF’: allele frequency; ‘Frac.’: fraction. (f) PCA of genotypes of all 251 *R* loci. The first two principal components (PC1 and PC2) are plotted. Gap statistics shows that one cluster is the optimal number of clusters.

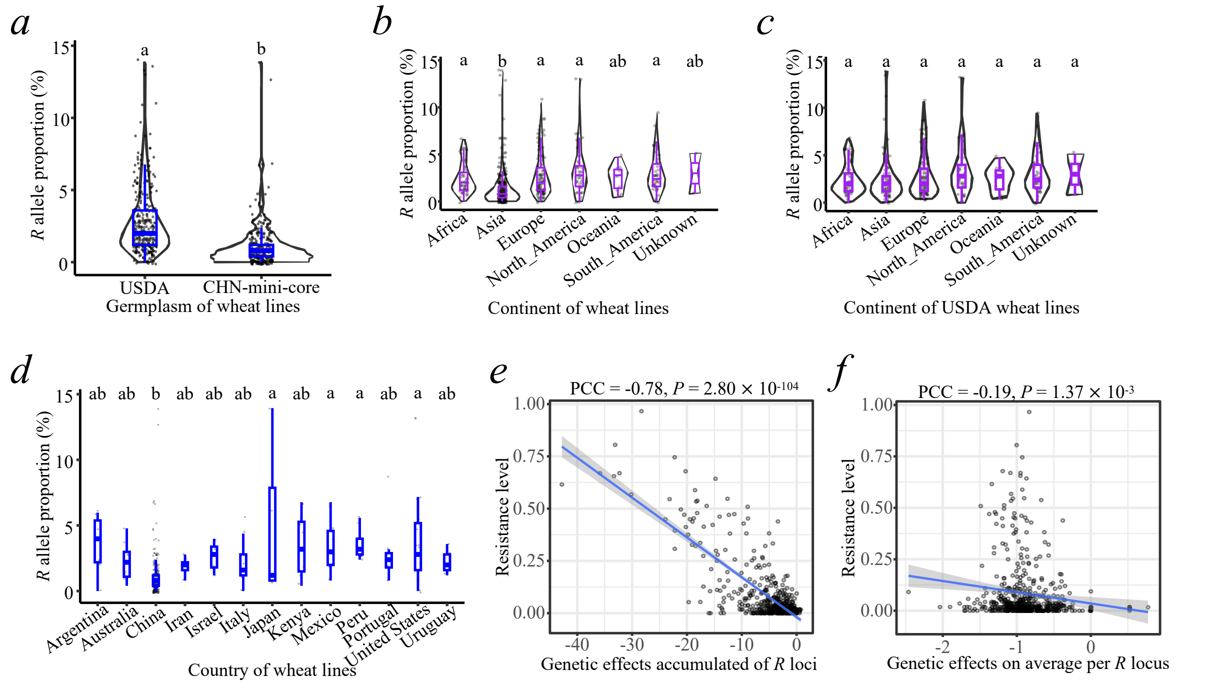


**Supplementary Figure 6. Proportion comparisons and genetic effects of *R* alleles in the wheat panel**. (a) The percentage of *R* alleles for each identified non-redundant *R* loci in USDA wheat lines and Chinese-mini-core (CHN-mini-core) wheat lines. Each dot represents a wheat line. Different letters on top indicate statistically significant differences ($P < 0.05$) based on one-way ANOVA test. (b) The percentage of *R* alleles for each identified non-redundant *R* loci across different continents. Most wheat lines from Asia originate from China. Different letters on top indicate statistically significant differences ($P < 0.05$) based on one-way ANOVA test. (c) The percentage of *R* alleles for each identified non-redundant *R* loci in USDA germplasms across different continents. Most wheat lines from Asia are not from China. Different letters on top indicate statistically significant differences ($P < 0.05$) based on one-way ANOVA test. (d) The percentage of *R* alleles for each identified non-redundant *R* loci across different countries with more than seven wheat lines. Most wheat lines from Asia originate from China. Significant differences between means are indicated by asterisks. Different letters on top indicate statistically significant differences ($P < 0.05$) based on one-way ANOVA test. (e) Dot plot showing the resistance level and the accumulated genetic effects of *R* alleles for each wheat line. Regression curve with 95% confidence interval using general line model in shadow was draw as indicated. (f) Relationship between the resistance level and average genetic effect of *R* alleles per wheat line. A regression curve with a 95% confidence interval, fitted using a generalized linear model, is shown.

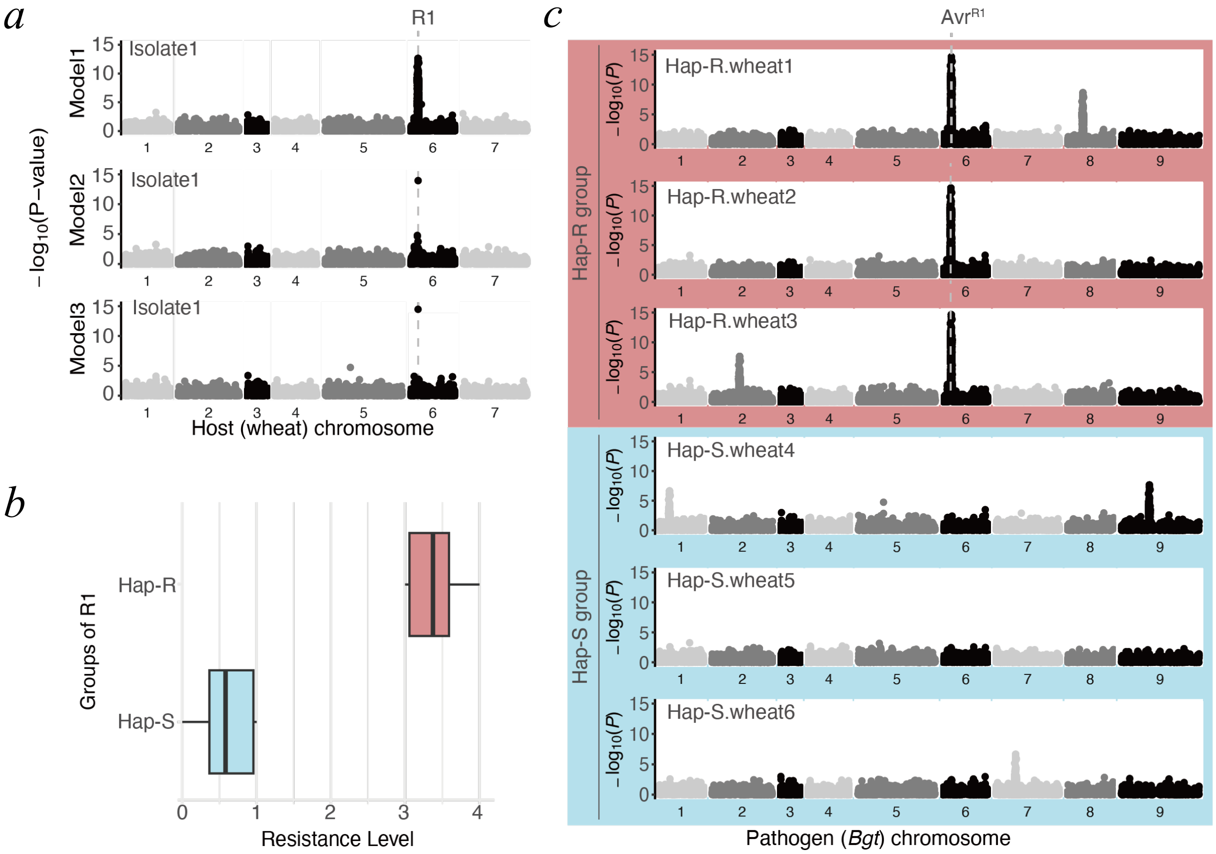


**Supplementary Figure 7. Diagram of biGWAS for mining *R-Avr* interacting pairs**. (a) Schematic representation of *R* locus identification (R1) in the wheat panel against *Bgt* isolate1 by detecting common significant GWAS in at least three statistical models. (b) Haplotype analysis of R1 classifies wheat lines into resistant (Hap-R) and susceptible (Hap-S) groups. (c) DEL analysis between peaks of Hap-R and Hap-S wheat lines identifies Avr1 as the candidate *Avr* gene recognized by R1.

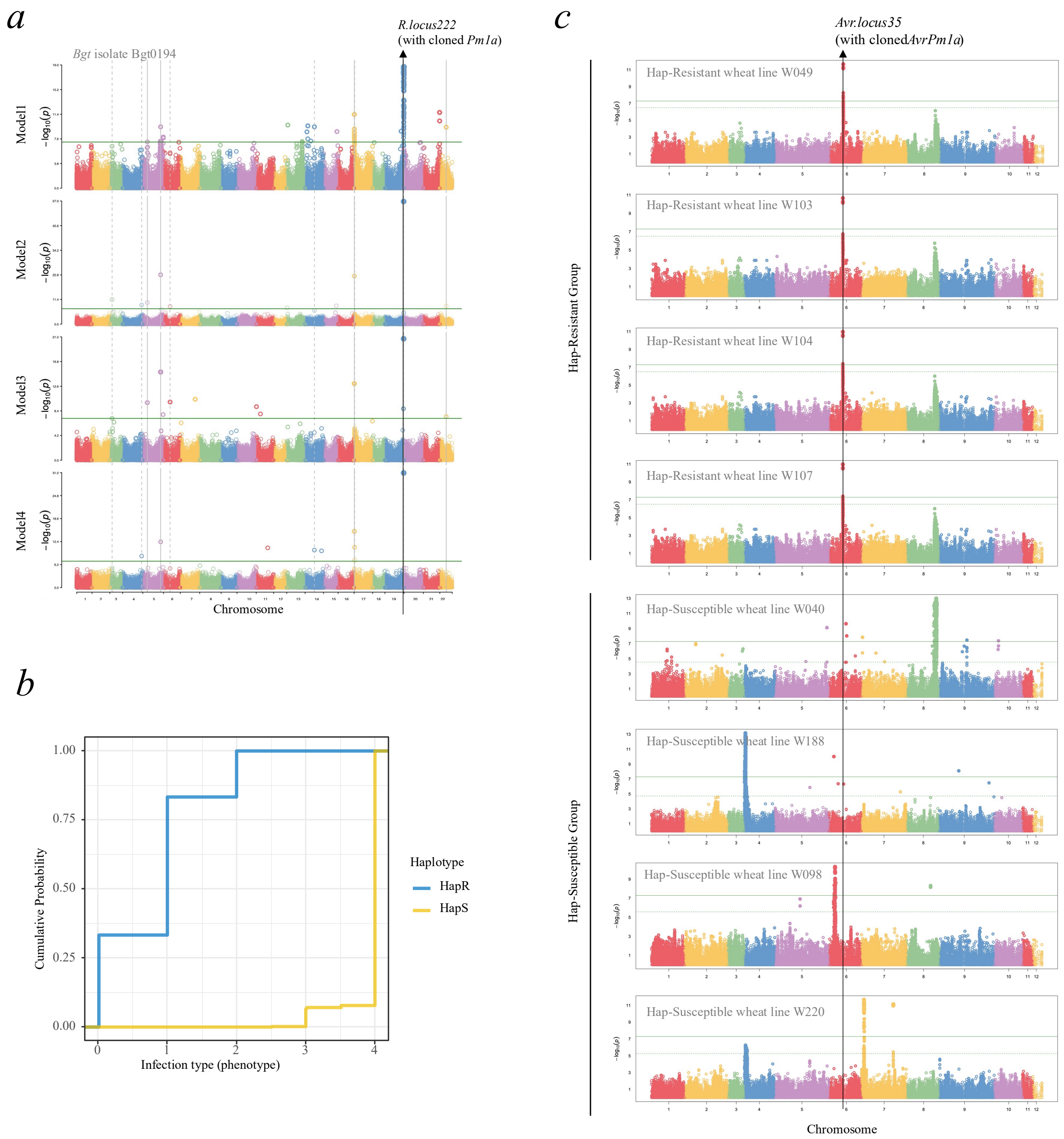


**Supplementary Figure 8. Example of biGWAS: the *R.locus222-Avr.locus35* (*Pm1a-AvrPm1a*) interacting pair identified in forward-DEL analysis.** (a) Manhattan plots of 4 models in association mapping of *R.locus222*. Trait (Infection type) of *Bgt* isolate Bgt0194 was used as an example. The common peak at *R.locus222* was labelled with a black arrow. (b) The haplotype analysis of the representative SNP of *R.locus222*. The cumulative probability of phenotypes of the two haplotype groups was shown. HapR: resistant haplotype group, HapS: susceptible haplotype group. (c) Manhattan plots of HapR wheat lines and HapS wheat lines in *Bgt* genome. Results of representative HapR wheat lines and representative HapS wheat lines were shown. *Avr.locus35* was identified as a DEL between the two groups, as shown by the black arrow.

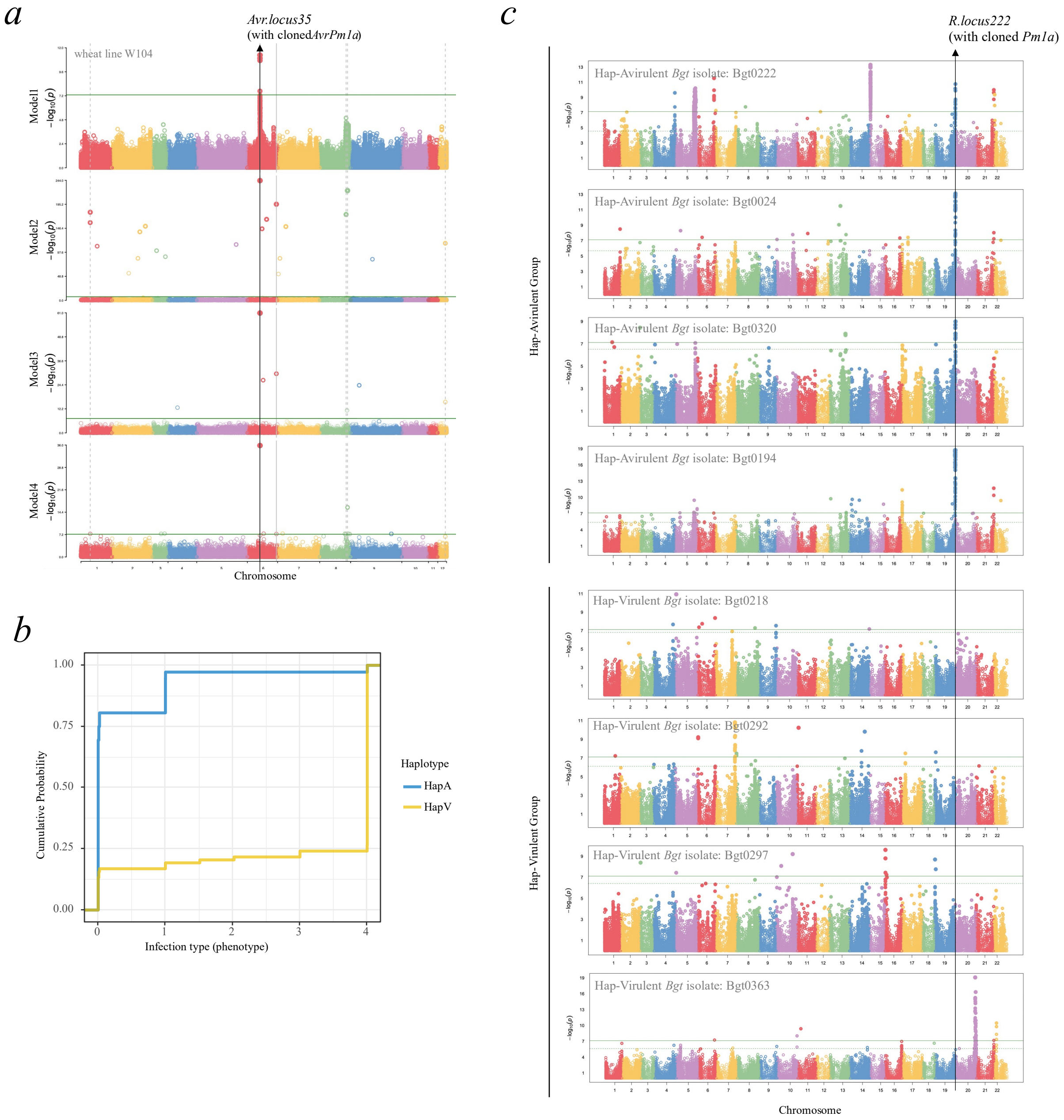


**Supplementary Figure 9. Example of biGWAS: the *Avr.locus35-R.locus222* (*AvrPm1a-Pm1a*) interacting pair identified in reverse-DEL analysis.** (a) Manhattan plots of 4 models in association mapping of *Avr.locus35*. Phenotypes of wheat line W104 was used as an example. The common peak at *Avr.locus35* was labelled with a black arrow. (b) The haplotype analysis of the representative SNP of *Avr.locus35*. The cumulative probability of phenotypes of the two haplotype groups was shown. HapA: avirulent haplotype group, HapV: virulent haplotype group. (c) Manhattan plots of HapA *Bgt* isolates and HapV *Bgt* isolate in wheat genome. Results of representative HapA *Bgt* isolates and representative HapV *Bgt* isolates were shown. *R.locus222* was identified as a DEL between the two groups, as shown by the black arrow.

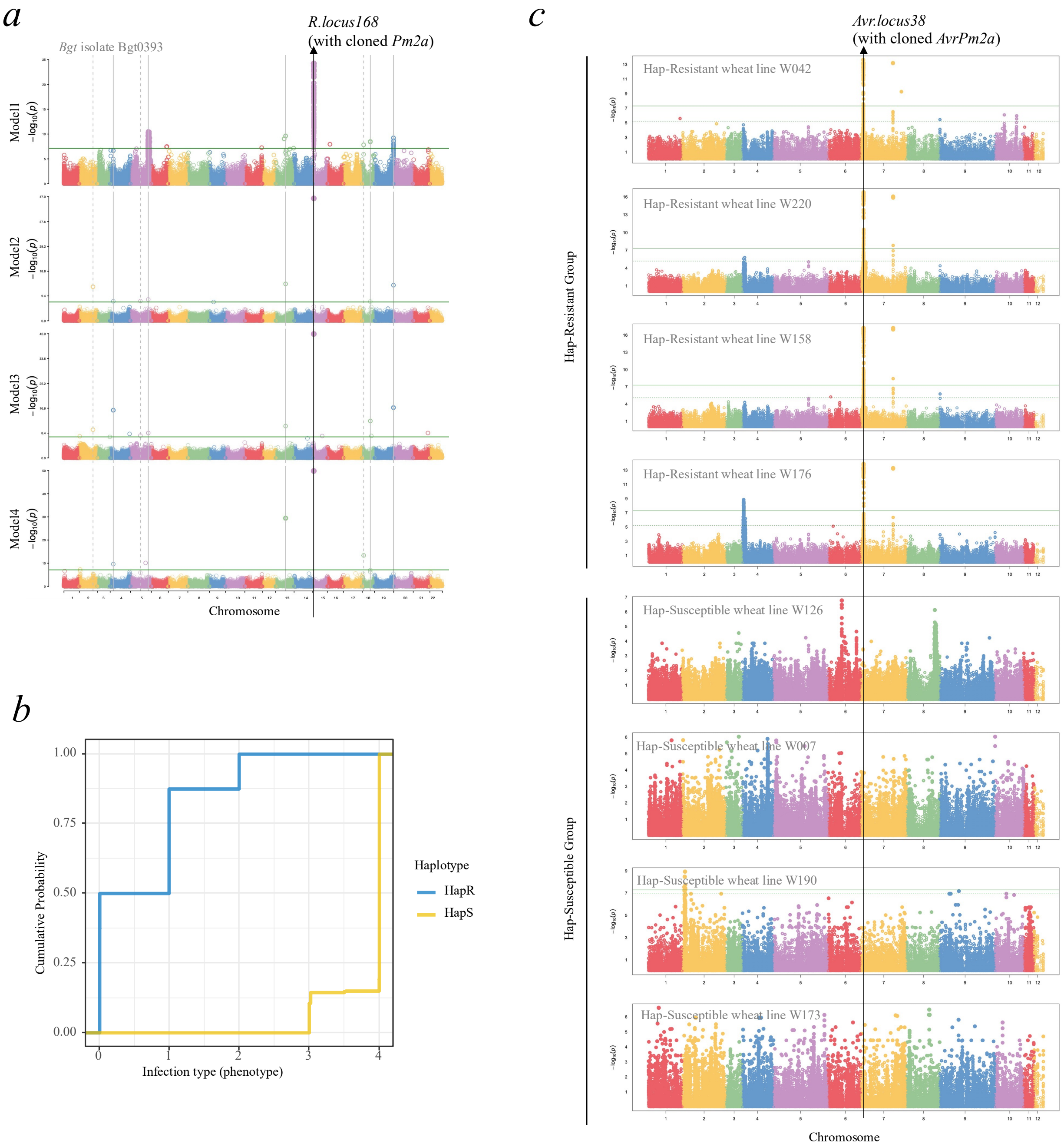


**Supplementary Figure 10. Example of biGWAS: the *R.locus168-Avr.locus38* (*Pm2a-AvrPm2a*) interacting pair identified in forward-DEL analysis.** (a) Manhattan plots of 4 models in association mapping of *R.locus168*. Trait (Infection type) of *Bgt* isolate Bgt0393 was used as an example. The common peak at *R.locus168* was labelled with a black arrow. (b) The haplotype analysis of the representative SNP of *R.locus168*. The cumulative probability of phenotypes of the two haplotype groups was shown. HapR: resistant haplotype group, HapS: susceptible haplotype group. (c) Manhattan plots of HapR wheat lines and HapS wheat lines in *Bgt* genome. Results of representative HapR wheat lines and representative HapS wheat lines were shown. *Avr.locus38* was identified as a DEL between the two groups, as shown by the black arrow.

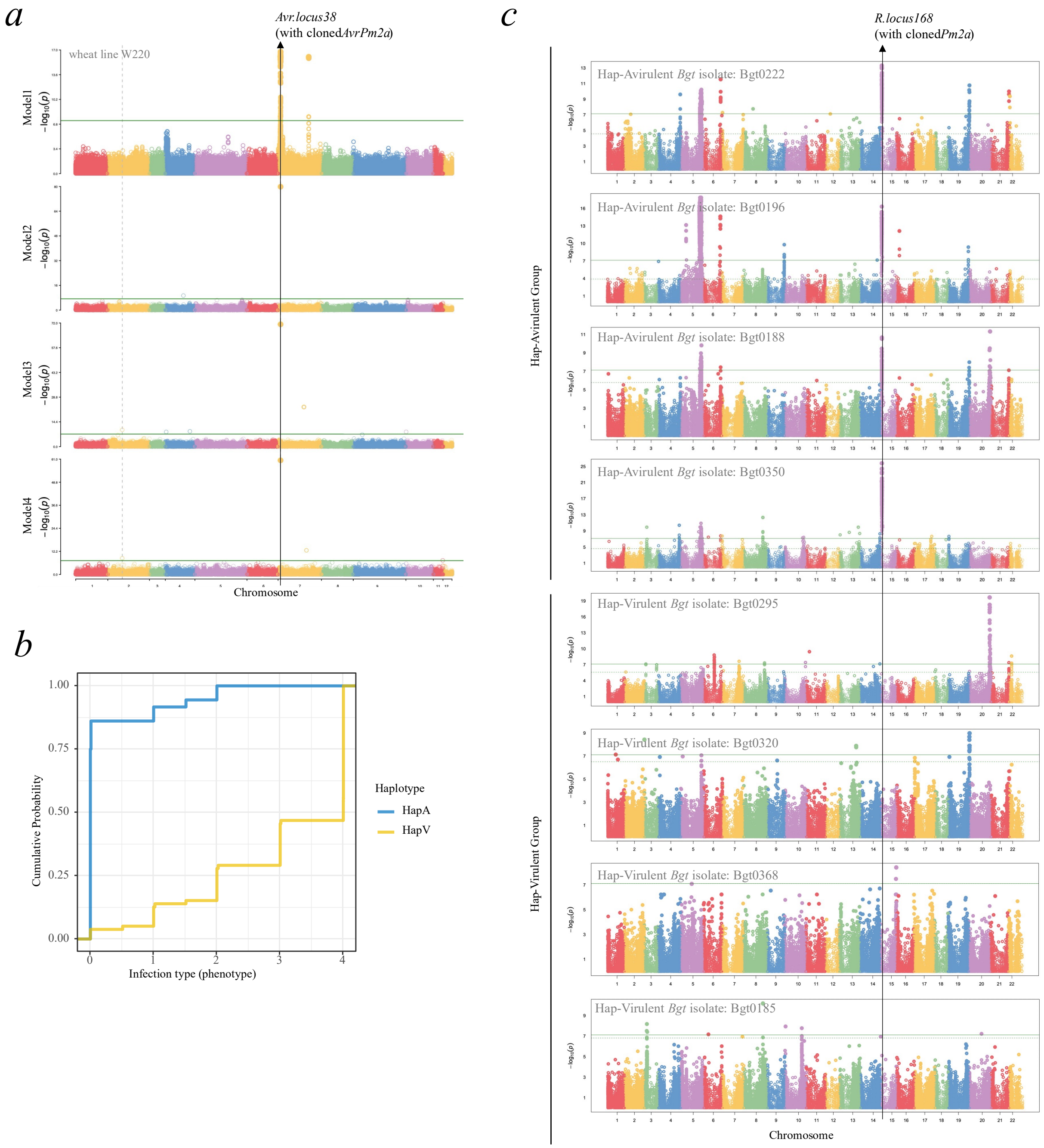


**Supplementary Figure 11. Example of biGWAS: the *Avr.locus38-R.locus168* (*AvrPm2a-Pm2a*) interacting pair identified in forward-DEL analysis.** (a) Manhattan plots of 4 models in association mapping of *Avr.locus38*. Phenotypes of wheat line W220 was used as an example. The common peak at *Avr.locus38* was labelled with a black arrow. (b) The haplotype analysis of the representative SNP of *Avr.locus38*. The cumulative probability of phenotypes of the two haplotype groups was shown. HapA: avirulent haplotype group, HapV: virulent haplotype group. (c) Manhattan plots of HapA *Bgt* isolates and HapV *Bgt* isolate in wheat genome. Results of representative HapA *Bgt* isolates and representative HapV *Bgt* isolates were shown. *R locus168* was identified as a DEL between the two groups, as shown by the black arrow.

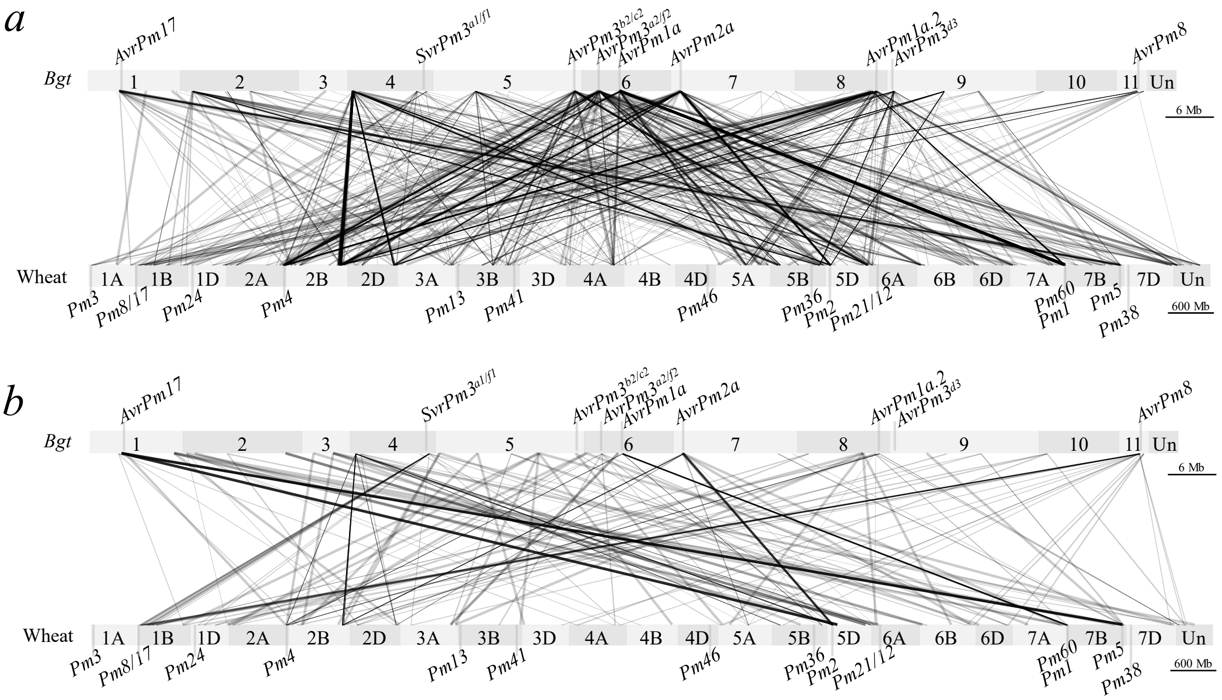


**Supplementary Figure 12. Illustration of *R-Avr* interacting pairs identified in forward-DEL and reverse-DEL**. (a) Forward-DEL analysis reveals 47 *Avr* loci recognized by *251 R* loci, with each line linking an *R* locus to its corresponding *Avr* locus. Each grey box denotes a chromosome, with chromosome ID labelled in the middle. (b) Reverse-DEL analysis identifies 86 *R* loci recognizing 41 *Avr* loci, with each line connecting an *Avr* locus to its corresponding *R* locus. Each grey box denotes a chromosome, with chromosome ID labelled in the middle.

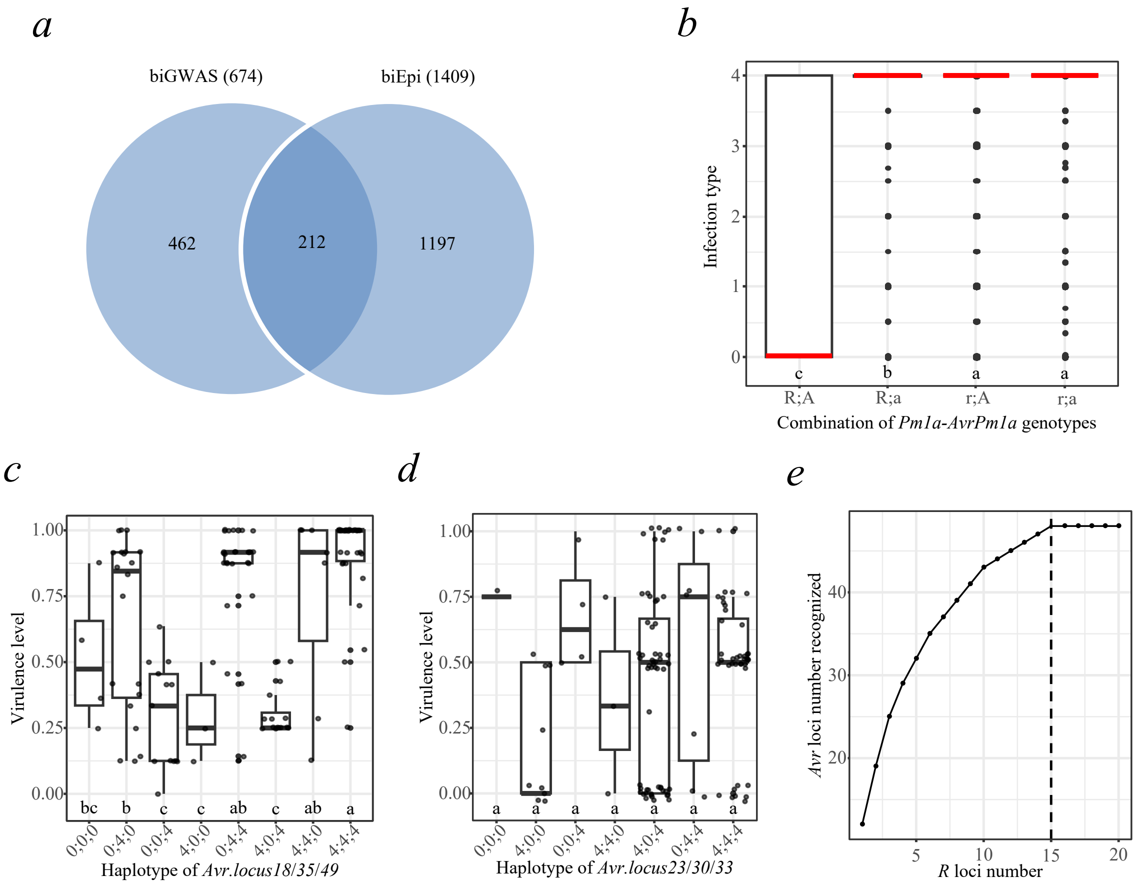


**Supplementary Figure 13. Overlaps and combinations of *R*/*Avr* loci in the biGWAS and biEpi maps.** (a) Venn diagram displaying the overlap of *R-Avr* interaction pairs identified through both biGWAS and biEpi analyses. (b) Box plot showing that the predicted *Pm1a-AvrPm1a* interacting pair results in significantly lower phenotyping scores (infection types) when *R* allele encounters the cognate *Avr* allele. 'R' represents the resistant allele of *Pm1a*; 'r' denotes the susceptible allele; 'A' denotes the avirulent allele of *AvrPm1a*, while 'a' represents the virulent allele. The red line indicates the median value. (c) Box plot depicting phenotypic differences among haplotype combinations of three *Avr* loci (*Avr.locus18/35/49*) recognized by *Pm1a*. Only wheat lines with *Pm1a* (*R* allele) were included in virulence calculation. '0' represents the avirulent allele, and '4' represents the virulent allele. The edges of the box represent the first and third quartiles, and whiskers extend 1.5 times the interquartile range. Different letters indicate statistically significant differences ($P < 0.05$, one-way ANOVA). (d) Box plot showing phenotypic differences among haplotype combinations of three *Avr* loci (*Avr.locus23/30/33*) recognized by *Pm3*. '0' represents the avirulent allele, and '4' represents the virulent allele. The edges of the box represent the first and third quartiles, with whiskers spanning 1.5 times the interquartile range. Different letters indicate statistically significant differences ($P < 0.05$, one-way ANOVA). (e) The accumulation of *Avr* loci recognized along with the number of *R* loci included. The dotted line refers to the minimal number of *R* loci needed to reach the maximal number of *Avr* loci recognized.

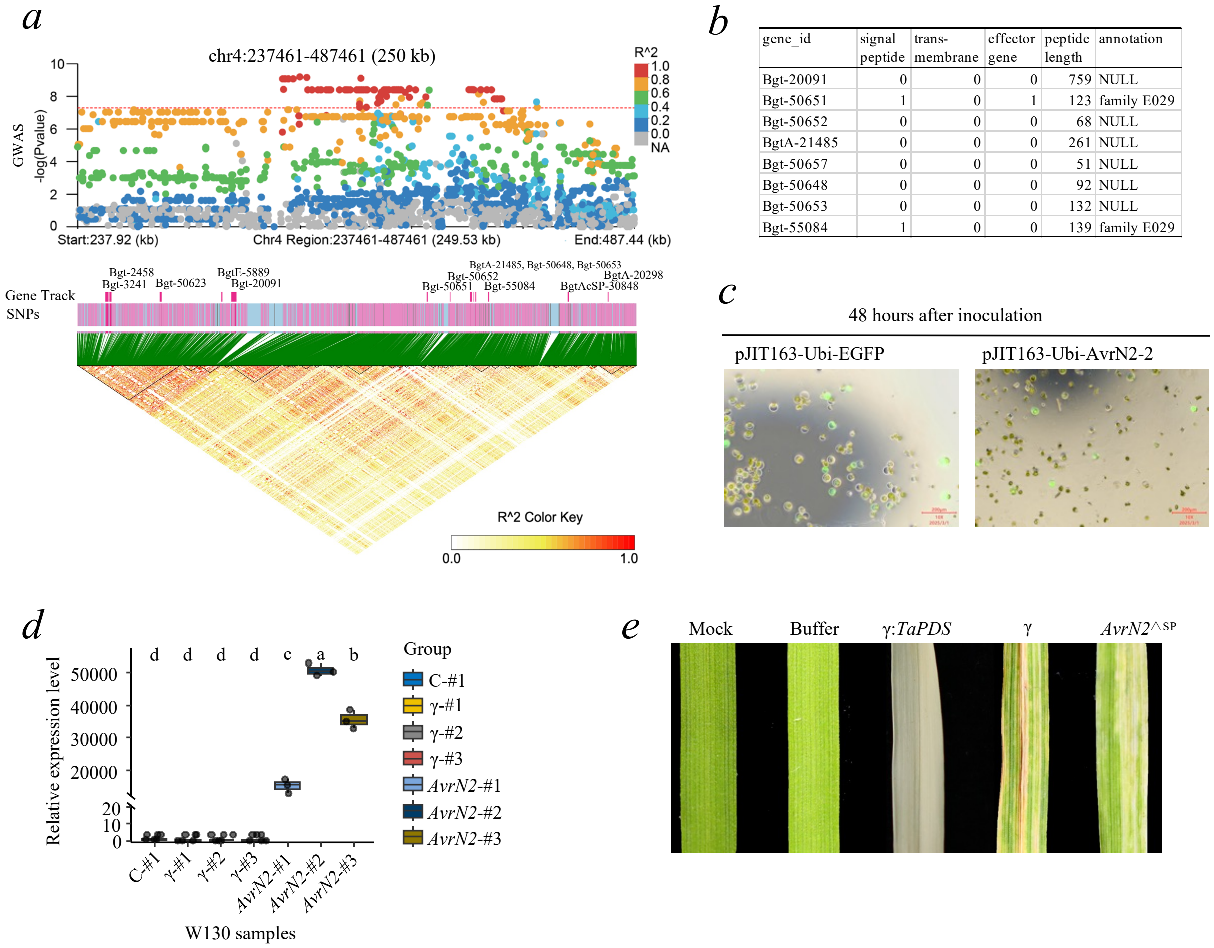


| gene_id | signal peptide | trans-membrane | effector gene | peptide length | annotation |
|---|---|---|---|---|---|
| Bgt-20091 | 0 | 0 | 0 | 759 | NULL |
| Bgt-50651 | 1 | 0 | 1 | 123 | family E029 |
| Bgt-50652 | 0 | 0 | 0 | 68 | NULL |
| BgtA-21485 | 0 | 0 | 0 | 261 | NULL |
| Bgt-50657 | 0 | 0 | 0 | 51 | NULL |
| Bgt-50648 | 0 | 0 | 0 | 92 | NULL |
| Bgt-50653 | 0 | 0 | 0 | 132 | NULL |
| Bgt-55084 | 1 | 0 | 0 | 139 | family E029 |

**Supplementary Figure 14. Association mapping and molecular validation of the *Avr.locus18*.** (a) Manhattan plot of GWAS for mapping *Avr.locus18*, showing the 250 kb genomic region surrounding the associated peak on chromosome 4. SNPs are color-coded based on their linkage disequilibrium (LD) strength with the top SNP (measured as the squared correlation coefficient, $R^2$). The accompanying LD heatmap illustrates pairwise LD ($R^2$) among SNPs in the 250 kb interval, with stronger LD represented by higher $R^2$ values (red) and weaker LD by lower $R^2$ values (white). (b) The annotation information of the genes in the mapping interval of *Avr.locus18*. '1': yes; '0': no. (c) Transient expression of *AvrN2* in W130 protoplasts was observed under fluorescence microscope after 48 h with W5 medium inoculation. Scale bar, 200 μm. (d) The relative expression level of *AvrN2* in BSMV-vox plants obtained via qRT-PCR. *AvrN2*-specific primers were used. C: control group, '#': plant ID. Three technical replicates were done for each group. The edges of the box represent the first and third quartiles, with whiskers spanning 1.5 times the interquartile range. Different letters on top of boxes indicate statistically significant differences ($P < 0.05$, one-way ANOVA). (e) BSMV-mediated expression of controls and the avirulence protein AVRN2 in leaves of susceptible wheat cv. Zhongzuo 9504. Mock: wheat plants without any treatment. Buffer: wheat plants treated with the buffer solution used for application. γ-TaPDS: wheat plants overexpressing the etiolation gene TaPDS. γ: Wheat plants overexpressing the empty vector. AvrN2$^{\Delta sp}$: Wheat plants overexpressing *AvrN2* with the signal peptide removed. Photos were taken at 10 dpi.

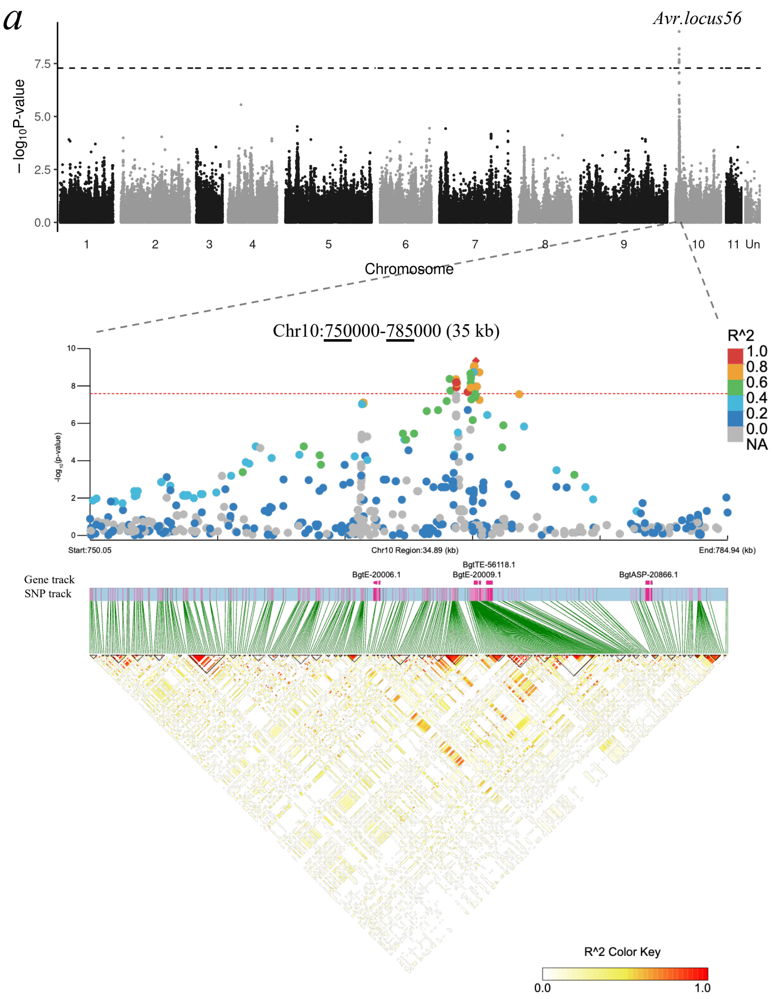


b

| gene_id | signal peptide | trans-membrane | effector gene | peptide length | annotation |
|---|---|---|---|---|---|
| BgtE-20009 | 1 | 0 | 1 | 112 | family E008 |
| BgtTE-56118 | 0 | 0 | 0 | 121 | TE gene |
| BgtASP-20866 | 1 | 0 | 0 | 109 | family E008 |

**Supplementary Figure 15. Association mapping and molecular validation of the *Avr.locus56*.** (a) Manhattan plot of GWAS for mapping *Avr.locus56*, showing the 35 kb genomic region surrounding the associated peak on chromosome 10. SNPs are color-coded by their LD strength with the top SNP ($R^2$). The corresponding LD heatmap illustrates pairwise LD ($R^2$) among SNPs in the 250 kb genomic interval with stronger LD represented by higher $R^2$ values (red) and weaker LD by lower $R^2$ values (white). (b) The annotation information of the genes in the mapping interval of *Avr.locus56*. ‘1’: yes; ‘0’: no.

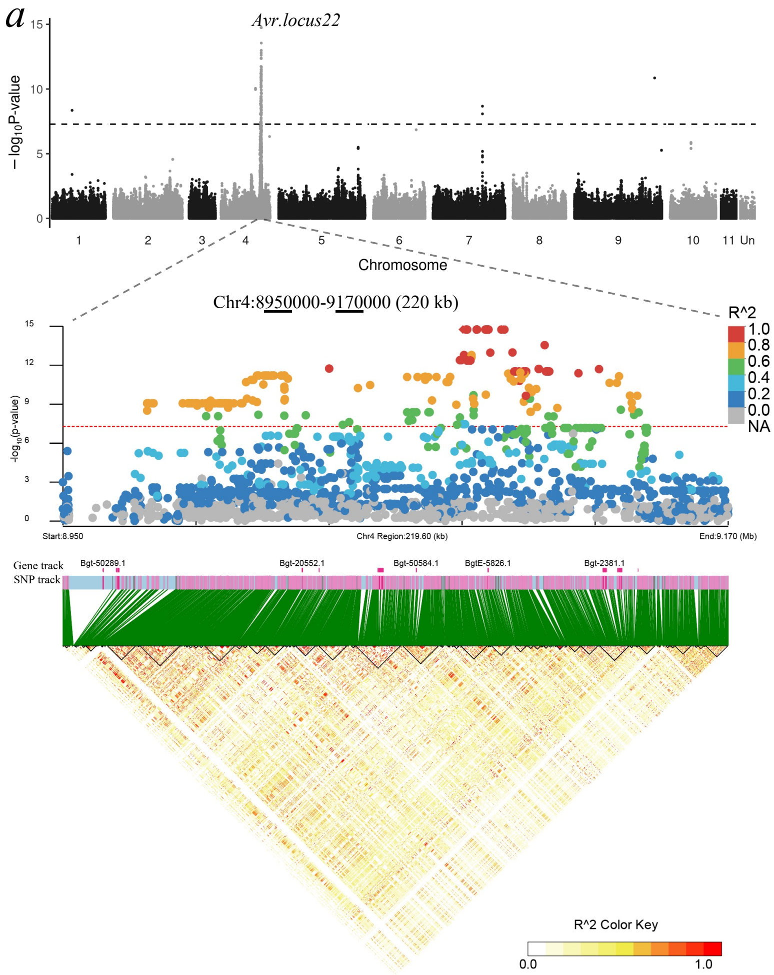


*b*

| gene_id | signal peptide | trans-membrane | effector gene | peptide length | annotation |
|---|---|---|---|---|---|
| BgtE-5826 | 1 | 0 | 1 | 110 | family E050 |
| Bgt-2381 | 0 | 0 | 0 | 447 | |

**Supplementary Figure 16. Association mapping and molecular validation of the *Avr.locus22*.** (a) Manhattan plot of GWAS for mapping *Avr.locus22*, showing the ~220 kb genomic region surrounding the associated peak on chromosome 4. SNPs are color-coded by their LD strength with the top SNP ($R^2$). The corresponding LD heatmap illustrates pairwise LD ($R^2$) among SNPs in the 220 kb genomic interval with stronger LD represented by higher $R^2$ values (red) and weaker LD by lower $R^2$ values (white). (b) The annotation information of the genes in the mapping interval of *Avr.locus22*. ‘1’: yes; ‘0’: no.

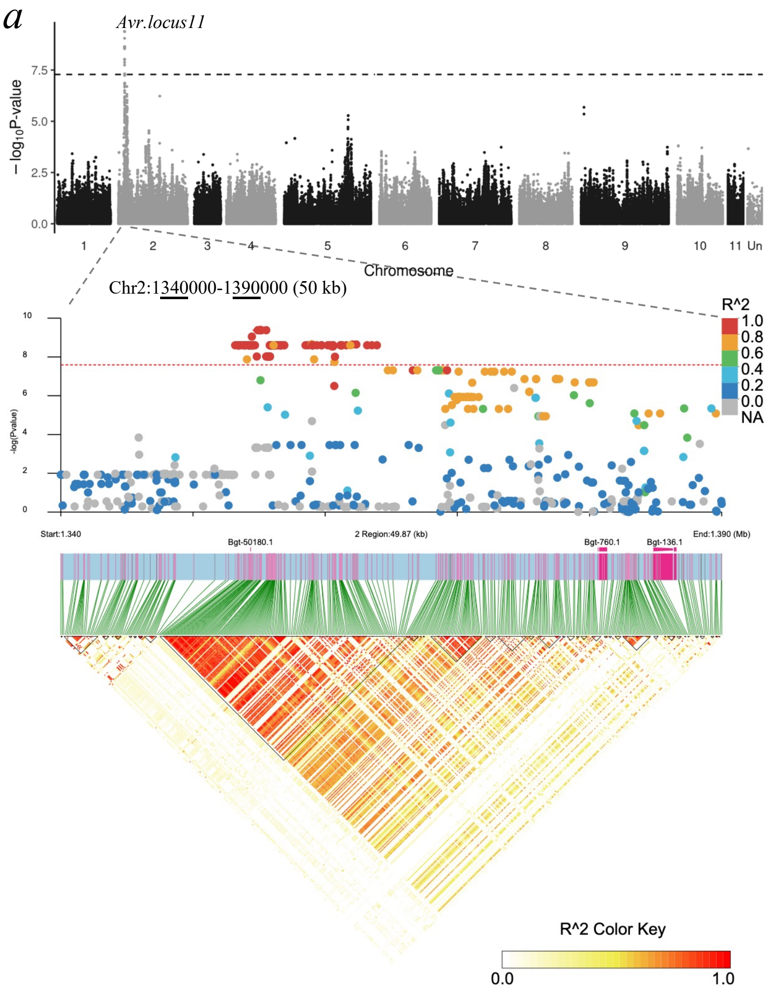


b

| gene_id | signal peptide | trans-membrane | effector gene | peptide length | annotation |
|---|---|---|---|---|---|
| Bgt-50180 | 1 | 0 | 1 | 124 | NULL |
| Bgt-760 | 0 | 0 | 0 | 216 | superoxide dismutase |

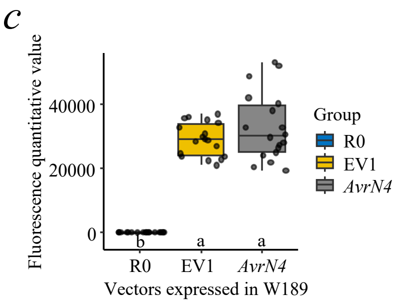


**Supplementary Figure 17. Association mapping and molecular validation of the *Avr.locus11*.** (a) Manhattan plot of GWAS for *Avr.locus11* mapping, showing the ~50 kb genomic region surrounding the associated peak on chromosome 2. SNPs are color-coded by their LD strength with the top SNP ($R^2$). The corresponding LD heatmap illustrates pairwise LD ($R^2$) among SNPs in the 50 kb genomic interval with stronger LD represented by higher $R^2$ values (red) and weaker LD by lower $R^2$ values (white). (b) The annotation information of the genes in the mapping interval of *Avr.locus11*. '1': yes; '0': no. (c) Comparison of fluorescence levels between *AvrN4* constructs in protoplasts of wheat accession W189. '*AvrN4*': over-expression vector of the *Avr* allele *AvrN4*; 'EV1': empty vector; 'R0' water control. Different letters above x-axis indicate statistically significant differences ($P < 0.05$) based on one-way ANOVA test.

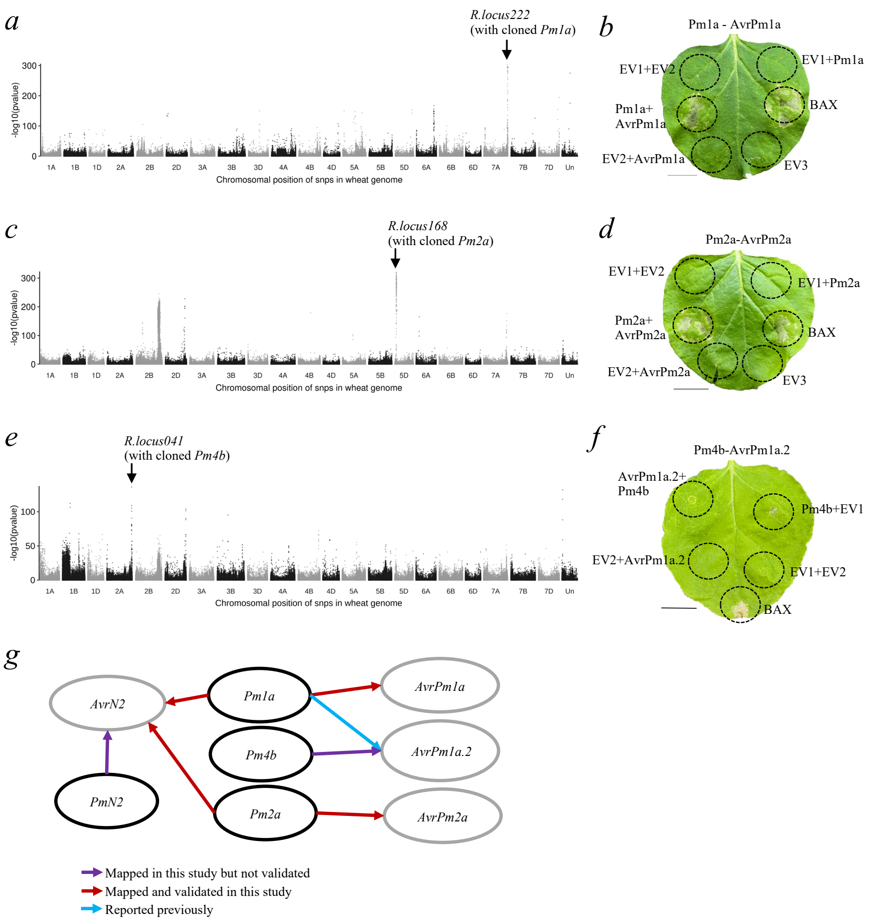


**Supplementary Figure 18. Molecular validation of the predicted *R-Avr* interacting pairs**. (a) Association levels of *AvrPm1a* with all SNPs in the wheat genome based on the epistasis map. Association p values were obtained using a general linear model. Each dot represents a single SNP. The know *Pm* genes are annotated with arrows and corresponding gene symbols. (b) Transient expression assay of the *Pm1a-AvrPm1a* interacting pair in *N. benthamiana*. The *Avr* allele *AvrPm1a* was expressed alone or co-expressed with *Pm1a* in *N. benthamiana leaves*. Empty vector (EV1/2/3) and BAX were used as controls. $OD_{600}$ = 0.5. Four-week-old *N. benthamiana* plants were used, and photos were taken at 4 dpi. Three biological replicates were performed and showed consistent results. Scale bar: 1 cm. (c) Association levels of *AvrPm2a* with all SNPs in the wheat genome based on the epistasis map. Association p values were obtained using a general linear model. Each dot represents a single SNP. The know *Pm* genes are annotated with arrows and corresponding gene symbols. (d) Transient expression assay of the *Pm2a-AvrPm2a* interacting pair in *N. benthamiana*. The *Avr* allele *AvrPm2a* was expressed alone or co-expressed with *Pm2a* in *N. benthamiana* leaves. Empty vector (EV1/2/3) and BAX were used as controls. $OD_{600}$ = 0.5. Four-week-old *N. benthamiana* plants were used, and photos were taken at 4 dpi. Three biological replicates were performed and showed consistent results. Scale bar: 1 cm. (e) Association levels of *AvrPm1a.2* with all SNPs in the wheat genome based on the epistasis map. Association p-values were obtained using a general linear model. Each dot represents a single SNP. The know *Pm* genes are annotated with arrows and corresponding gene symbols. (f) Transient expression assay of the *Pm4b-AvrPm1a.2* interacting pair in *N. benthamiana*. The *Avr* allele *AvrPm1a.2* was expressed alone or co-expressed with *Pm4b* in *N. benthamiana* leaves. Empty vector (EV1/2) and BAX were used as controls. $OD_{600}$ = 0.5. Four-week-old *N. benthamiana* plants were used, and photos were taken at 4 dpi. Three biological replicates were performed and showed consistent results. Scale bar: 1 cm. (g) Experimentally verified *R-Avr* interactions displayed as a network graph.

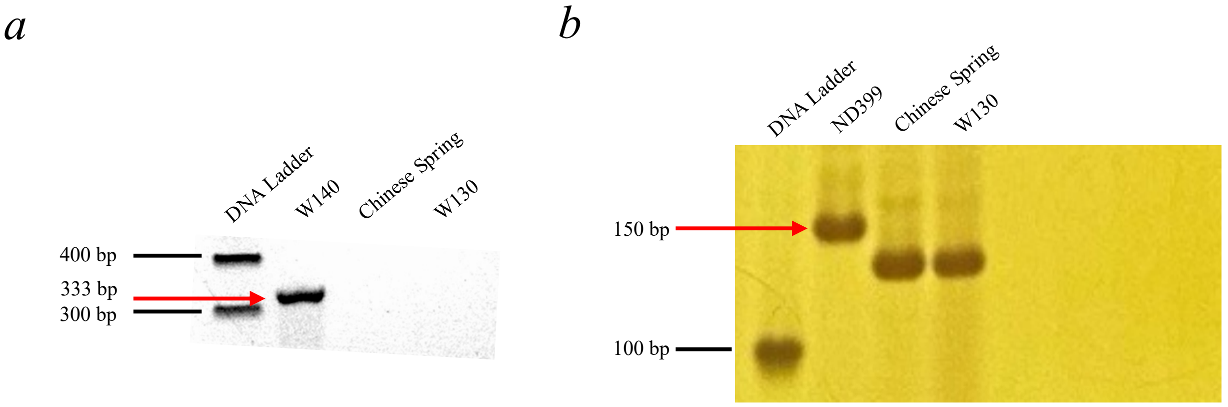


*Pm1aSTS1* amplicons    *Pm2a-map-3* amplicons

**Supplementary Figure 19. Gel electrophoresis results for PCR products of *Pm1a* and *Pm2a*.** (a) Agarose gel electrophoresis results for PCR products of *Pm1a* diagnostic marker *Pm1aSTS1*. Wheat line W140 was used as a positive control, in which the full-length cDNA sequence of *Pm1a* was amplified and confirmed by Sanger sequencing. Wheat line Chinese Spring known for its absence of *Pm1a* was used as a negative control. The target amplicon was labelled with a red arrow. (b) PAGE gel electrophoresis results for PCR products of *Pm2a* diagnostic marker *Pm2a-map-3*. Wheat line ND399 known for its presence of *Pm2a* was used as a positive control. Wheat line Chinese Spring known for its absence of *Pm2a* was used as a negative control. The target amplicon was labelled with a red arrow.

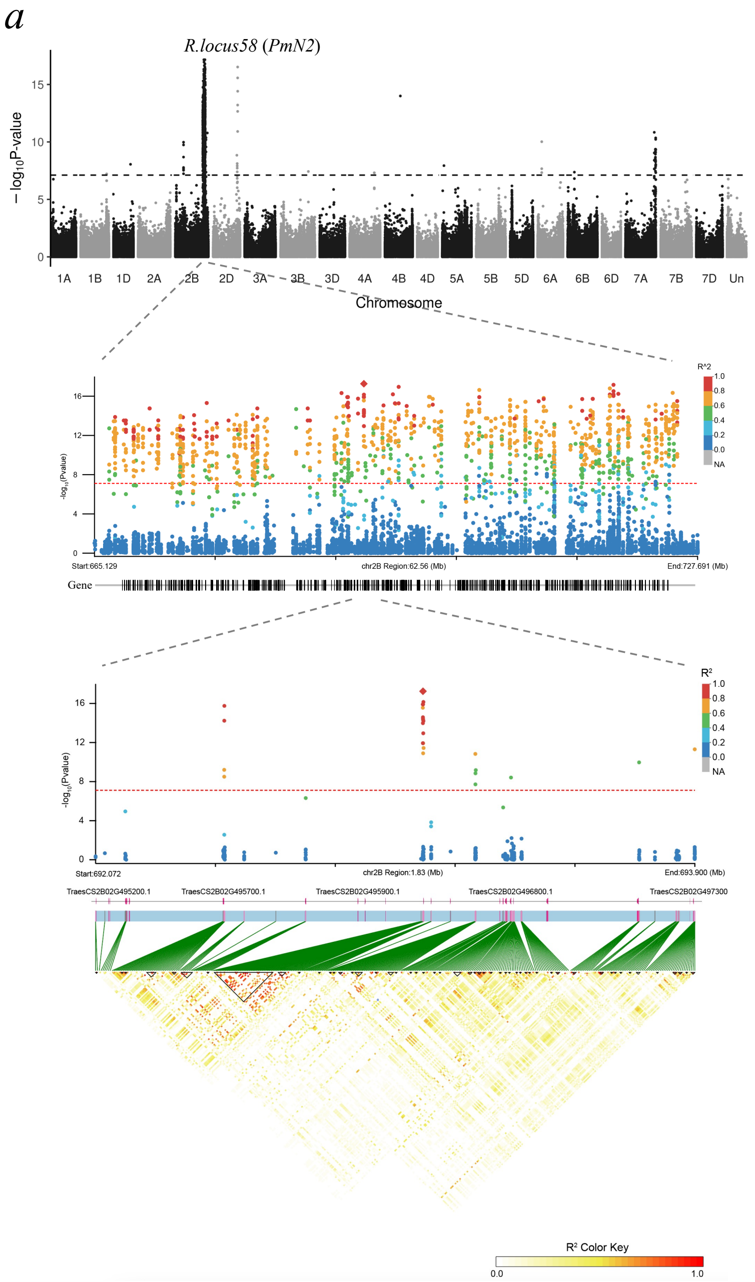


b

| Gene_ID | Chr. | Position | Annotation |
|---|---|---|---|
| TraesCS2B02G496200 | chr2B | 693071730 | BTB/POZ domain |
| TraesCS2B02G496300 | chr2B | 693095105 | BTB/POZ domain |
| TraesCS2B02G496400 | chr2B | 693153766 | BTB/POZ domain |
| TraesCS2B02G496500 | chr2B | 693304708 | BTB/POZ domain |
| TraesCS2B02G496600 | chr2B | 693313921 | Apolipoprotein D |
| TraesCS2B02G496700 | chr2B | 693321417 | NLR |
| TraesCS2B02G496800 | chr2B | 693336594 | NLR |
| TraesCS2B02G496900 | chr2B | 693346300 | NLR |
| TraesCS2B02G497000 | chr2B | 693447357 | NLR |
| TraesCS2B02G497200 | chr2B | 693723708 | NLR |
| TraesCS2B02G497300 | chr2B | 693885306 | NLR |

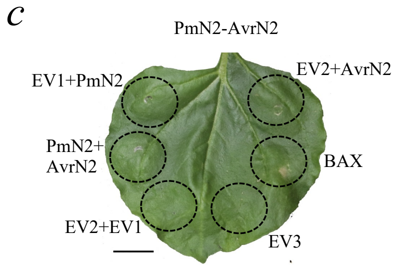


**Supplementary Figure 20. Association mapping and molecular validation of the *R.locus058*.** (a) Manhattan plot of GWAS for mapping *PmN2* in trait Bgt0199, showing the ~1.83 Mb genomic region surrounding the associated peak on chromosome 2B. SNPs are color-coded by their LD strength with the top SNP ($R^2$). The corresponding LD heatmap illustrates pairwise LD ($R^2$) among SNPs in the 1.83 kb genomic interval with stronger LD represented by higher $R^2$ values (red) and weaker LD by lower $R^2$ values (white). (b) The annotation information of the 11 candidate genes in the mapping interval of *R.locus058*. One candidate, *TraesCS2B02G496800* designated as *PmN2*, were randomly chosen for downstream functional validation. (c) Transient expression assay of the *PmN2-AvrN2* interacting pair in *N. benthamiana*. The *Avr* allele of *AvrN2* was expressed alone or co-expressed with *PmN2* in *N. benthamiana* leaves. Empty vector (EV1/2/3) and BAX were used as controls. OD 600 = 0.5. Four-week-old *N. benthamiana* plants were used, and photos were taken at 4 dpi. Three biological replicates were performed and showed consistent results. Scale bar: 1 cm.

*a*

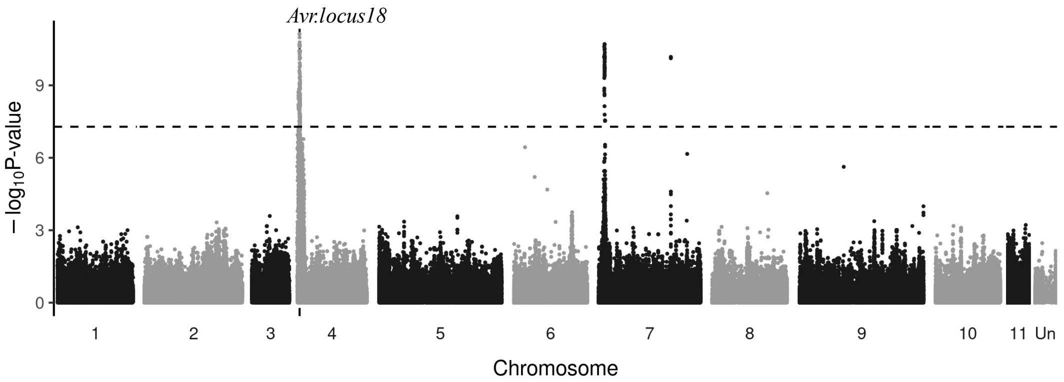


**Supplementary Figure 21. *Avr* GWAS results of the wheat cultivar Liangxing 99 carrying *Pm52*.** (a) Manhattan plot of GWAS for mapping *Avr* loci with Liangxing 99 in the *Bgt* panel. *Avr.locus18* on chromosome 4 was identified as the most significant peak.
