## Supplemental Notes for "A biGWAS strategy reveals the genetic architecture of the interaction between wheat and *Blumeria graminis* f. sp. *tritici*"

### Supplementary Notes

#### Supplementary Note 1. Detailed protocol of high-throughput phenotyping experiments.

This note outlines the detailed protocol used for high-throughput phenotyping of wheat lines resistance to *Bgt* isolates. Initially, *Bgt* isolates were inoculated onto ten-day-old, *Bgt*-free susceptible wheat seedlings (cv. Zhongzuo 9504) on a benchtop to propagate sufficient conidial spores for subsequent infections. Two days after this inoculation, seeds of the wheat lines to be tested were sown. A 276-well (23×12) tray was filled with soil, and the soil surface was moistened with an appropriate amount of water. Each well received three seeds of a single wheat line. The four corner wells were planted with the susceptible cultivar Zhongzuo 9504 as a negative control. After sowing, an elevated frame and a ventilated cover with closed ventilation ports were placed over the tray to maintain a sealed, *Bgt*-free microenvironment. The microenvironment in the tray maintained approximately 70% humidity, and the greenhouse was kept at 18°C with a 16-h light/6-h dark photoperiod. Eight days post sowing, infection was initiated on the two-leaf-stage seedlings. *Bgt* isolates propagated on susceptible wheat Zhongzuo 9504 and the seedlings of the wheat lines to be tested were transferred outdoors. Excess conidial spores of the propagated *Bgt* isolates were inoculated onto the wheat seedlings using a dusting method to ensure saturated infection. A new *Bgt*-free, ventilated cover was placed over the tray. The tray with inoculated seedlings was returned to the greenhouse to maintain the proper microenvironment. Once susceptible controls exhibited severe disease symptoms, the infection types of wheat accessions were evaluated in a separate greenhouse, usually eight days post-inoculation. The reactions to *Bgt* inoculation were examined visually with infection type (IT) recorded on a 0–4 scale: '0' for no visible symptoms; '0;' for hypersensitive reaction; '1' for disease spots less than 1 mm with a thin hypha layer and fewer spores; '2' for disease spots less than 1 mm with a thick hypha layer and more spores; '3' for scattered disease spots greater than 1 mm with a thick hypha layer and more spores; and '4' for continuous disease spots greater than 1 mm with a thick hypha layer and more spores. Plants with IT of 0–2 (no disease colonies or spots less than 1 mm) were classified as resistant, and those with IT of 3–4 (scattered spots greater than 1 mm) as susceptible. These procedures enabled a weekly experimental schedule, generating 4–6 sets of phenotyping data of the wheat diversity panel infected with *Bgt* isolates. Details are as follows:

1. Monday: Inoculate and propagate the *Bgt* isolates to obtain sufficient conidia for subsequent experiments.
2. Wednesday: Sow the wheat population for phenotyping, ensuring the number of sowing replicates corresponds to the number of *Bgt* isolates propagated on the preceding Monday.
3. Thursday: Inoculate the wheat seedlings sown on the preceding Wednesday with the prepared *Bgt* isolates.
4. Friday: Evaluate the infection types of the wheat accessions inoculated on the preceding Thursday.

**Supplementary Note 2. Assignment of previously cloned *Pm*/*AvrPm* genes to the GWAS *R*/*Avr* loci.**

This supplementary note details the process of assigning previously cloned *R*/*Avr* genes to *R*/*Avr* loci identified in this study. The assignment was conducted in three steps: determining gene positions, locating *R*/*Avr* loci, and assigning genes based on physical distance.

1. Determining gene positions:

The representative physical positions of the cloned *R*/*Avr* genes were determined in reference genomes (Chinese Spring assembly RefSeq v1.0 for wheat and isolate 96224 assembly v3.16 for *Bgt*) based on gene collinearity.

For *Bgt AvrPm* genes, published CDS sequences from NCBI GenBank of the 9 cloned *Avr* genes were aligned to CDS sequences of all genes annotated in the *Bgt* isolate 96224 genome assembly v3.16 using BLASTP program. The physical position of the top BLASTP hit (sequence identity > 80%) was classified as the position of that cloned *Avr* gene. This allows us to identify the physical positions of all nine cloned *Bgt AvrPm* genes on the *Bgt* isolate 96224 genome assembly v3.16, as follows:

(1) *AvrPm1a*'s homolog *BgtE-5612* is at 4863441 bp of *Bgt* Chromosome 1 (i.e. *AvrPm1a*: *BgtE-5612*: Chr1:4863441),

(2) *AvrPm2*: *BgtE-5845*: Chr7:935269,

(3) *AvrPm3*$^{a2/f2}$: *Bgt_avrF2_7*: Chr6: 2135223,

(4) *AvrPm3*$^{b2/c2}$: *BgtE-20002*: Chr5:18862922,

(5) *AvrPm3*$^{d3}$: *BgtE-20069b*: Chr9:369328,

(6) *SvrPm3*$^{a1/f1}$: *Bgt_BCG-1*: Chr4:10109686,

(7) *AvrPm17*: *Bgt-51729*: Chr1:4370469,

(8) *AvrPm8*: *Bgt-50847*: Chr11:2661079,

(9) *AvrPm1a.2*: *Bgt-51526*: Chr8:10726079.

For wheat Pm genes, *Pm2/3/4/5/24/38/46*, derived from bread wheat, had their physical positions determined based on reported homologs in the Chinese Spring genome. *Pm8/17/13/12/21/WTK4*, derived from wheat relatives, had their physical positions determined based on orthologs found in

Chinese Spring via BLASTP search with the top hit identified as the ortholog. The homologs or orthologs of *Pm1/41/36/60* are absent in Chinese Spring. The physical positions of *Pm1/41/36/60* were transferred from homologous/orthologous regions in Chinese Spring, identified through BLASTP of flanking genes. This allows us to identify the physical positions of all 16 wheat *Pm* genes, as follows:

(1) *Pm1*: *TraesCS7A02G553500*-3800: chr7A: 726422641,
(2) *Pm2*: *TraesCS5D02G044600*: chr5D: 43405783,
(3) *Pm3*: *TraesCS1A02G008100*: chr1A: 4497200,
(4) *Pm4*: *TraesCS2A02G558300*: chr2A: 761856303,
(5) *Pm5*: *TraesCS7B02G441700*: chr7B: 706811897,
(6/7) *Pm8/17*: *TraesCS1B02G014800*: chr1B: 7252011,
(8) *Pm13*: *TraesCS3B02G013500LC*: chr3B: 5792601,
(9) *Pm21/12*: *TraesCS6A02G120000*: chr6A: 91868733,
(10) *Pm24*: *TraesCS1D02G058900*: chr1D: 38780587,
(11) *Pm36*: *TraesCS5B02G361900*-2000: chr5B: 541342032,
(12) *Pm38*: *TraesCS7D02G080300*: chr7D: 47412062,
(13) *Pm41*: *TraesCS3B02G535900-6000*: chr3B: 775831156,
(14) *Pm46*: *TraesCS4D02G243100*: chr4D: 405770757,
(15) *Pm60*: *TraesCS7A02G553900-4200*: chr7A: 726604240,
(16) *WTK4*: *TraesCS7D02G011700*: chr7D: 5172851.

2. Locating all *R*/*Avr* loci:

The representative physical positions for the 251 *R* loci and 65 *Avr* loci were determined from the positions of representative SNPs for each locus (Supplementary Table S3 & S7).

3. Assigning the cloned *R*/*Avr* genes to *R*/*Avr* loci based on physical distance:

In the subsequent step, *Pm* or *AvrPm* genes located within the surrounding region (50 Mb for wheat genes, 50 kb for *Bgt* genes) of *R* and *Avr* loci were assigned to the nearest locus. The following assignments were made:

Nine cloned *Avr* genes to nine *Avr* loci.

(1) *AvrPm17* to *Avr.locus04* (i.e. *AvrPm17*:Avr04),
(2) *SvrPm3*$^{a1/f1}$: Avr23,
(3) *AvrPm3*$^{b2/c2}$: Avr30,
(4) *AvrPm3*$^{a2/f2}$: Avr33,
(5) *AvrPm1a*: Avr35,
(6) *AvrPm2a*: Avr38,
(7) *AvrPm1a.2*: Avr49,

(8) *AvrPm3d3*: Avr50,

(9) *AvrPm8*: Avr64,

Eight cloned *R* genes to eight *R* loci.

(1) *Pm1* to R.locus222 (i.e. *Pm1a*:R222),

(2) *Pm2*: R168,

(3) *Pm3*: R001,

(4) *Pm4*: R041,

(5) *Pm5*: R232,

(6) *Pm8*: R005,

(7) *Pm13*: R096,

(8) *Pm36*: R154.

**Supplementary Note 3. The bi-directional GWAS pipeline.**

The pipeline, utilizing genotypic data from all tested *Bgt* isolates and wheat lines, along with their interaction phenotypic data, comprises seven steps:

(1) GWAS: Perform GWAS using four statistical models (MLM, MLMM, FarmCPU, and Blink) in GAPIT for all phenotypes, encompassing both wheat resistance and *Bgt* virulence traits.

(2) Peak identification: Identify common significant peaks across the four models' GWAS results for each trait and extract the representative SNPs for each common peak.

(3) Locus merging: Merge peaks into unique loci. For *Avr* peaks, pairwise linkage disequilibrium (LD) values of representative SNPs were calculated. Peaks were merged if their most significant SNPs ( Bonferroni cutoff: p-value $\leq 0.05$/SNP number) exhibited an LD ($R^2$) of less than 0.8. The most significant SNP was then selected to represent the merged locus. This iterative process yielded the final loci. For *R* peaks, due to lower SNP density, both LD and physical distance were considered for merging. Two peak clusters were merged if their lead SNPs met any of the following criteria: a) The SNPs are separated by less than 0.25 Mb with LD $\geq 0.2$; b) Separated by less than 0.5 Mb with LD $\geq 0.35$; c) Separated by less than 1.0 Mb with LD $\geq 0.5$; d) Separated by less than 2.5 Mb with LD $\geq 0.7$.

(4) Haplotype classification: Classify resistant (avirulent) and susceptible (virulent) haplotypes for each locus in each trait.

(5) Forward DEL analysis: Perform forward DEL analysis. To identify potential *R* loci recognizing a specific *Avr* locus, isolates were divided into avirulent and virulent haplotype groups based on their *Avr* locus haplotypes. For each *R* locus, occurrence and absence frequencies were determined in both groups. In the avirulent group, the locus presence frequency was represented as AP (A for

avirulent group; P for presence) and AA (A for avirulent group; A for absence). In the virulent group, the locus presence frequency was represented as VP (V for virulent group; P for presence) and VA (V for virulent group; A for absence). These frequencies were used to construct a contingency table, and a Fisher exact test was performed. Significant p-values (p-value < 0.05) indicated potential *Avr-R* interactions, with the *R* locus frequency difference between the two groups serving as the *Avr-R* interaction weight.

(6) Reverse DEL analysis: Perform reverse DEL analysis. To identify *Avr* loci recognized by an *R* locus, wheat lines were classified into resistant and susceptible haplotype groups. For each *Avr* locus, occurrence and absence frequencies were determined. In the resistant group, the *Avr* locus presence frequency was represented as RP (R for resistant group; P for presence) and RA (R for resistant group; A for absence). In the susceptible group, the *Avr* locus presence frequency was represented as SP (S for susceptible group; P for presence) and SA (S for susceptible group; A for absence). These frequencies were used to construct a contingency table, and a Fisher exact test was performed. Significant p-values (p-value < 0.05) indicated potential *R-Avr* interactions, with the *Avr* locus frequency difference serving as the *R-Avr* interaction weight.

(7) Interaction pair integration: Integration of *Avr-R* recognition pairs from forward and reverse DEL analyses was performed. Pairs identified in both analyses were considered highly credible; those identified in only one analysis were considered potential *R-Avr* interaction pairs.

**Supplementary Note 4. Detailed protocol of wheat protoplasts system.** The protocol includes the following steps:

1. Vector construction

The cloned CDS sequences of *Bgt Avr* candidates were optimized for expression in wheat by company BGI (China, https://www.bgi.com), and subsequently the optimized CDS sequences were inserted into the unary vector pJIT163-Ubi-EGFP, which contains the NOS terminator. Additionally, the unary vector pA-35S-LUC was used as a reporter vector.

2. Cultivation of wheat seedlings

Wheat seeds were sown in a growth chamber and cultured at 23°C under a light intensity of 1000 lux. The photoperiod was maintained at 14–16 hours per day. The cultivation period lasted approximately 1–2 weeks.

3. Protoplast extraction

1) Preparation of enzyme solutions

a. Enzyme solution(40ml)

| | 40ml |
|---|---|

| | |
|---|---|
| MES (40mM) | 10ml |
| BSA | 0.04g |
| C-R10 | 0.28g |
| M-R10 | 0.14g |
| M mannitol | 4.372128g |

b. W5 solution (500ml)

| | 500ml |
|---|---|
| 0.1%glucose | 0.5g |
| 0.08%KCl | 0.4g |
| 0.9%NaCl | 4.5g |
| 1.84%CaCl2.2H2O | 9.2g |
| MES-KOH(40mM) | 25ml |

c. MMG solution（10ml）

| | |
|---|---|
| DDW | Add to10ml |
| MgCl2 (1M) | 0.15ml |
| MES(200mM） | 0.2ml |
| Mannitol（0.8M） | 5ml |

d. PEG solution（4ml）

| | |
|---|---|
| DDW | Add to 4ml |
| Cacl2（1M)） | 0.4ml |
| Mannitol（0.8M） | 1ml |
| PEG4000 | 1.6g |

2) Tender wheat leaves were cut into strips of 0.5–1 mm and immersed in a 0.6 M mannitol solution for 10 minutes in the dark. The leaves were then filtered through a mesh and placed into 50 ml of enzyme solution for digestion for 5 hours.

3) To stop enzymatic digestion, 10 ml of W5 solution was added. The enzyme solution was then filtered through a 75-μm nylon mesh into a round-bottom centrifuge tube.

4) The filtered solution was centrifuged at 100g with acceleration and deceleration both set to 3 at 23°C. The supernatant was discarded.

5) The precipitated cells were gently resuspended in 10 ml of W5 solution and placed on ice for 30 minutes.

6) The protoplasts were suspended in an appropriate volume of MMG solution and prepared for transformation.

4. Transformation of wheat protoplasts.

1) A total of 20 μg of plasmid DNA was added to a 2-ml centrifuge tube, followed by 200 μl of protoplasts. The mixture was gently mixed and left to stand for 5 minutes. Subsequently, 250 μl of PEG 4000 was added, and the mixture was gently flicked to mix. Transformation was conducted in the dark for 30 minutes.

2) To terminate the transformation, 900 μl of W5 solution was added at room temperature and inverted to mix. The transformation product was then centrifuged at 100g with acceleration and deceleration both set to 3 for 3 minutes.

3) The precipitated cells were resuspended in 1 ml of W5 solution and incubated at 23°C for 48 hours.

5. Quantitative Measurement

1) After transformation, the cells were cultured at 23± 2°C for 48 hours and then collected by centrifugation at 100g for 3 minutes, with acceleration and deceleration both set to 3.

2) The precipitated cells were lysed with 1× cell lysis buffer (Luciferase Assay System, Promega, E1501, USA) and briefly centrifuged at 12,000 rpm for 15 seconds.

3) A total of 20 μl of the supernatant and 100 μl of LUC luciferase reagent (Luciferase Assay System, Promega, E1501, USA) were added to a 96-well plate. The LUC signal was detected using a SpectraMax iD3 reader. Each sample contained three biological replicates.

**Supplementary Note 5. Detailed protocol of BSMV experiments**.

To conduct transient effector overexpression in wheat using binary Barley stripe mosaic virus (BSMV) derived vectors, the BSMV-vox system [1] was used. The plasmid carrier of BSMV-vox system was provided by Dr. Wei Wang from Nanjing Agricultural University. All plants were grown in a round pot with a diameter of 25 centimeters which filled with the growing mix (moss peat: flower nutrient soil: perlite is 1:1:1). All the plants were grown in a growth chamber at 24 °C day/20 °C night conditions with 16 h supplemental light. The BSMV-vox system includes three binary vector plasmids, pCaBS-α, pCaBS-β and pCaBS-γ. Binary vectors pCaBS- α and pCaBS-β [2] directing expression of the BSMV genomic RNAs α and β, respectively. The vector pCaBS-γP2A-AvrN2 directing expression of the modified RNA γ that is used for insertion of foreign sequences targeted for *in planta* expression. The self-cleaving peptide of P2A (ATNFSLLKQAGDVEENPGP) with SGGS linker has high cleavage efficiency, approaching 100% in some cases. The construction of pCaBS-γP2A-AvrN2 is as follows.

(1) Synthesize the P2A cleavage peptide by BGI (Beijing).

(2) Amplify the nucleotide sequences of P2A and AvrN2 (wheat-optimized sequence) using Tks Gflex™ DNA Polymerase (R060).

(3) Use restriction enzyme ApaI (R0114) to perform a single-enzyme digestion of the BSMV-γ vector, and then purify the linearized vector using gel electrophoresis.

(4) Purify the PCR production and the linearized vector by FastPure Gel DNA Extraction Mini Kit (DC301).

(5) Recombine linearized vector and PCR production using pEASY®-Basic Seamless Cloning and Assembly Kit (CU201).

(6) Pick single colonies performing the colony PCR and sequence.

(7) Sequence the correct samples and transform them into *Agrobacterium* to conduct functional validation.

The method was adapted from Dr. Yanpeng Wang's previous paper [3]. The specific operation is as follows:

1. BSMV-sg Vector Inoculation in *Nicotiana benthamiana*.

(1) Transform the plasmids pCaBs-BSMVα, pCaBs-BSMVβ, and pCaBS-γP2A-AvrN2 into the *Agrobacterium* strain GV3101 psoup-p19. Incubate at 28 °C for 48 hours. Select single colonies and screen for positive clones using colony PCR.

(2) The following primer combinations can be used for colony PCR: BS8 + BS32 (α-chain); BS9 + BS32 (β-chain); BS11 + BS32 (γ-chain). Primer sequences: BS8: CAACTGCCGATGATCTGTCGTGTAG, BS9: CCGACGCGGAAATTCGTCAAGC, BS11: GGTAGAACTGATGTGAGAGATGTAGAAG, BS32: TGGTCTTCCCTTGGGGGACCGAA.

(3) Select positive single colonies and inoculate them into an appropriate volume of liquid LB medium containing rifampicin (25 mg/L) and kanamycin (100 mg/L). Incubate at 28°C for 12–16 hours. Centrifuge the *Agrobacterium* cultures at 4,000 rpm for 10 minutes, discard the supernatant, and collect the pellet. Resuspend the pellet in *Agrobacterium* suspension buffer (10 mM MES, 10 mM $MgCl_2$, 150 μM Acetosyringone, pH 5.8) and incubate at room temperature for at least 2 hours. For single-gene overexpression, mix *Agrobacterium* suspensions containing pCaBs-BSMVα, pCaBs-BSMVβ, and pCaBS-γP2A-AvrN2 to achieve a final $OD_{600}$ of 0.6 for each. Inject the mixture into leaves of 4- to 6-week-old *Nicotiana benthamiana* plants using a needleless syringe.

2. Transferring Crushed Leaf Sap from *Nicotiana benthamiana* to wheat.

(1) Collect the inoculated *Nicotiana benthamiana* leaves 3–7 days post-inoculation.

(2) Grind the leaves with transfer buffer (10 mM sodium phosphate buffer, pH 5.8, supplemented with freshly prepared 0.5% (w/v) sodium sulfite) at a ratio of 1:5 (w/v).

(3) Depending on the plant's growth stage, each *Nicotiana benthamiana* plant can accommodate 3–4 leaves for injection.

(4) To inoculate 10 wheat plants (including all tillers), approximately 4–6 *Nicotiana benthamiana* plants are needed, depending on the leaf size and the number of wheat tillers.

(5) Once the leaves are finely ground without large debris, wear latex gloves and use your thumb and index finger to dip into the sap for transferring.

(6) Transfer the sap to wheat leaves at the jointing stage. For each tiller, transfer the sap to the two most recently emerged leaves. Rub the sap from the leaf base to the tip while stabilizing the leaf with the other hand to prevent detachment. Each leaf should be rubbed 6–10 times in one direction. Change gloves when transferring different constructs.

(7) Approximately 7 days post-transfer, BSMV infection symptoms can be observed on systemic leaves that were not inoculated.

**Supplementary Note 6. Detailed protocol of transient expression in *Nicotiana benthamiana*.** The protocol includes the following steps:

1. Plant materials.

Wheat cultivars were grown in a greenhouse under a 14-hour light/10-hour dark photoperiod at 23°C/18°C with 70% relative humidity. *Bgt* isolates were maintained on excised wheat leaves, which were cultivated in culture dishes at 18°C and 65% relative humidity under a 16-hour light/8-hour dark photoperiod with a light intensity of 130 μE $m^{-2}$ $s^{-1}$. *Nicotiana benthamiana* was used for transient gene expression analysis. Seedlings were grown in a controlled growth chamber at 25°C and 65% relative humidity under a 16-hour light/8-hour dark photoperiod with a light intensity of 130 μE $m^{-2}$ $s^{-1}$.

2. RNA extraction and candidate gene cloning.

Total RNA was extracted and purified using the FastPure Universal Plant Total RNA Isolation Kit (RC411, Vazyme, Beijing, China). cDNA synthesis was performed using the PrimeScript™ RT Reagent Kit with gDNA Eraser (RR047, TAKARA, Beijing, China). All primers used in this study were designed using NCBI Primer-BLAST (https://www.ncbi.nlm.nih.gov/tools/primer-blast/). Cloning of candidate R and *Avr* genes was performed using the Tks Gflex™ DNA Polymerase Low DNA Kit (R091, TAKARA, Beijing, China). The CDS sequence of the R or *Avr* candidate gene was cloned by PCR (see methods). Afterwards, signal peptides in translated protein sequences of the cloned CDS sequences were predicted using the online SIGNALP algorithm (http://www.cbs.dtu.dk/services/SignalP-3.0/) and subsequently replaced with a start codon. The resulting sequences were predicted to identify mature peptide. The coding regions of the cloned sequences for mature peptides were integrated into the binary vector pCambia2300 via homologous recombination using pEASY®-Basic Seamless Cloning and Assembly Kit (CU201, TransGen Biotech, Nanjing, China). Subsequently, the recombinant plasmids were introduced into Escherichia coli DH5α cells. All constructs were verified by Sanger sequencing and transformed into *Agrobacterium* tumefaciens strain GV3101 using the freeze-thaw transformation protocol described by Weigel & Glazebrook [4].

3. Transient gene expression analysis in *Nicotiana benthamiana*.

The third and fourth leaves of one-month-old seedlings were selected as optimal tissues for infection. Candidate genes were cloned into expression vectors for expression in *Nicotiana benthamiana*. Agrobacterium-mediated expression in *N. benthamiana* and hypersensitive response (HR) assessment were performed 5-10 days after *Agrobacterium* infiltration, following the protocol described by Bourras *et al.* [5]. Three biological replicates were used for each injection.

**Supplementary Note 7. The instruction of the online bi-directional GWAS demonstration.**

To illustrate the GWAS results of this study and demonstrate the bi-directional GWAS pipeline, we built an online portal, the Wheat-Powdery Mildew Bidirectional GWAS Database, accessible at http://crop-pathogen-genomics.cn/Wheat_Bgt_biGWAS/. The following figure illustrates the schematic representation of the website.

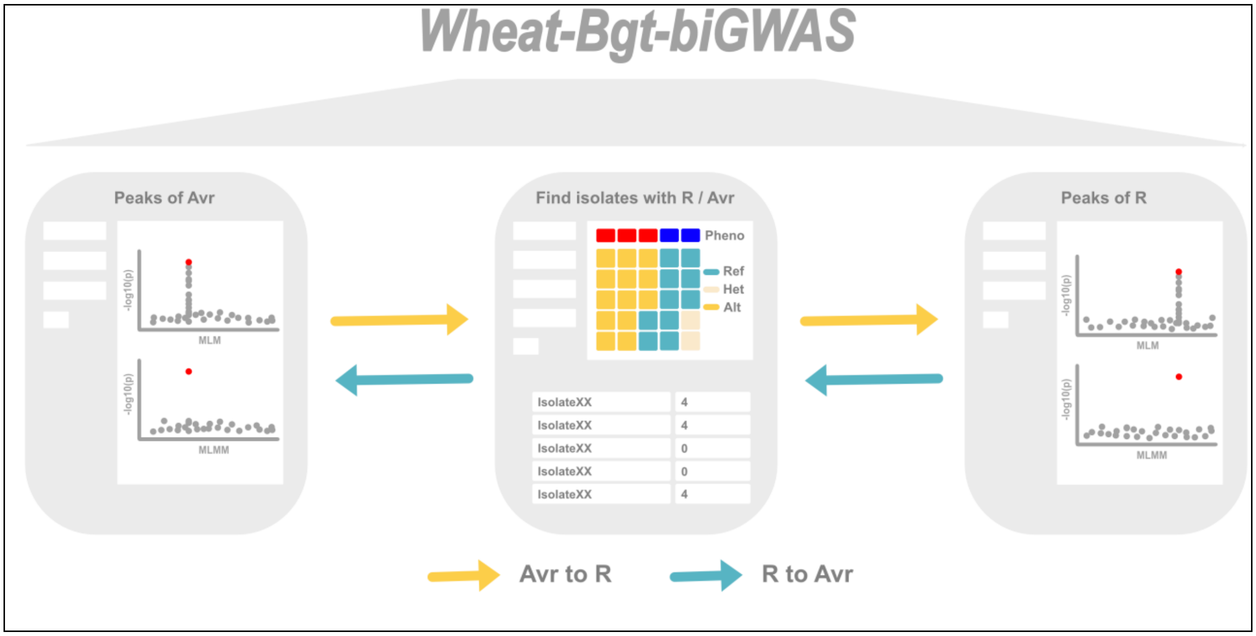


This website visualizes the bi-directional GWAS workflow and enables online retrieval of GWAS results through the integration of three panels, as illustrated in the following figure. The first panel (Peaks of *Avr*) allows users to search for specific *Avr* loci and display GWAS Manhattan plots. Clicking the red dot, which represents an *Avr* locus, will input the locus information into the search box of the second panel (Find isolates with *R*/*Avr*). The second panel displays the population haplotype heatmap for the retrieved *Avr* locus and generates a table indicating whether tested isolates carry this *Avr* allele. Subsequently, users can select an isolate from the table and input it into the search box of the third panel (Peaks of *R*). The third panel visualizes the GWAS results of *R* loci. By comparing the *R* peak distribution differences between GWAS results from carrying-*Avr* and not carrying-*Avr* isolates, the *R* loci associated with the *Avr* locus can be identified. Similarly, the reverse process, from *R* to *Avr*, can also be performed.

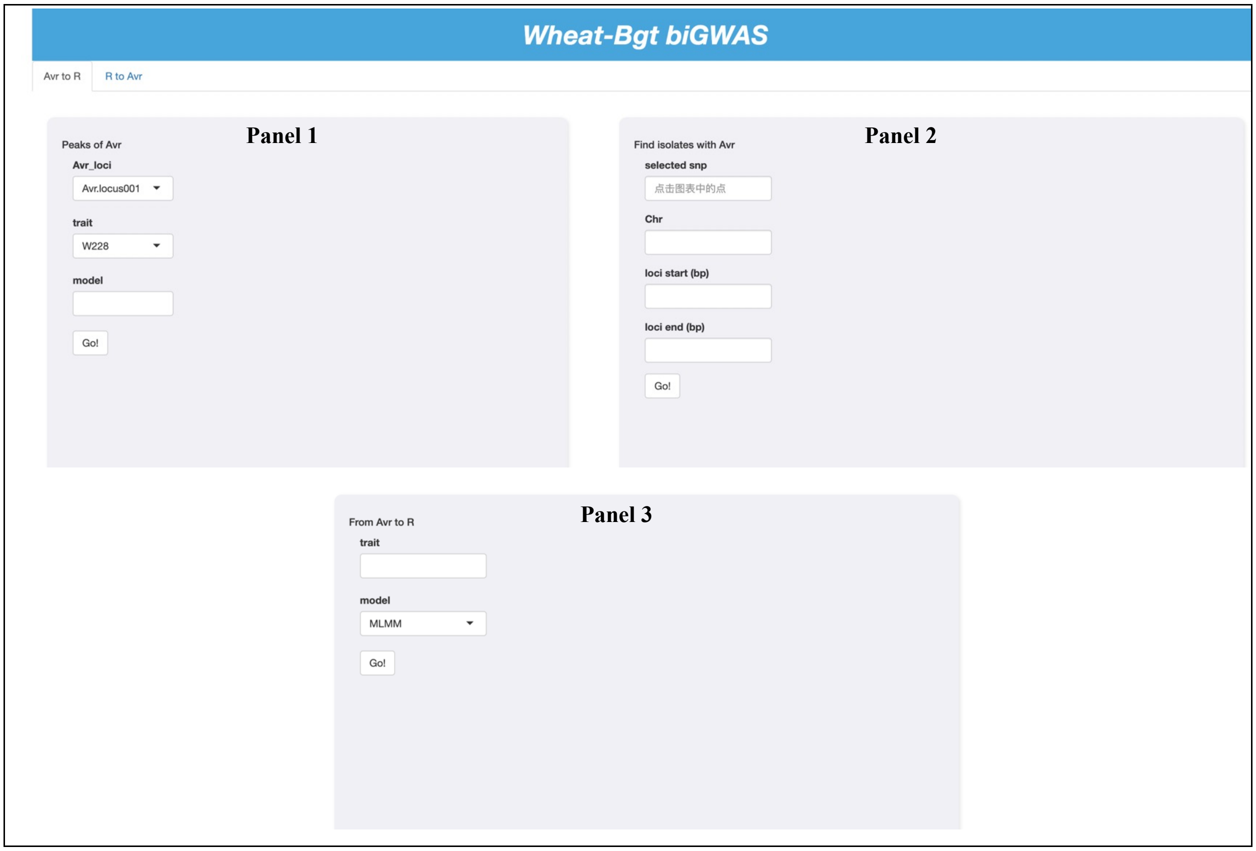


Below is the detailed usage of the website, including multiple steps, using *Avr.locus18* as an example.

**Step 1:** Selecting an *Avr* locus. In Panel 1, as illustrated in the following figure, select an *Avr* locus (e.g., *Avr.locus18*) and a wheat that locates this *Avr* locus. Choose the desired GWAS model and click "Go" to view the GWAS results.

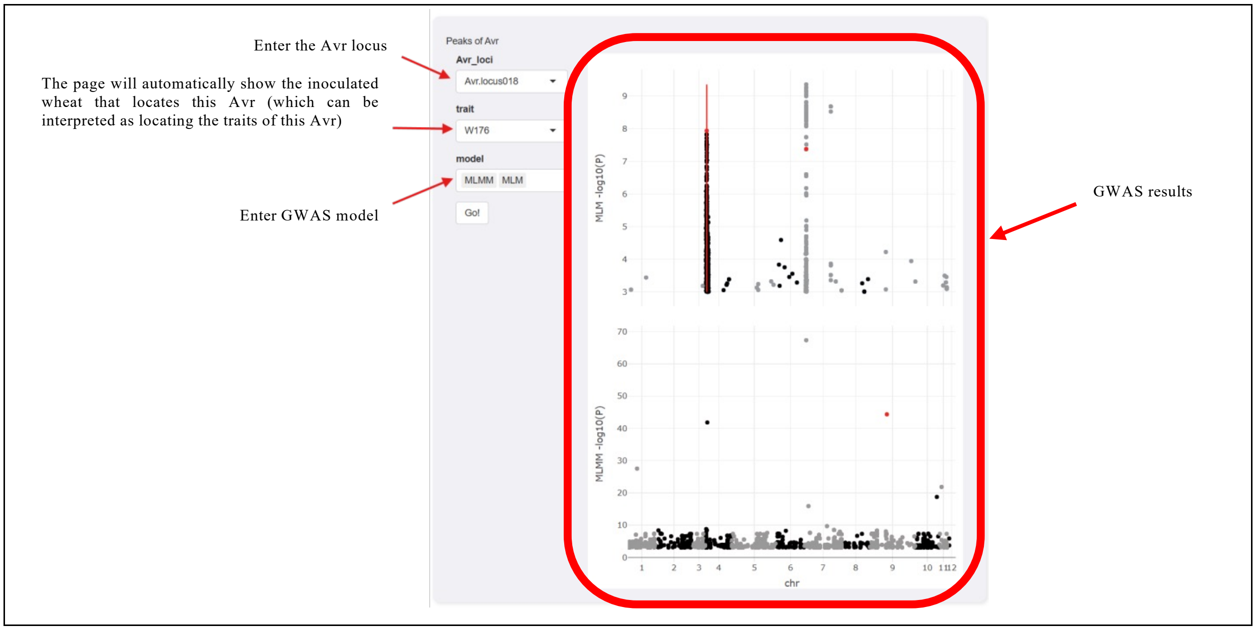


**Step 2:** Inputting the *Avr* locus information. Click the red dot representing the *Avr* locus on the Manhattan plot in Panel 1. This automatically inputs the selected *Avr* locus information into Panel 2.

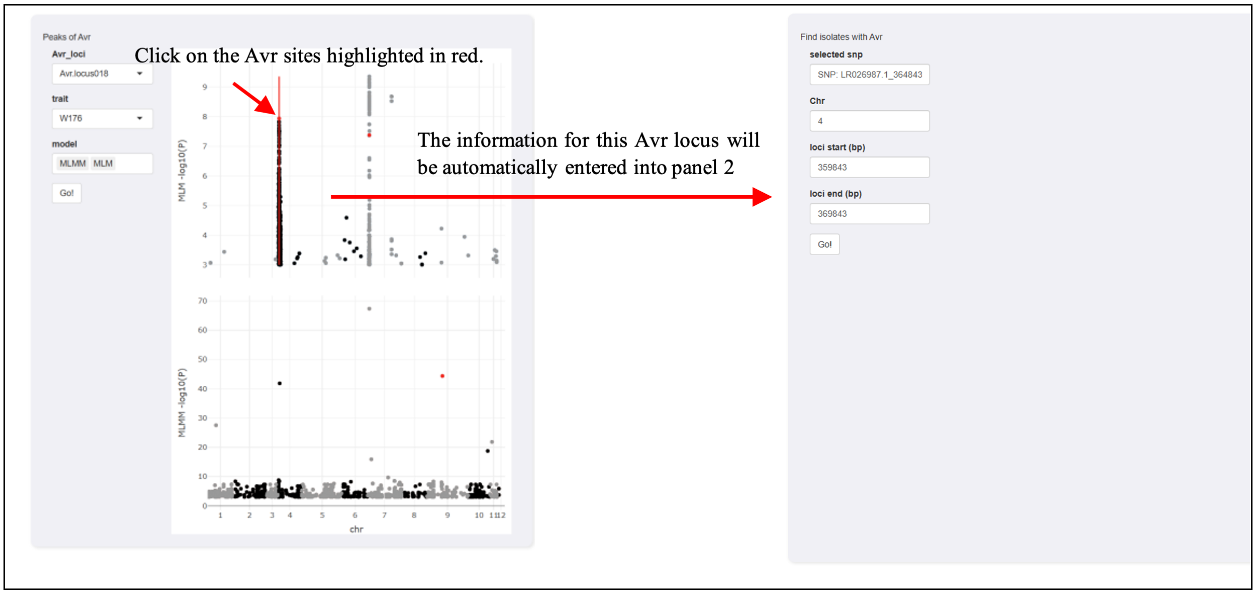


**Step 3:** Displaying the haplotype heatmap. In Panel 2, click "Go." This displays a haplotype heatmap for the inputted *Avr* locus and generates a table listing isolates that carry or do not carry this *Avr* locus.

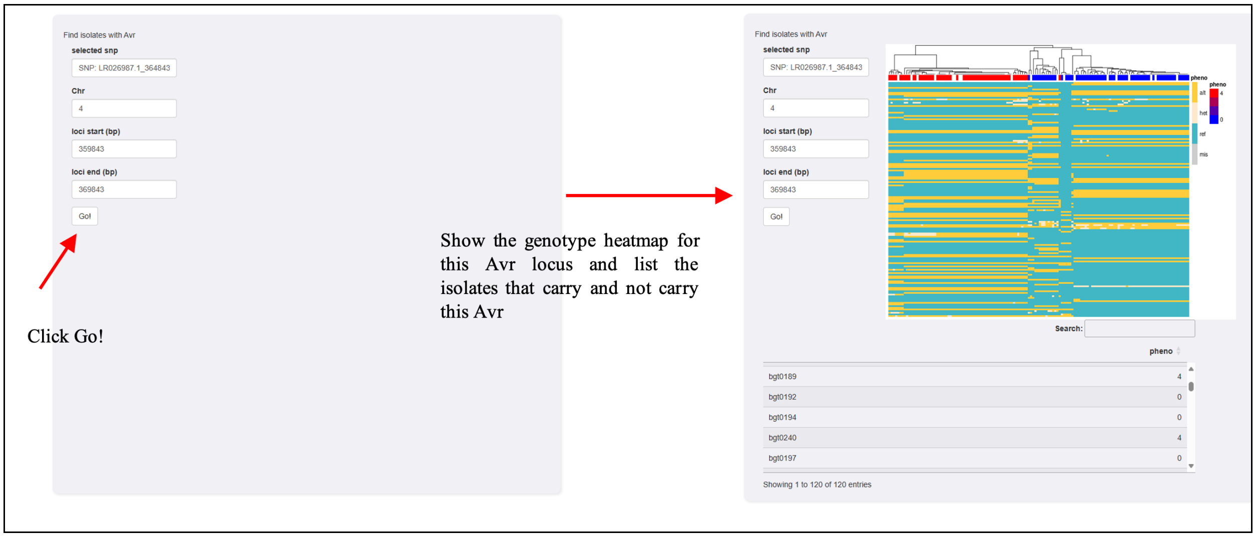


**Step 4:** Selecting isolates. Select isolates from the table in Panel 2. These selected isolates are input into Panel 3.

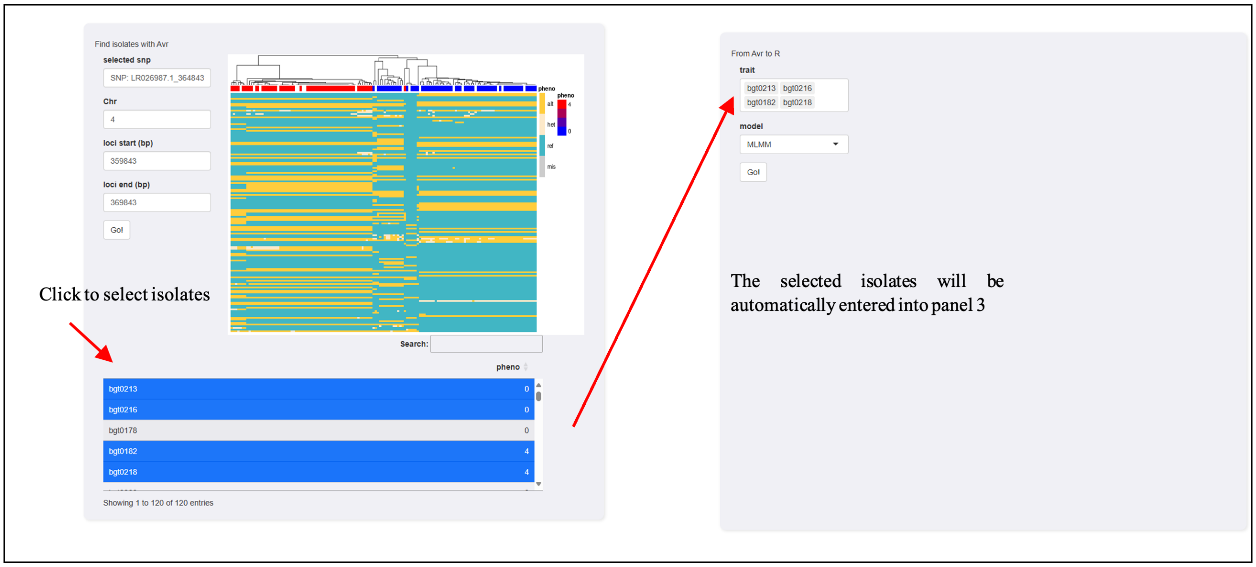


**Step 5:** Viewing *R* peak distributions. In Panel 3, click "Go." This panel displays the GWAS results of *R* peaks. By comparing the differences in peak distributions, the *R* loci associated with the *Avr* locus interaction can be predicted. For example, for *Avr.locus18*, we selected isolates bgt0213 and bgt0216, which carry this *Avr*, and bgt0218 and bgt0182, which do not. The *R* peaks identified by the four isolates' GWAS results in the wheat population were different. The isolates carrying the *Avr* locus showed a peak marked within a red box, while those without the *Avr* did not. This indicates that the *R* locus associated with this peak corresponds to the *Avr* locus, suggesting an interaction relationship.

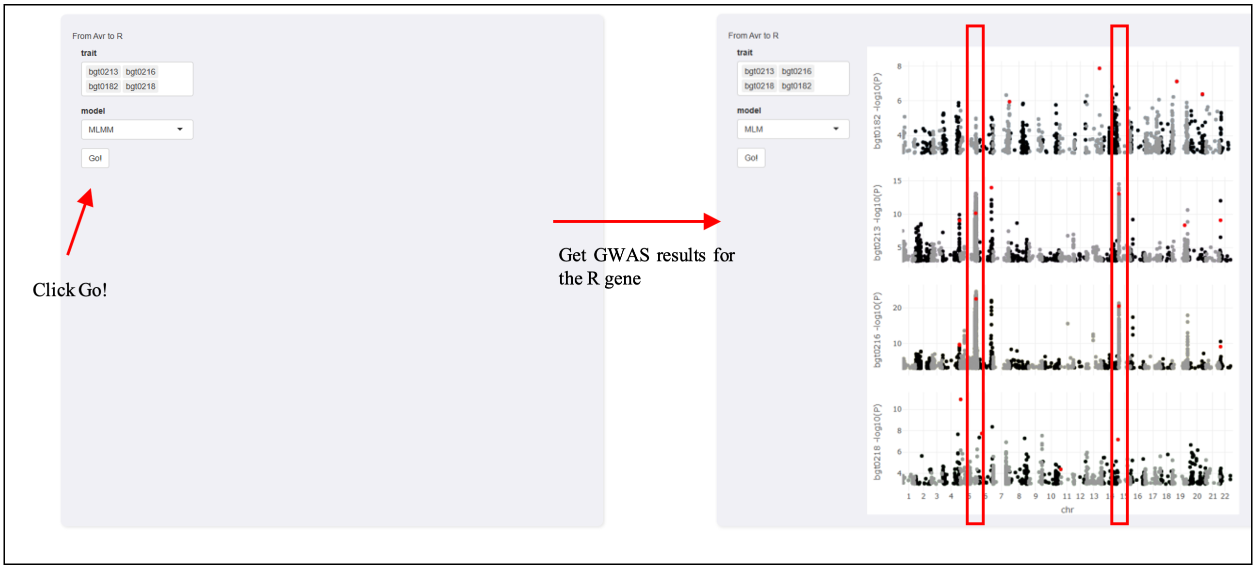


**Step 6:** Reverse Process**.** Similarly, the reverse process, *R* to *Avr*, can be performed in the same way. The website provides two separate pages with a consistent design for each workflow.

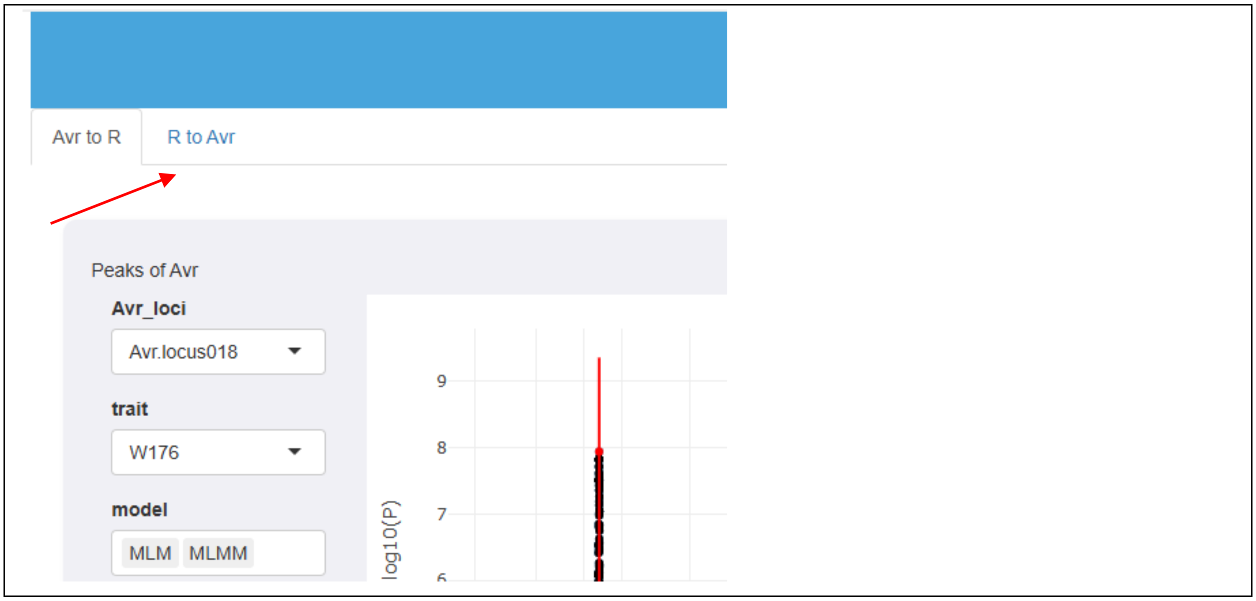


**Notice**. Please consider the following when using the website:

1. Users are strongly advised to follow the operation process outlined in the user manual to avoid inefficient searches and errors.
2. The website is hosted on a cloud server with limited computational resources. Consequently, the website's response time may be slow. Users are requested to refrain from frequent or repetitive interactions, as this may cause service interruptions.
